# Recombine and succeed: a story of Cry toxins to expand the host range

**DOI:** 10.1101/2023.12.13.571466

**Authors:** Anton E. Shikov, Ruslan O. Alagov, Anton A. Nizhnikov, Maria E. Belousova, Kirill S. Antonets

**Affiliations:** Laboratory for Proteomics of Supra Organismal Systems, All Russia Research Institute for Agricultural Microbiology (ARRIAM), 196608 St. Petersburg, Russia; Faculty of Biology, St. Petersburg State University (SPbSU), 199034 St. Petersburg, Russia

**Author notes:** Correspondence: Kirill S. Antonets.

**Keywords:** Cry toxins, *Bacillus thuringiensis*, recombination, domain swapping, host specificity, purifying selection, MGEs, insertion sequences, conservative regions

## Abstract

**Background:** Cry toxins are the most widely used sources of bioinsecticides in agriculture; therefore, improving their functionality requires a deep understanding of natural evolution. It is thought that Cry toxins emerge via domain III swapping, yet the underlying mechanism remains unclear.

**Results:** We detected 50 recombination events using a dataset of 368 clusters representing a known diversity of Cry toxins using a computational analysis. Not only do domain swaps engage all the domains, but they also occur continuously with approximately 70% of toxins participating in domain exchanges. Once they happen, hybrid toxins face purifying selection pressure reflecting the advantageous nature of receiving novel domains. When these domain exchanges occur, their host specificity changes dramatically. Strains housing these loci are enriched with *cry* genes and can kill a broader spectrum of hosts, thus implying that recombination allows them to occupy novel niches. The respective recombination-affected *cry* genes are flanked with insertions and harbor highly conservative blocks between the domains’ borders suggesting that the genomic context governs the intra-domain recombination.

**Conclusions:** Our study expands the established views of the role of recombination in the emergence of Cry toxins. Here, we demonstrate that the domain exchanges shape both Cry sequences, the composition of toxins in bacterial strains, and the sets of hosts affected. The collected data allowed us to propose a mechanism for how these toxins originate. Overall, the results suggest that domain exchanges have a profound impact on Cry toxins being a major evolutionary driver.

## Introduction

Fitting the paradigm of organic farming, biopesticides applying *Bacillus thuringiensis* (*Bt*) and its metabolites account for 90 to 97% of the biocontrol agents in pest management [1,2]. *Bt* crops have spread across the globe slowly but surely replacing chemical insecticides. By current reckonings, more than 50% of the cotton and 40% of the corn in the USA are genetically modified to carry *Bt* genes [3]. The worldwide usage of transgenic crops has risen from 32 to 191 million hectares by 2018 [4–6]. While biopesticides market, being relatively new, constitutes only about 5% of the global pesticide market, itshowcases almost 10% annual growth which is expected to bring about nearly equivalent market sizes for chemical and biological pesticides by the late 2040s-early 2050s [7,8].

*Bt* is a natural soil dweller in varied conditions and geographic areas [9]. It occupies dozens of ecological niches, e.g., invertebrates and vertebrates [10], including humans [11], plant phylloplane and rhizosphere [12], aquatic systems [13], etc., participating in an intricate system of pathogenic, commensal, and symbiotic interactions [14]. *Bt* belongs to the *B. cereus sensu lato* along with infection agents, i.e., *B. cereus sensu stricto* and *B. anthracis* [15]. Nonetheless, commonly used genetic markers and complete genomes fail to distinguish these species; thus, *Bt* should be treated as a biovar that bears genes encoding specific insecticidal toxins [16–18]. The most prominent characteristic of *Bt* is the presence of crystalline inclusions termed the parasporal body that can account for up to 25% of the dry weight of the bacterium [4]. The crystals mainly consist of δ-endotoxins comprising crystal (Cry) and cytolytic (Cyt) toxins [19]. Nevertheless, *Bt* is capable of producing a wide range of excreted moieties such as secreted insecticidal proteins (Sip), vegetable insecticidal protein (Vip), β-exotoxins, chitinases, metalloproteases, etc. [20].

Cry toxins are considered major virulence factors able to kill certain species [6,21,22]. Despite the immense diversity in amino acid sequences, all Cry proteins possess a conservative organization made up of three domains [4]. Domain I is formed by seven α-helix bundles with six amphipathic helices ensuring insertion into the cytoplasmic membrane, followed by pore formation [23]. Domain II incorporates 11 beta-sheets with antiparallel sets surrounding the hydrophobic core, constituting a β-prism structure [3]. It resembles carbohydrate-binding proteins such as lectins and performs binding to larval gut cell receptors [24]. Domain III is a β-sandwich with two antiparallel β-sheets forming a jelly roll topology [25]. Domains II and III are jointly viewed as specificity determinants [3,21]. We should note that Cry toxins enter the insects’ gut as longer protoxins with N- and C-terminal regions that are cleaved upon entering [19]. The current nomenclature of Cry toxins is built upon the pairwise identity of amino acid sequences gaining four ranks presented by Arabic numbers and letters with distinct identity thresholds [26].

A never-ending battle between hosts and pathogens provokes the development of resistance to Cry toxins. The resistance could appear through mutations in receptors [27], intensification of the immune response [3], and reduced expression of receptor-encoding genes [28]. Since the host-pathogen dialogue is a two-way street, several trends in *Bt* evolution were described. Apart from mutationing in domains II and III under positive selection pressure [29], swapping the third domain has been suggested as the primary evolutionary route allowing *Bt* to infect new hosts [30]. The hypothesis was backed by *in vitro* experiments of domain III exchange improving toxicity [31] and a comparison of Cry sequences as well as the topologies of phylogenetic trees reconstructed on the individual domains [32,33]. Phylogenetic clustering also provided insights into the spectrum of activity. For instance, predicted anti-coleopteran properties of Cry1B and Cry1I due to grouping with Cry3, Cry7, and Cry8 were later confirmed experimentally [34]. The domain-shuffling technique became a powerful tool for overcoming resistance and enhancing toxicity [35–41]. With that being said, it is quite surprising that there are few reports on natural chimeric toxins in *Bt* isolates [42–44].

Here, we conducted a large-scale bioinformatic study to estimate the frequency of domain swapping and concomitant host specificity changes focusing on the known diversity of Cry toxins. In this research, we first explicitly illustrated how prevalent domain swappings are and how profound their impact on phylogeny and host preference is. Taken together, the obtained results reveal that recombination events mostly happen during the contact of specialist populations which then entail the emergence of strains that actively recombine and maintain a broad host range. This process modifies the composition of *cry* genes harbored by the respective strains while conservative blocks between domains govern intra-genic recombination. Our results suggest domain exchanges are the key driver of Cry toxins’ evolution leading to the continuous expansion of their diversity and causing severe host specificity changes which allow *B. thuringiensis* strains to infect new hosts and/or circumvent their resistance.

## Results

### The existing data hides a variety of unexplored Cry toxins

To characterize the natural diversity of Cry toxins, we gathered sequences from four publically available databases, namely, NCBI Assembly, Genebank, IPG (Identical Protein groups), and the BPPRC (Bacterial Pesticidal Protein Resource Center) [45]. A total of 3305 sequences of 3-D cry toxins were retrieved, 941 of which remained after deduplication with a 100% identity threshold, and 368 reference clusters with a 95% identity threshold were selected afterward (fig. 1a, tab. S1). According to pairwise identities between domains, the first one was slightly more conservative (53.2%) than the second (50.2%) and the third (51.8%) (fig. S1a, tab. S2). In addition, the first and the second domains were longer (612 b.p. on average) than the third one (418 b.p.) (fig. S1b, tab. S2) and contained a higher fraction of conservative sites (fig. S1c, tab. S3). There was no correlation between domain lengths and mean identity (fig. S1d).

**Fig 1.**
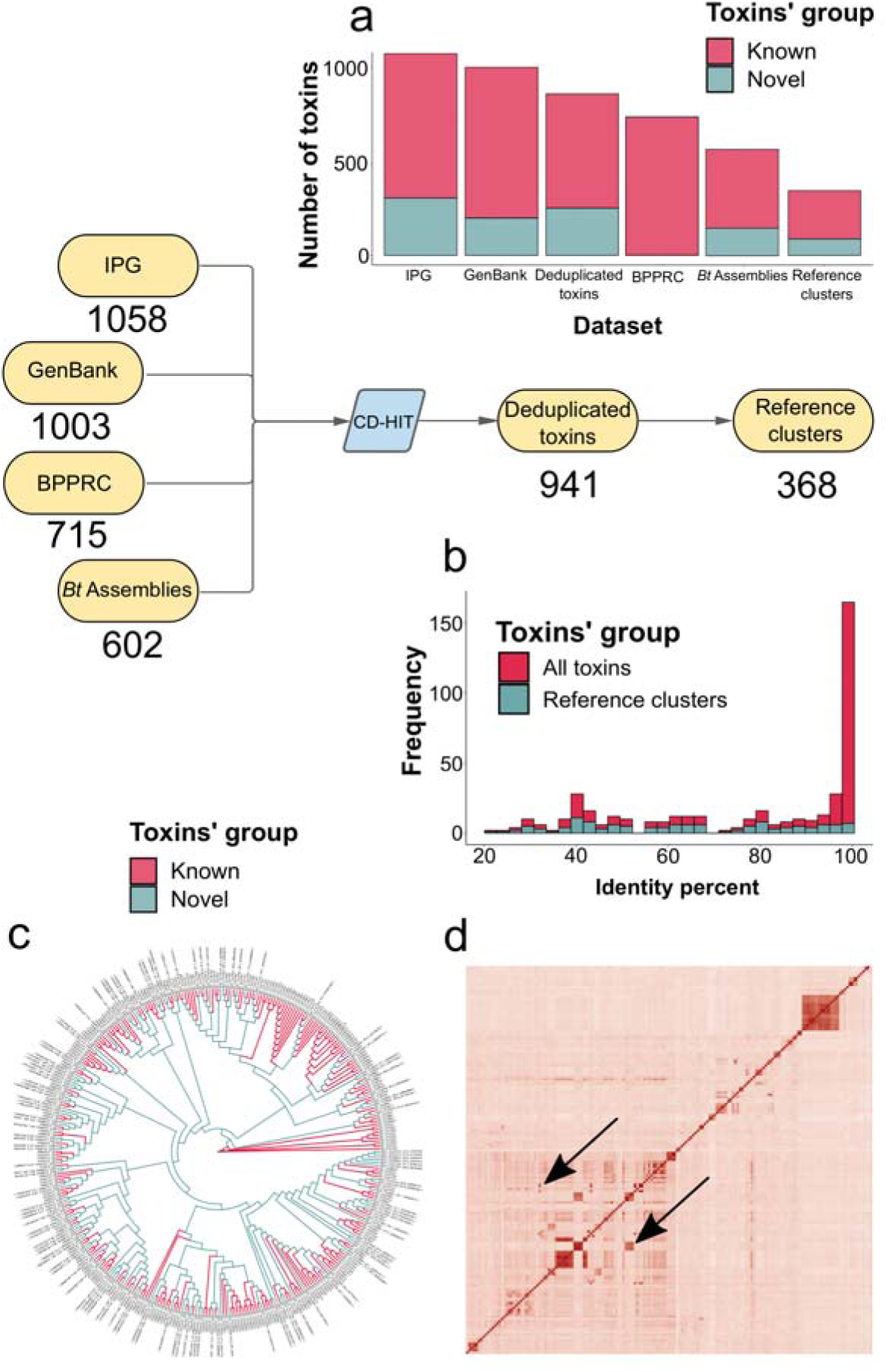
The properties of the Cry toxins dataset and reference phylogeny. (**a**) The distribution of Cry toxins amount extracted from four datasets, namely, NCBI Assembly, Genebank, IPG (identical protein groups), and BPPRC (Bacterial Pesticidal Protein Resource Center) databases. The color code denotes the type of toxins with a red color corresponding to known toxins deposited in BPPRC and a blue color designating putatively novel proteins. (**b**) The identity of the novel toxins with the closest homolog in BPPRC. The color code depicts the source of the dataset. Plotted are all unique toxins and reference clusters obtained using CD-HIT with a 95% identity threshold. (**c**) Reference ML-phylogeny (maximum likelihood) based on the concatenated alignments of the domains with partitioned evolutionary models. (**d**). The excised part of the heatmap with the pair-wise identity of sequences of the third domain. The order of rows and columns follows the above-described phylogeny. Arrows point to areas in which local identity does not comply with the expected order.

Most of the clusters were singletons (298) or doubletons (24); however, 7 clusters included over 10 toxins (fig. S2a, tab. S1). Within-cluster pairwise identity between sequences of the domains exceeded 99% (fig. S2b-c, tab. S4) and was independent of cluster size (fig. S2d). We then summarized codon-wise differences in the clusters’ alignments. Interestingly, non-synonymous mutations occurred more frequently than synonymous ones in all the domains (fig. S2e-f, tab. S5). Despite the first and the second domains harboring more substitutions (fig. S2f), after normalizing the non-synonymous ratio, the third domain was more divergent reaching 0.00051 compared with 0.0003 and 0.00035, respectively. (fig. S2g). This observation supposedly pinpoints a more intensive mutation rate and/or intensive recombination dovetailing with greater allelic diversity.

Of 941 unique sequences, 351 represented putatively novel groups absent in the BPPRC (fig. 1a, tab. S1) with 203 of them being substantially similar (>97%) to the closest entity from BPPRC which can imply point mutations and/or sequencing errors. In contrast, within reference clusters, 93 toxins were less than 95% similar to the closest homolog reaching 64% on average (fig. 1b). Notably, 30 toxins were attributed to other species and genera with 14 of them marked as novel, and some being remarkably different from the BPPRC references such as toxins from *Clostridium botulinum* and *Brevibacillus brevis* with identity estimates of 32% and 23% to Cry4Ba1 and Cry1Ie2 sequences (fig. S3a, tab. S6), respectively. No interdependence between the length of the domain and the identity with the closest deposited reference was found (fig. S3b). Hence, the aforementioned facts underline the abundance of undiscovered insecticidal potential yet to be harnessed, and at least some of the toxins might be revised for inclusion in the current nomenclature (Supplementary Text).

### Phylogenetic inferences assume independent evolution of the domains

We reconstructed phylogenies both from the sequences of the individual domains and a concatenated partitioned alignment (fig. 1c, fig. S4a-c). We compared the trees topologically using three metrics, i.e., quartet distance, Robinson–Foulds distance, and cophenetic correlation. The topology of the tree emanated from the third domain sequences varied sufficiently from the others according to the metrics (fig. S4d-f, tab. S7). The dissimilarity, albeit not so explicit, was reported for the phylogeny of the first domain as well. The phylogeny of the second domain seems to contribute to the shape of the tree based on all the domains more than others (fig. S4e).

To characterize the trees’ features, we evaluated mean support values, total branch length, tree balance indices, and the consistency index (CI) as a marker of homoplasy (tab. S8), being the most typical hallmark of recombination [46]. All the trees exhibited a high homoplasy rate with CI being less than 0.1 and were also unequally balanced. The third domain-based tree possessed the shortest branches. Our observations hence indicate that the recombination-driven domain III swapping is the most widespread type of exchange. In this regard, recombinant and parental sequences would group on the respective tree, while in other trees these recombination events are left unaccounted causing intense homoplasy and distortion of the molecular clock leading to longer terminal branches [46]. Altogether, the results presume that the evolution of individual domains could be considered independent being consistent with generally accepted views [6,22,24,30].

In order to demonstrate the exchanges in our data, we generated domain-wise identity heatmaps arranged to the order in the reference tree. If the evolution of the toxins is modulated by mutations only, one should expect diagonal similarity zones only, whereas unmatched areas could indicate possible recombination events. These inconsistencies were particularly evident in the case of the third domain (fig. 1d) and, albeit not as vivid, the first domain, while no conspicuous signals were tracked in the domain II-based heatmap (fig. S5a-c, tab. S9). These findings support the incongruity between tree topologies, disturbed branch lengths, and extreme homoplasy rate. Therefore, we hypothesized that domain exchanges drive the evolution of Cry toxins.

### Recombination between Cry toxins occurred multiple times during their evolution

To study the presence of domain shuffling, we utilized the RDP4 software and revealed 120 events accordingly (tab. S10). Participants in the events were classified as recombinants as well as minor and major parents that transferred only one domain or two domains jointly, respectively. If putative parents were absent in the dataset, they were called unknowns. After that, we performed a two-step filtration by discarding the exchanges in which less than 70% of the domain was involved (marking them as partial). We also verified the congruence in the domain-wise phylogenetic trees. In brief, we inspected whether at least one parent and children formed a compact phylogenetic clade, and, in this case, updated the list of parents for a single domain if the clade contained extra toxins (Supplementary text). The number of reports was then reduced to 100 and 50 events, respectively (tab. S11, tab. S12). There was sufficiently more swapping of the third (33) and first (13) domains, while only 4 events pertaining to the second domain were detected in the final dataset, and this pattern was a general feature for all the datasets (fig. 2a, fig. S6a-c). It is worth noting that most of the breakpoints were arranged around the boundaries between the domains (fig. 2b, fig. S6d-f). Apart from reducing the total number of inferences, the filtration allowed discarding events with a large number of participants (fig. S7a-b, tab. S13), decreasing the percentage of unknown parents (fig. S7c), and, conversely, increasing similarity between recombinants and parents within the transferred regions (fig. S8a, tab. S14) as well as the difference with non-transferred domains (fig. S8b, tab. S15). This procedure lowered the number of minors and, contrarily, leveled up the abundance of majors (fig. S9a, tab. S16). We also found that 14 events were characterized by more parents providing only a single domain which might evolve from possible ancestral exchanges, especially engaging the third domain (fig. S9b).

**Fig 2.**
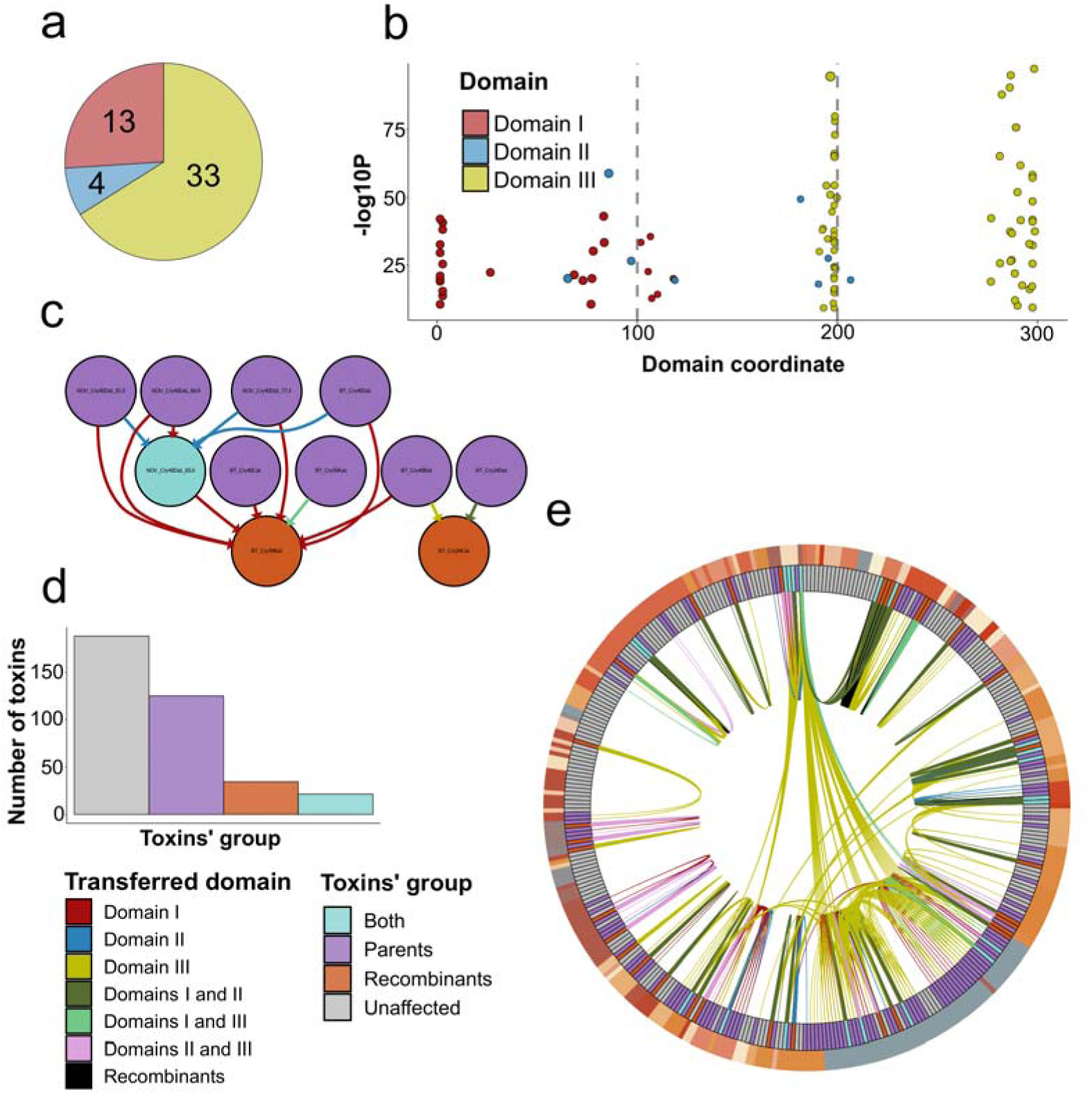
The recombination landscape of 3-D Cry toxins. (**a**) The total number of recombination events identified by the RDP4 utility and classified according to swapped domains. Shown are data for the filtered dataset, i.e., including events with more than 70% of the respective domain subjected and forming compact clades within domain-wise phylogenetic trees. For a detailed description of the filtration process, consult the Supplementary text. (**b**) Distribution of recombination breakpoints. On the x-axis, the relative coordinate in the sequence is presented. Plotted on the y-axis, is the mean p-value from all recombination detection tests implemented in the RPD4 software. Grey dotted lines mark the boundaries of the domains. (**c**) An example of one weakly connected component extracted from the graph of domain exchanges (fig. S11). The color of the node displays the role of the toxin in recombination events, i.e., recombinants, parents, and those participating in multiple swaps being parents and recombinants simultaneously. Edges’ colors denote the type domain(s) transferred, namely, a single domain for minor parents and two domains for major parents, respectively. (**d**) The number of toxins according to their type in recombination events. (**e**) Circos plot illustrating the map of domain exchanges with sectors’ order corresponding to reference phylogeny. The color of the inner circle represents the toxins’ types in recombination events. Links are colorized in accordance with the domain transferred. Black curves connect toxins if a particular event contains more than one recombinant. The inner circle marks the first-rank Cry toxins class according to the established Cry nomenclature.

Finally, we analyzed the domain-wise number of mismatches between parents and recombinants within the transferred regions normalized by the alignment length. There was a significant difference between the mismatch rate of minors and recombinants in the third domain compared with the respective rate within the first and the second domains between majors and recombinants reaching the median values of 0.23, 0.14, and 0.18 accordingly (fig. S9c, tab. S17). This presumably reflects intensive swapping of the third domain that affected both recombinants and parents in other putatively unaccounted events that is supported by the inflation of parents providing the third domain when correcting the list of parents depending on the content in the subtree of domain III (fig. S9b) and by the higher normalized non-synonymous ratio in the clusters pooling more than 10 toxins as well (fig. S2g). The results further prove that recombination sparks elevated levels of allelic diversity, and the more frequent exchanges happen, the more non-synonymous mutations emerge.

We then studied the recombination rate using ClonalFrameML, Gubbings, and fastGEAR which reported 13290, 369, and 1711 events, respectively (fig. S10a-c, tab. S18-20). The programs provided incommensurable ratios of effects of recombination and mutation (r/m), namely, 372.2 and 1.1 estimated by ClonalFrameML and Gubbings. Due to the high number of events, the latter seems to be an underestimation. A substantial incongruence might be explained by inaccurate predictions when dealing with extremely divergent and intensively recombining sequences [46]. However, the abundance of the events corroborates the presence of extensive recombination between Cry toxins.

To draw a map of domain exchanges, we built a directed graph with nodes representing toxins, and edges corresponding to the transferred domains regarding possible ancestral events derived from the filtering procedure (fig. 2c, fig. S11). The graph is constituted of 310 edges, 180 nodes, and 22 connected components. Of these, the first component encompassed 19 events, five components included from 2 to 5 events, whereas the rest 16 components were singletons (fig. S12a-c, tab. S21). We also categorized toxins with regard to their role in recombination as parents, recombinants, and the ones being both donors and recipients (fig. 2d). These classes constituted 125, 34, and 21 toxins, respectively (tab. S22). It, therefore, follows that 48.9% of the toxins from the reference dataset took part in recombination. All the reference sequences from the clusters uniting more than 10 toxins were parents. We also took all nodes and edges from the graph and mapped them on the reference phylogeny to design the Circos plot arranged according to the order of the tree built from the concatenated partitioned alignment which vividly illustrates both the intensity of the exchanges and their influence on phylogeny (fig. 2e). The combination of two approaches, namely, adding parents from domain-wise phylogenetic trees per recombination events and creating the exchange graph allowed us to discover three pairs of linked events in which the same toxin was spotted (fig. S13, tab. S23). In contrast to other events, these toxins received each of the domains from distinct parents possibly implying that true major parents are absent in the dataset of currently available sequences of Cry toxins. Notably, two events involved the second domain, which suggests that such findings could arise from two sequential swaps of the first and the third domains. Since two other events in which the transferring of the second domain was more evident were present (tab. S12), we do not exclude the probability of domain II exchanges, while they seem to be quite rare if exist. To deal with unknown parents, we proposed an algorithm relying on the depth of the phylogenetic clades combining parents and recombinants within trees devoid of toxins with domains of unknown origin (Supplementary text). In brief, we calculated the ratio of nodes with parents and recombinants and the mean number of parents in these phylogenetic inferences within the given depth. We selected minor parents and major parents for adjacent domains omitting the opposite ones to exclude distortion due to possible ancestral exchanges (tab. S24). Roughly perceiving domains as full toxins, we predicted that at least 40 toxins were putatively absent in our dataset.

It is worth noting that toxins that are both parents and children predominantly reside in the largest connected components in the graph presuming the presence of bacterial populations with more frequent domain exchanges. However, the presence of numerous events *per se*, albeit implying biological importance, neither provides functional consequences of such processes nor explains what causes this intergenic recombination. Therefore, we proceeded with studying the genomic surroundings of *cry* genes.

### Domain exchanges are driven by inter-domain conservative regions and flanking insertions

To uncover possible mechanisms orchestrating domain swapping, we first compared the recombination rate in *Bt* genomes bearing toxins either with exchanges detected or devoid of them. We used 467 *Bt* assemblies, with 169 comprising 3-D Cry toxins and 65 including parents/children in recombination events. We reconstructed the pangenomes using these genomes and launched ClonalFrameML on core gene alignments accordingly. All the pangenomes were rich in accessory components, and only 7-10% of core genes were reported (fig. S14a-c). Moreover, pangenomes were considered open due to the alpha parameter of the Heaps’ law ranging from 0.61 to 0.71 implying high genetic plasticity (tab. S25). The ratio of rates of recombination and mutation (R/θ) ranged between 0.07 and 0.30 indicating that mutations, in general, happen more often than recombination (tab. S26). Nevertheless, the r/m scores were higher, reaching 2.54, 2.38, and 0.85 for the datasets, namely, all *Bt* assemblies, those having *cry* genes, and genomes harboring participants of recombination (fig. S14d-f, tab. S26).

The fact that the dataset with *cry* genes engaged in recombination showed a lower r/m rate may imply that the intensity of homologous recombination *per se* does not explain the emergence of domain exchanges. Next, we searched for inclusions of *cry* loci into mobile genetic elements (MGEs), in particular, insertions (IS), prophages, and genetic islands (GIs). The number of these elements was moderately yet significantly (p < 2.2e-16) correlated as the respective Pearson coefficient hovered between 0.39-0.49 (fig. S15a-c, tab. S27). It thereby might be hypothesized that if the genome is rich in some groups of MGEs, e.g., genetic islands, it will likely house more mobile elements of other classes. Interestingly, *cry*-harboring assemblies, both with or without domain swaps, incorporated a significantly higher portion of MGEs (p < 1e-08 according to Wilcox test) compared to all assemblies, and no difference between the former two groups was detected (fig. 3a). When inspecting which *cry* genes fell into specific MGEs, we identified that of all 602 toxins, 172 Cry-encoding genes resided in GIs, and 15 were located within IS, whereas no relations with prophages were reported (tab. S28). Of note, 36% of *cry* genes that had undergone recombination were associated with GIs and IS vs. 28% of all toxins (fig. S15d). However, Fisher’s exact test demonstrated no significant difference (p = 0.102) between these groups. It is noteworthy, that several *cry* loci were flanked by ISs as shown for the strain HS18-1 (fig. 3b).

**Fig 3.**
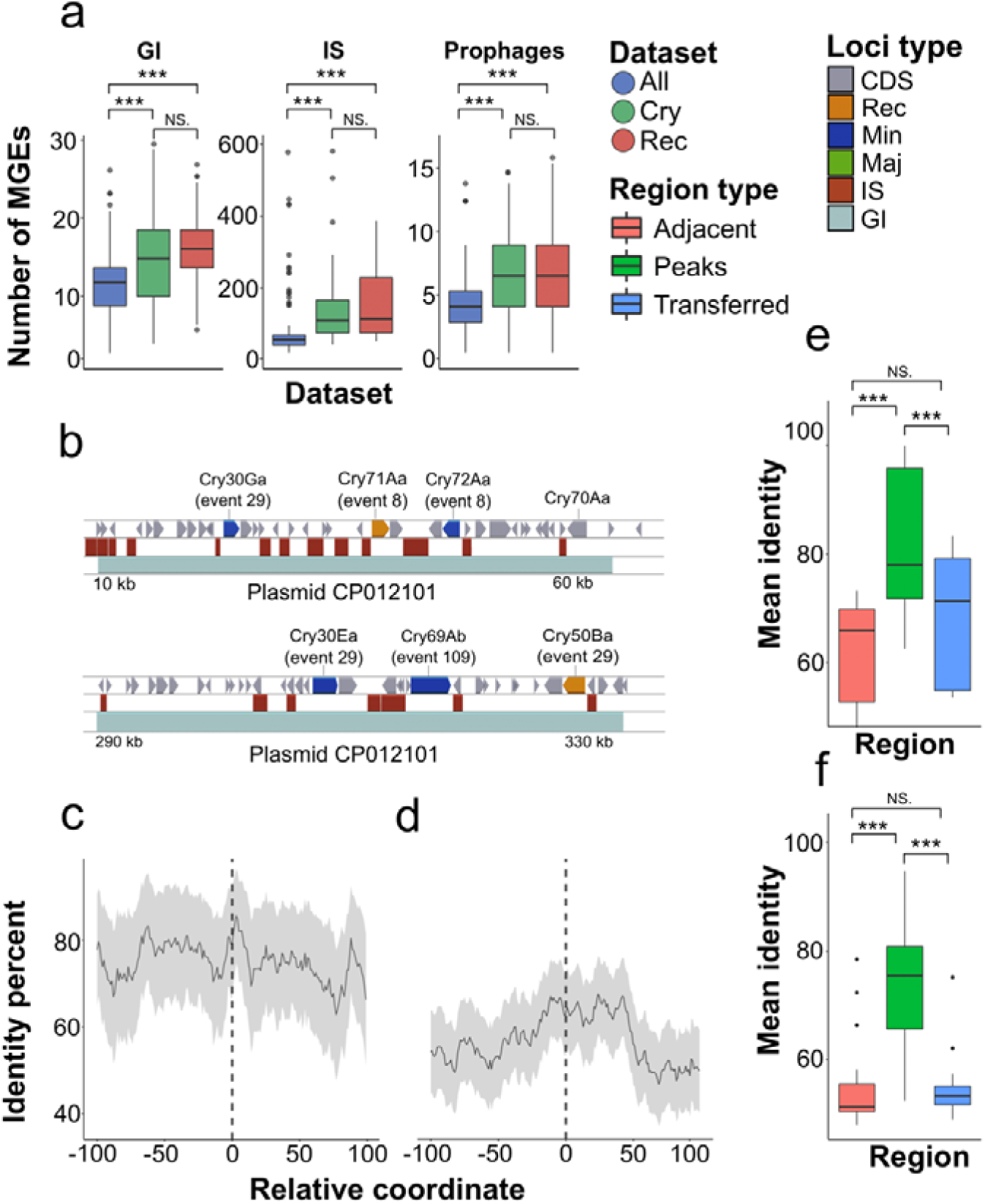
Putative mechanisms of domain exchanges driven by mobile genetic elements and domain-wise identity within breakpoints. In all plots, asterisks denote significant differences according to the Wilcox test with p-values adjusted using the Benjamini-Hochberg procedure. (**a**) The average number of MGEs (mobile genetic elements), including prophages, insertion sequences (IS), and genetic islands (GIs) presented in *Bt* genomes. The color of box plots refers to the sets of *Bt* assemblies, namely, all genomes (“*All*”), assemblies bearing *cry* loci (“*Cry*”), and those incorporating participants in domain exchanges (“*Rec*”). (**b**) Loci surrounding genes encoding recombinants and parents from individual recombination events in the strain HS18-1. The color of the blocks on the first track corresponds to the toxins’ role in recombination events, e.g., major and minor parents, recombinants, and unaffected toxins. (**c**) Coordinate-wise mean identity between parental sequences in recombination events within two regions located between the first and the second domains and between the second and the third domains (**d**). The identity is calculated for 100 b.p. flanking regions before and after the breakpoints. The dashed lines point to the breakpoints. Grey areas represent the standard error. (**e**) The comparison between the mean identity of peaks within the event-wise clusters of flanking regions and sequences of the whole domains either those that were transferred from the minor parent to the respective recombinant and the adjacent non-transferred domains. Shown are comparisions for regions surrounding the first and the second domains (**e**) as well as the second and the third domains (**f**), respectively.

Next, we set out to find conservative blocks within the boundaries of the breakpoints that could serve as focal points for domain exchanges. To achieve that, we selected recombination events without unknown parents and aligned the respective sequences to get the coordinates of breakpoints in parental toxins. Thereupon, we re-aligned the parents only and, through mapping to the initial coordinates, excised 100 b.p. flanking regions surrounding the breakpoints (tab. S29). After that, we ran the rolling average function on the coordinate-wise per-site identity. We found that both breakpoints (between the first and the second domains and between the second and the third domains) we surrounded by local peaks flanked with two local minimums (fig. 3c-d). To make a more detailed picture, we carried out a clusterization regarding the event-wise per-site distribution of identity scores. We first opted for the most suitable clustering algorithm and distance metrics by determining the highest silhouette score. The best option implied cosine distance applied on rolling mean-corrected identity scores with the following hierarchical clustering (fig. S16a-b, tab. S30). The optimal number of clusters reached 2 and 6 for the first and second breakpoints, respectively (fig. S16c-f, tab. S31). According to the inter-cluster identity distribution, distinct clusters displayed from one to three peaks offset from the breakpoints (fig. S17a-b). Following that, we summarized per-site identity within these peaks and compared them with the respective values of the full domain sequences, both transferred from the minor parent and the adjacent non-transferred domains (fig. 3e-f). The average within-peak identity constituted 81% and 73% for the first and the second breakpoints vs. 62-68% and 54% for the respective adjacent and transferred domains (tab. S32). All the comparisons were significantly different according to the Wilcox test (p < 0.05).

These results show that the richness in chimeric toxins is not delineated by the overall rate of homologous recombination but rather relies on the enrichment of MGEs within the genome. The loci encoding Cry toxins, often located within genomic islands, are also flanked by insertion sequences. Given the presence of highly conserved regions in the boundaries between the domains, we suggest these homologous regions coupled with insertion surrounding the respective genes might serve as breakpoints for domain swapping.

### Recombination leads to changes in host specificity and toxins’ composition in strains

Having shown the prevalence of the domain exchanges between Cry toxins, we proceeded to study their impact on the spectrum of toxic activities. We first compiled the available information on affected hosts within the fourth and third rank of Cry proteins utilizing the specificity database from BPPRC, comprehensive reviews [47–49], and other literature sources (research articles and patents) (tab. S33). Since the data for putative novel toxins often remains unknown, we also searched for the insecticidal activity of bacterial strains producing Cry toxins. As a result, we collected host range data for 284, 184, and 148 proteins within the fourth and the third ranks, and strains, respectively (fig. S18a). Unsurprisingly, the number of individual assays predominantly included orders Lepidoptera, Diptera, and Coleoptera accounting for 75-85% of the examples (fig. S18b, tab. S34). For most cases, proteins and strains exhibited activity against a single order, whereas there were two notable instances such as Cry1Ba and the C18 strain related to four different orders (fig. S18c, tab. S35). A similar picture was obtained for species (fig. S18d). However, it should be said that three Cry toxin families, namely, Cry1Aa, Cry1Ab, and Cry1Ac, being the most studied, were toxic to a broad range of host species ranging from 48 to 65 (fig. S18d).

We first focused on host orders for participants of recombination exchanges by using three approaches. We attributed toxicity based on the fourth and the third toxins’ ranks, and imputed data associated with strains if the respective information for proteins was unavailable. Only those events in which annotations for at least one parent and child were present were studied. Due to the fact that the abovementioned parents of a single domain may be a trace of ancestral events, we retained only true majors hereinafter. Using three counting strategies, 19, 22, and 30 events were reported (fig. 4a, fig. S19a-b, tab. S36-38). Differences in the affected orders constituted a higher fraction of events for each dataset. In the events where the orders remained unchanged mainly anti-lepidopteran and nematocidal toxins were involved, while those infecting hemipteran and dipteran pests fell into the category with a non-identical host range. It is noteworthy, that host specificity switching seems to be related to the identity of the transferred domains between recipients and donors, and the number of toxins as well (fig. S19c-e, tab. S39). Logistic regression revealed that the only statistically significant variable (p<0.03 according to Wald z-statistic) determining unequal host orders was the number of toxins in the event. Such events might be more distant than those, presumably recent, comprising only one child, minor and major parent, respectively.

**Fig 4.**
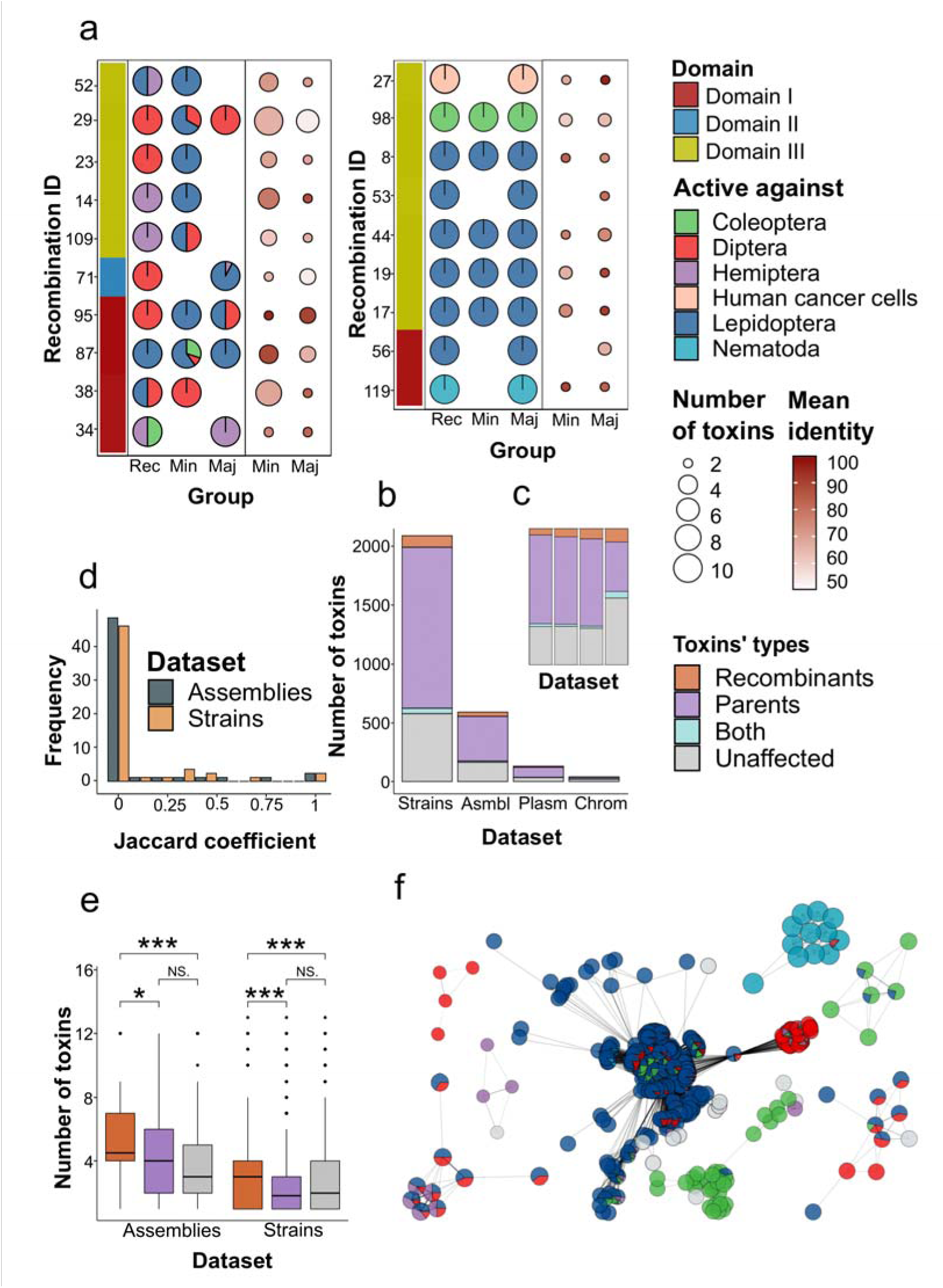
The impact of recombination on hosts specificity of Cry toxins and toxins’ content in bacterial strains. (**a**) The percentage of species attributed to different orders against which recombinants and parents are active. The data on the toxicity spectrum is based on the fourth rank of the toxins. The portion of host orders within the pie charts is color-coded. The events are split into two panels, namely those in which at least one toxin exhibits the activity to host orders differing from other types of parent and/or child. The left adjacent bar is designated to the domains transferred in the events. The right adjacent panel shows the number of toxins. ‘*Rec*’ indicates recombinants, ‘*Min*’ – minor, and ‘*Maj*’ – major parents, respectively. The size of the dots relies on the number of participants, whereas the intensity of the color delineates the mean pair-wise identity between the transferred domains. (**b**) The number of particular toxin types in the datasets such as strains, assemblies, plasmids, and chromosomes containing recombinants, parents, and unaffected toxins, and the fraction (**c**) of these toxins’ groups. (**d**) The similarity between all sets of toxins that are housed by strains and assemblies in which at least one parent and recombinant reside using the Jaccard coefficient (the length of the intersection divided by the union of the sets). Event-wise comparisons refer to toxins’ compositions in the assemblies and strains attributed to recombinants and all parents of the recombination events. (**e**) The number of toxins comprised in the assemblies and strains bearing recombinants, parents, and *cry* loci devoid of recombination signals. Asterisks denote significant differences according to the Wilcox test with p-values adjusted using the Benjamini-Hochberg procedure. (**f**). Selected connected components from the weighted specificity-wise graph of strains sharing the common Cry toxins. The nodes represent strains, whereas edges are drawn if at least one toxin within the third rank is found in two connected strains. The weight of the edges is determined by the number of shared toxins. The color of the nodes visualizes the overall proportion of species attributed to the orders that the toxins produced by the strain exert the effect on.

Next, we examined the range of species to which parents and recombinants were toxic. Due to the imbalance in the number of tested species for several Cry holotypes (fig. S18d), we decided to use the Szymkiewicz-Simpson coefficient as a similarity measure between the sets of hosts that belonged to donors and acceptors of the domains. Unlike tracking changes in orders, the relationship between the coefficients and the number of toxins was not straightforward: in some events with multiple toxins involved, host ranges were highly close while others bore only one toxin of a certain type but displayed no intersections between the host sets (fig. S20a, tab. S39). By using linear regression, we identified that the degree of similarity significantly depends (p < 0.048) on the type of parent. More common hosts between recombinants and major parents in comparison to minors are observed (fig. S20b). In addition to qualitative differences, we traced the impact of recombination on the number of reported toxic activities and thus considered children, parents, and toxins devoid of recombination signals. Concerning the number of orders, irrespective of which information was taken into account, the differences were non-significant, and the mean values hovered around 1.14-1.34 (fig. S20c, tab. S40). Nonetheless, if the third rank-related data were included, the number of affected species for parents was slightly higher than within toxins without detected swappings, namely, 3.7 vs 2.7 (fig. S20c, tab. S40). This observation was retained in case of keeping data from well-studied toxins with the number of hosts exceeding 15 (fig. S20d). As such, the results suggest that recombination does not increase the absolute number of infected hosts but more likely alters host specificity.

To characterize the possible relationships between domain exchanges and the composition of Cry-encoding genes in isolates with such chimeric proteins, we summarized all toxins attributed to certain assemblies and strains (tab. S41-42). Considering *cry* sequences in the fourth rank-wise rank led to overestimations sparked from sequencing errors, especially if *cry* genes of the strain were sequenced both by Sanger and NGS methods (fig. S21a-b, tab. S41). As a result, the extensive number of *cry* genes was reduced sufficiently if using the third rank, and we retained it for further examinations (tab. S41). In order to understand, if and how the presence of recombinants in the isolates shapes the composition of toxins subjected to recombination or unaffected ones, and also characterize the patterns of co-occurrence of domain donors and acceptors, we classified assemblies and strains concerning the types of toxins they possessed and evaluated the inter-group intersections. Here and in further analysis, we considered the datasets in two ways: (i) the strains/assemblies were attributed to toxin types (parents, recombinants, and the unsubjected ones) if they comprised at least one toxin of a certain type regardless of the other types; (ii) the strains/assemblies containing exclusively one type were considered (e.g., those including recombinants but not parents and/or non-chimeric toxins, etc.). We found that parents and children from diverse events tend to co-occur in the same assembly/strain (fig. S21c-d). Of those harboring only a single type of toxins the ones harboring parents prevailed (324 strains). The second largest group encompassed both parents and intact *cry* sequences (254 strains). We hence could propose that these instances might hide potential events left undetected due to the limited number of available *cry* sequences. Following that, we looked at the content of Cry toxins within strains, assemblies, contigs, plasmids, and chromosomes to find which genomic regions harbor *cry* genes. According to median estimates, strains, and contigs were singletons, assemblies bore 3 *cry* genes, whereas plasmids and chromosomes – 2 (fig. S21e, tab. S43). The results might be explained by the absence of genomes for a plethora of strains for which only *cry* sequences are available and the high fragmentation of assemblies, respectively. Unlike the frequency of toxin types on the basis of their role in recombination, when tallying up third rank-wise distribution within the datasets, participants of domain exchanges accounted for 73% of all Cry toxins (fig. 4b-c). Of note, focusing on the entities incorporating specific toxins groups revealed that the type of interest prevailed in these datasets. More specifically, recombinants constituted 30%, parents – 79%, and unaffected by recombination toxins – 65% of the data within the respective subsets (fig. 21f-h).

Having found that isolates harboring at least one recombinant are generally enriched with both parents and recombinants, we further assessed the impact of individual recombination events on toxins’ content and abundance. To achieve that, we picked all strains or assemblies in which parents and recombination resided and calculated the Jaccard coefficient using all toxins in these strains both when comparing children with parents and minor/major parents only. We observed a strikingly high prevalence of distinct toxin sets in both analyses (fig. 4d, fig. S22a, tab. S44). No overlaps were observed for almost 70% of the events, namely, 34 and 35 for the above-defined comparisons. The median coefficient for parents only slightly but insignificantly exceeded the respective measurement for child-parent comparisons (0.38 vs 0.52) (fig. S22b). As the domain transfer could also entail the acquisition of new toxins due to the concomitant propagation of MGEs, we took the assemblies bearing *cry* genes subjected to domain exchanges and the unaffected ones. The presence of parents and especially recombinants is significantly associated with a greater number of toxins in contrast to the unsubjected isolates reaching median values of 4.5, 4, and 3, respectively (fig. 4e, tab. S45). On the other hand, expanding the scale to strains, only representatives including recombinants bore 3 toxins vs. one and two, respectively (fig. 4e). If scrutinizing strains that contain recombinants, parents, or intact toxins exclusively without other types, most of them harbored one toxin only (fig. S22c). There were no assemblies that bore recombinants without at least one toxin of another type, whereas genomes with parents carried significantly more *cry* genes (fig. S21c).

Owing to the fact that there were some cases in which parental and children toxin sets showed intersections (fig. 4d), we further set out to find whether they could collocate on the same genomic regions or reside in the same genome. Six events were identified with five of them displaying recombinants with one of the minor parents, and one with both types of parents (tab. S46). Toxin-encoding genes were located either within close proximity to each other or in different regions. Noteworthy, strain HS18-1 was associated with two events with *cry* genes residing within two large pathogenic islands (fig. S23a). Genetic loci coding for toxins Cry71Aa (recombinant) and Cry72Aa (minor parent) from event 8 were linked with the first island, as well as Cry30Ga (minor parent) from event 29 with the respective child (Cry50Ba) being adjacent to another minor parent (Cry30Ea). The second pathogenic island also housed a minor parent from event 109 (Cry69Ab). As stated in the previous section, the *cry* genes described were flanked by arrays of insertion sequences, and this pattern was demonstrated for other events as well (fig. S23a). We also counted the overall number of MGEs in the genomes accordingly. Strains HS18-1, C15 (event 104), and MYBT18246 (event 73) were enriched with GIs, and the latter two were also rich in IS and exhibited nematocidal activities. The remaining assemblies were congruent with median values (fig. S23b). The sequences of the domain donors and acceptors for all 8 events forming sister clades on the reference tree suggest them as relatively recent (fig. S23c).

We then attempted to pry out whether the strains/assemblies carrying chimeric loci exhibit a broader host range. To avoid a possible bias in comparisons due to the presence of well-studied toxins, we performed two series of calculations. The first implied including all datasets, and the second approach was based on the dataset excluding Cry types toxic to over 15 host species. For convenience, we would call these series of calculations trimmed and untrimmed. Within the fourth rank, the strains and assemblies harboring toxins that have undergone recombination and parents seem to exert an effect on more orders compared to unsubjected Cry toxins, which, however, was mostly insignificant (fig. S24a, tab. S47). In contrast, third rank-based metadata provided a significant difference, whereas the inferences were incongruent. In the trimmed mode strains/assemblies with recombinants were associated with 3 orders, parents and toxins devoid of exchanges with 1-2, respectively (fig. S24a). If considering the untrimmed series of calculations, entities with parents were linked with 3-4 orders, which significantly exceeded those with domain acceptors (2-3), and the unaffected ones (1-2) (fig. S24b). As with orders, fourth rank-derived data of species provided insignificant comparisons (fig. S24c). Within the third rank-wise trimmed datasets, assemblies and strains that bore *cry* loci that received domains were related to 7, parents – 3, and toxins devoid of recombination– 4-5 host species (fig. S24c). Accounting for well-studied toxins gained 9, 43-63, and 5-7 infected species within the mentioned groups accordingly (fig. S24d, tab. S47). While the inflation in the number of hosts gained inconsistent comparisons between parents and children, it is clear that the presence of any recombination participants broadens the host range which is possibly due to the greater overall content of Cry proteins consistent with previous findings (fig. 4c).

Finally, we made a summary of the data by designing a weighted specificity-wise graph with nodes corresponding to individual strains and edges denoting common toxin sets of the third rank to overcome artifacts sparked from assembly errors (fig. S25, fig. 4f). The graph was composed of 948 nodes forming 169 connected components, and 114 of them were singletons (fig. S26a). The largest component was formed by 556 strains, 23 were comprised of 4 to 36 strains, and the rest included 2-3 nodes (tab. S48). The number of components possessing participants in recombination events was 73 (tab. S49), and the largest one incorporated 16 events. Significant dependence (p < 0.01 as shown by linear regression mode) between the component’s size and the number of events was revealed (fig. S26b). We then analyzed the toxin content of components into which parents and children fall using the Jaccard coefficient. Strikingly, even with this extended data, the same picture as with strain-wise toxin compositions remained. Irrespective of the parent type, when comparing parents only, for more than 50% of the events (26-31) no intersections were reported (fig. S26c, tab. S49). We compared the properties of the components devoid of recombination signals in the toxins they housed considering the number of strains, individual toxin types, and affected host orders. Significant differences (p < 0.01 according to the Wilcox test) were observed only for the number of toxins (fig. S26d-f). The sets of orders to which toxins in the components show activity most frequently were presented by singletons linked with only single host types. Nevertheless, there were examples of generalist groups, mainly those with plenty of strains (fig. S25, fig. S27a). The proportion of these generalist strains was slightly higher in components with toxins undergone recombination in contrast to the intact ones (fig. S27a). Moreover, the former group comprised a higher percentage of components with at least one known host attribution (tab. S48), namely 63% vs. 50% (p<0.02 according to Fisher’s exact test). Intriguingly, the components with nematocidal toxins often collocate with those active against orders Hymenoptera and Hemiptera as well as parasporins (fig. 27a). By plotting the distributions of order-wise toxicity spectra regarding particular events, we noticed that, in many cases, different toxins fell into both specialists and generalists’ categories, which was most evident for anti-dipteran and anti-lepidopteran strains (fig. 27b). Finally, we calculated the similarity between hosts that toxins/strains within the components harboring parents and children exert effect on. Unlike comparing toxins *per se*, linear regression has not determined that the parent type contributes to the fraction of shared hosts (fig. 27c-d, tab. S50). Szymkiewicz-Simpson coefficient demonstrated that the components tested in most cases overlap sufficiently, while the Jaccard coefficient was skewed towards lower values corroborating previous observations being the possible marker of exchanges between generalists and specialists.

In order to prove our suggestions, we combined the strain graph with recombination events and linked the connected components by edges attributed to domain transfers (fig. S28a-b). The graph clearly showed that larger connected components with extensive host range contain more edges lining them with smaller components with specialized strains. Keeping in mind that the recombination graph resembles this pattern (fig. S11), we speculate that after the contact of the populations adapted to a certain host, the exchange of MGEs housing *cry* loci with the concomitant domain swaps leads to the emergence of more successful strains with an expanded list of affected hosts. We then proceed with studying the exchanges within the context of selection pressure.

### Domain swapping is accompanied by purifying selection

Once recombination occurs, the chimeric sequences are relatively often subjected to evolutionary selection, and we hence analyzed whether domain exchanges are linked with selection signals. To track such signals, we used LRT (likelihood ratio test) on the estimates of branch, site, and branch-site models within clades uniting toxins with domain exchanges reported. The respective procedures were applied to the domain sequences independently using phylogenetic trees adjusted for recombination (Supplementary text). Forasmuch as domain-wise parents were considered, we took all parents within the domain without omitting those that are not true majors.

We first focused on the branch model by marking either children or parents on the trees. After performing calculations, 83 signals were detected, and only 4 of them represented positive selection, whereas the majority were associated with purifying selection (fig. S29a, fig. 5a, tab. S51-52). The respective signals encompassed 26 events in total, and 19 of them remained when considering only those with recombinants marked with 35 individual signals for the latter group. We classified the events if the usage of domain-based trees gained patterns absent in the respective data for parents, and reported such inconsistencies in 13 events (fig. 5a, tab. S52). In some cases, especially if the first domain was transferred, selection influences the domain that was received from a minor parent while others showed the opposite (fig. S29a). Interestingly, two instances subjected to positive selection were presented by chimeric events (103 and 45) in which the recombinant obtained the first and the second domain from distinct parents with an unknown source of the third domain (fig. S13). The two-step filtration of the events based on the LRT test significantly changed the mean ω for the foreground branches reaching 0.52, 0.71, and 0.16 for all events, non-neutral yet identical within branches, and with unequal rates in the foreground branches (fig. S29b, tab. S52) being associated with the number of marked toxins (fig. S29c-d). At the same time, the mean ω for the background remained unchanged and hovered around 0.367. Notably, foreground ω within the threes built on diverse domain sequences exhibited no significant differences (fig. 5b, fig. S29e).

**Fig 5.**
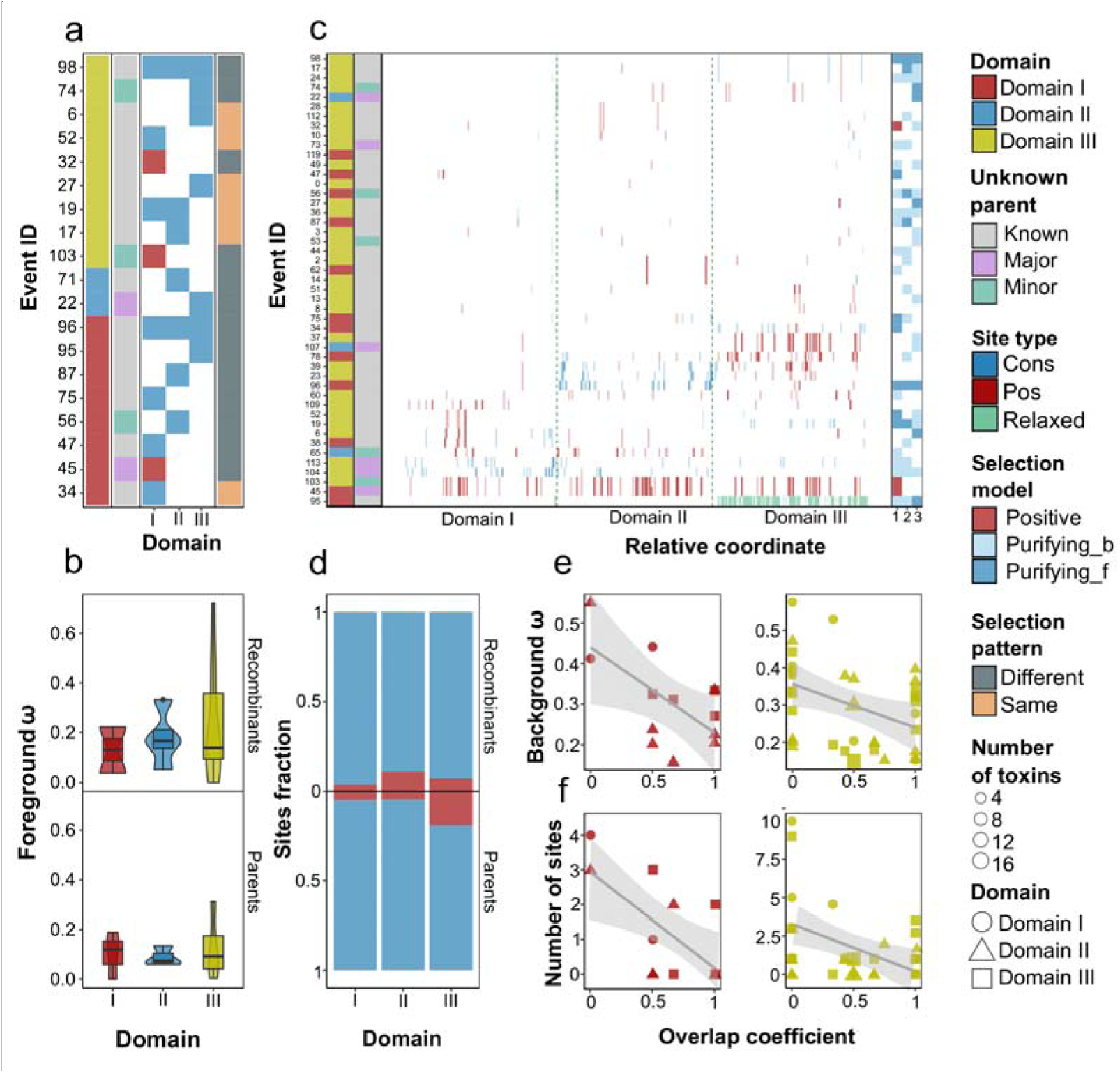
The relationships between domain exchanges and evolutionary selection signals in *cry* sequences. (**a**) The optimal evolutionary models calculated on branches in which recombinants were marked. Here and in other figures the selection pressure was determined on the sequences of different domains independently. The heatmap shows the models reported for the individual domains. The color of the blocks indicated whether the model chosen implies positive or purifying selection. The left adjacent subpanels correspond to the domain that was transferred from the minor parent and show whether a parent, either minor or major, is unknown, i.e., absent in the dataset. The right subpanel indicated whether for at least one domain a different selection pattern from the respective parental sequence was identified. (**b**) The average ω values (the non-synonymous to synonymous substitution rate ratio, dN/dS) based on branch models for the events affecting the third domain of parents and children, respectively. (**c**) The distribution of sites with significant signals of evolutionary selection reported by best-fit site models opted by the LRT (likelihood ratio test). The left adjacent subpanels reflect the same information as presented in Figure (**a**). The site color represents whether the site is positively selected, conservative, or relaxed. The intensity of the color depends on the probability ranging from 0.95 to 0.99. The right subpanel indicates the significant model on branches with recombinants marked, namely, positive and purifying selection either for the foreground or background only if the ω on branches is equal. (**d**) The percentage of positively selected and conservative sites determined using branch-site models for recombinants and parents within the events in which the third domain was transferred. (**e**) The relationship between the set of common species which recombinants and parents are active against and the ω or the number of positively selected sites (**f**), obtained from using branch and site models, respectively. The color indicates the domain transferred, while the shape of the dots encodes the domain for which the analysis was carried out.

We moved on to classify individual sites. A total of 228 significant signals with one M1, 69 - M3, and 157 - M8 optimal models were chosen indicating relaxation and positive selection on sites for the latter (tab. S52). For event 95 associated with the M1 model, the enrichment with relaxed sites was reported, while for most cases belonging to the M3 model, only a limited number of significant sites were reported, and they predominantly were located on the boundaries between the domains opposite from the ones transferred from the minor parents (fig. 5c). The highest portion of evenly distributed positively selected sites was a distinctive feature of all linked events (fig. S13). Other inferences represented sites concentrated in the middle part of the domains (fig. 5c). The number of positively selected and conservative sites reached 369 and 259; thus, 59% of sites were subjected to positive selection (fig. 30a, tab. S53). Regarding the sites’ distribution in recombination events, we discovered that when the first and the third domains are exchanged, a higher fraction of conservative sites is seen within the second domain (fig. S30b-c). Nevertheless, we have not found significant differences between the total number of positively selected and conservative sites in the domains which could be explained by individual but not average patterns in the particular events (fig. S30d-e). To figure out the factors delineating the number of positively selected sites, we compared the respective values with the number of toxins and the ω, both background and foreground in trees reconstructed to calculate branch models for parents and recombinants. Neither the total number of toxins nor the marked ones have shown any impact on the number of sites possibly indicating that obtaining a tree for revealing selection on sites did nor not influence the assessment procedure (fig. S31a-b, tab. S52). The foreground ω, both when considering all signals and those with significant selection signals has no relationship with the number of sites (fig. S31c-d), while a positive correlation with the background ω was detected (fig. S31e-f).

By analogy with the site models mentioned, we performed the same procedure to pick the best-fit branch-site models. From a total of 246 signals, only one was the mA model with the rest corresponding to mB models (tab. S52). Obviously, the model chosen tends to inflate the number of positive selection signals for some events marking the whole domain instead of providing particular sites (fig. S32a-b). Notably, events with sporadic relaxation sites were instead enriched with conservative regions except for event 95 which had prolonged relaxation regions and turned to a positive selection signal for the whole domain (fig. S32a-b). We compared the proportion of positively selected and conservative sites per domain in the parents and children omitting inferences with the over-estimation. As revealed by Fishers’ exact test, a significantly higher percentage of positively selected sites were found in the recombinants within the domains I and III when considering the transfer of the first domain as well as domain II of recombinants and domain III of parents for the events leading to the swapping of the third domain (fig. 5d, fig. S32c-d). However, differences in the proportion within the second domain were equal if event 23 with the extended positively selected region was not considered. Similar to site models, neither the amount of positively selected nor conservative sites were related to the number of all and marked toxins, which can be perceived as a further indication that the instances calculated reflect the evolutionary dynamics rather than the properties of subtrees chosen (fig. S33a-b). Linear models also exhibited that the respective abundance of both site types was significantly and positively associated with the foreground ω (fig. S33c-d).

Finally, we examined if and how evolutionary selection signals are linked with host specificity. First, we considered whether the parameters evaluated contribute to unequal sets of host orders between recombinants and parents. The inferred distributions provided some evidence, that the higher ω and amount of positive sites were associated with differences in host sets, while events with the same affected orders were enriched with conservative sites. However, the amount of data was not enough to gain significance except for the positively selected sites according to site models only (fig. S34a-e, tab. S52). We then considered the overlap coefficient in the context of host species, which sufficed to provide a finer picture. A significant negative correlation between the number of common species and ω, both background and foreground, as well as the number of positively selected sites was reported (fig. 5e-f, fig. S34f, tab. S52). In terms of branch-site models, no visible relationships were observed for positively selected sites, while the negative correlation between the number of conservative sites in the third domain and the similarity of host species was reported (fig. S34g-h).

The results obtained might be explained by higher purifying selection pressure once specificity is altered, which fits in with the previous observations. After the contact between host-restricted populations and the acquisition of specialized domains, a strong selection emerges to retain the received domains during the adaptation to novel hosts.

## Discussion

Recombination is considered one of the key evolutionary forces in the bacterial world. Apart from maintaining genomic diversity, this phenomenon increases environmental fitness thereby enabling organisms to thrive in novel niches during ecological diversification and the establishment of symbiotic/pathogenic relationships [50]. Although this phenomenon is mostly viewed as the replacement of the entire genes, recent studies show that intra-genic recombination causing domain swapping often occurs in loci encoding bacterial toxins, e.g., in *Clostridium botulinum* [51] or *Streptococcus salivarius* [52], being essential for the evolution of pathogenicity determinants (Supplementary text).

Shortly after sequencing the first *cry* gene, homologous recombination was proposed to take part in the evolution of Cry toxins [53]. With time, more evidence has been accumulated presuming several toxins, including Cry1Gb [43] and Cry2Aa17 [42], undergo swaps of domains I and III. The possible role of domain III swaps in expanding the natural diversity of Cry toxins was first explicitly stated by de Maagd et al. by comparing topologies of phylogenetic trees emanating from sequences of individual domains [30]. Notable examples were Cry1Cb clustering with Cry1Ca in terms of domains I and II, however, grouping with Cry1Eb and Cry1Be on the basis of the third domain [30]. We went on further with a full-scale study of the existing diversity of Cry toxins and revealed 50 recombination events, mostly referring to the third domain. Importantly, all the mentioned instances were identified by us as well (Supplementary text). Thus, we can conclude that not only our computational approach confirmed the existing examples of domain swaps but also sufficiently extended the list of chimeric toxins. In addition, there were several toxins in other species, namely, *Paenibacillus sp.*, *B. mycoides*, and *B. cereus* marked as parents in some events that were characterized either with larger amounts of toxins or domains of unknown origin (tab. S54).

We found recombination breakpoints surrounding the transferred regions exhibit substantial sequence similarity between parental sequences. In general, Cry toxins are known to contain several conservative blocks, which is a distinctive feature of Cry families. However, these short regions were viewed as structurally crucial regions modulating toxicity [23]. Most of the toxins pose 8 blocks with blocks 2 and 3 located between the boundaries of domains I/II and domains II/III, respectively. Moreover, some toxins have alternative and truncated blocks or are devoid of them except for blocks 1 and 2 in the Cry2 family. We showed that Cry2 toxins are involved in swaps of domain I exclusively. The presence of these conservative regions is quite useful for creating chimeric toxins [54] as sometimes they contain natural restriction sites [55]. The recombination process controlling swaps, at least in some cases, is RecA-independent [56]. Furthermore, computational prediction of thermodynamic patterns in *cry* sequences on the DNA level differed between phylogenetically distant groups causing domain swaps through shuffling among toxins with shared evolutionary history [57]. In this study, we focused on processed toxins leaving aside C-terminal parts. Nonetheless, domain-swapping experiments suggested that C-terminal regions enhance toxicity [40]. Moreover, C-terminal extension may as well be a consequence of homologous recombination [24]. A possible source for such exchanges and fusions could lie in partial sequences with C- and N-terminal regions [58]. Taking into account a multi-modular structure of C-terminal parts with up to five domains separated by conservative blocks, we can speculate that recombination and domain swaps might engage all functionally important distinct regions of Cry toxins.

Cry-encoding genes tend to reside in plasmids, however, they can also be located on chromosomes [59]; thus, exchanges of *cry*-carrying plasmids frequently occur in *Bt* isolates [60] with occasional inter-plasmid recombination [61]. The intensity of plasmid transfers seems to be restricted to adaptive clades [60]. Both chromosomal and plasmin *cry* genes are often flanked by IS (insertion sequences), especially those of the IS231 family [62,63], thought to induce recombination leading to the inclusion of virulence genes into chromosomes or between plasmids causing new combinations of toxins [59]. Genes coding for Cry1 proteins coupled with IS surrounding them form cassettes assumed to be the smallest mobile unit carrying insecticidal factors derived from the pathogenicity island disrupted by insertions [64]. We also revealed that assemblies carrying both parents and recombinants are surrounded by insertion sequences (IS) both in plasmids and chromosomes (fig. S23a).

It should be mentioned that collecting host specificity data illustrates that we still have a poor understanding of the spectrum of toxins’ activities apart from some Cry groups tested on multiple model objects belonging mainly to orders Diptera and Lepidoptera while others, both toxins and species, remain untested. For this reason, there is a strong demand for obtaining more high-quality complete genome sequences as well as mass screening of toxicity spectra on non-model organisms. In spite of these limitations, the usage of overlap coefficients as well as including/excluding well-studied toxins were sufficient for deciphering how recombination alters the host range.

The impact of domain exchanges on host specificity and general activity was first studied experimentally. Nevertheless, the ramifications for toxins ranged from enhancing activity to losing it. For instance, domain I exchange between Cry1A, Cry1C, and Cry1E sparked the formation of chimera non-toxic to *Spodoptera exigua* and *Plutella xylostella* [65], while the respective exchange between Cry9Aa and Cry1Ac, contrarily, increased the activity against *Helicoverpa armigera* fivefold [66]. The transfer of the third domain from Cry1Gc to Cry1Ab magnified toxicity to *Chilo suppressalis* [67]. This technique applied to Cry1Ac and Cry1Ca made a hybrid toxic to *Spodoptera frugiperda* unlike non-lethal parents [68]. Natural swaps described by us also do not adhere to a universal scheme with the host spectrum of recombinants being identical to parents or diverging sufficiently. The same activity towards *S. exigua* was shown to Cry71Aa and its donor of the third domain, Cry72Aa (tab. S36). The swap of domain I from Cry14Ab1_85.6_AXE15628.1 to Cry14Ab increased toxicity towards *Caenorhabditis elegans* twofold compared to major parent Cry14Aa [69]. Cry1Fa1, unlike its domain III parents, Cry1Ka2 or Cry1Jd1, and major parent Cry1Jb exerts an effect on *Bombyx mori* and *S. exigua* [47]. In the same manner, receiving the third domain by Cry1Cb gained toxicity against hemipteran together with lepidopteran insects, to which parents are toxic only [70]. Parasporins Cry41Ab and Cry41Aa also appeared to have undergone recombination with the former acquiring domain III from Cry32Ma1_57.7_AXY11484.1. Noteworthy, Cry73Ba which is close to the Cry41 family and kills hemipteran *Myzus persicae* [71] was related to the toxins of the Cry32 family of unknown specificity (tab. S37). Members of this abundant Cry class were reported to be toxic against Lepidoptera, Diptera, and Nematoda, and they belonged to 5 recombination events implying that this Cry group is highly promising and deserving further investigations. Another prospective Cry family is Cry7 forming a distinct connected component in a host-wise strain graph (fig. S25). While the host spectrum of these toxins remains unknown, they were reported to get certain domains Cry7Ca1, the only known protein to act against *Locusta migratoria manilensis* Orthoptera [72]. Therefore, the Cry7 family on the whole deserves testing for anti-orthopteran activity as a prominent source of possible biocontrol agents.

According to current genomic studies, recombination often comes with purifying selection during the co-evolution of pathogens and symbionts. For instance, many rounds of recombination events in major antigenic protein 1 (map1) family genes of pathogenic *Ehrlichia ruminantium* led to new loci, the most successful of which were kept by purifying selection [73]. The same observations were made in the genomes of symbionts *Rhizobium* [74] and *Verminephrobacter* [75]. To date, there is only one research item dedicated to evolutionary selection in Cry toxins. Using the maximum likelihood method applied on 18 Cry sequences, 24 positively selected sites located in domains II and III were identified [29]. Given the role of these domains, it was proposed that positive selection might induce overcoming host resistance. However, the set of toxins chosen was quite arbitrary. The clade with the toxins mentioned consists of 115 toxins according to our phylogeny built on all three domains. In our approach, we evaluated selection strength in the context of recombination by selecting clades encompassing chimeric toxins with their parents. Only the site models provided equivalent findings with a plethora of positively selected sites concentrated in the central regions of the domains (fig. 5c). The usage of branch and branch-site models, in their turn, revealed noticeable purifying selection, and the prevalence of conservative sites (fig. 5a, fig. S32c-d). Importantly, the more distant toxins are in terms of their host specificity, the higher selective pressure exists (fig. 5d-e, fig. S34f-h). We hence can hypothesize that positively selected sites reflect the general route of adaptation to mutations in hosts’ receptors to overcome resistance, while purifying selection implies that recent acquisition of functional domains provides a considerable advantage.

## Conclusion

To sum up, all the data gathered through an arsenal of bioinformatic techniques applied allowed us to explicitly demonstrate that recombination is one of the main mechanisms driving the evolution of the Cry toxins with only recent events affecting more than half of the known sequences. Most of the breakpoints are within close proximity of the toxins’ domain borders falling into relatively conservative flanking regions. Therefore, we proposed that these regions facilitate the domain changes thereby sparking the emergence of novel Cry toxins with diverse combinations of domains. As revealed by the analysis of the host specificity spectrum, acquiring these newly emerged toxins enables bacteria to adapt to new host species extending their ecological niches. This result is confirmed by the patterns of positive and purifying selection suggesting the adaptive process while interacting with new insect receptors. Overall, our research revealed the primary role of recombination in orchestrating the evolution of Cry toxins, what mechanisms govern recombination events, and what effects they exert on host specificity. The results provide insights into which domain compositions might extend the affected host range thus being useful for constructing new variants of artificial toxins with targeted specificity.

## Methods

### Data processing and visualization

All data-processing steps were performed using custom scripts written on Python v3.6.9. Statistical tests were carried out using the R v.3.6.3 programming language. Plots, phylogenetic trees, and graphs were visualized using libraries ggplot2 v3.3.5 [76], ggtree v1.16.6 [77], and pygraphviz v1.6 [78], respectively. The code used as well as the raw data is available at the git-repository: https://github.com/lab7arriam/Cry_phylogeny.

### Data acquisition

Four datasets were the sources of 3D-Cry toxins. These included protein sequences from the NCBI IPG database (Identical Protein Group, *Bacillus thuringiensis* taxonomy ID: 1428, accessed on the 3rd June 2020 (the same for all datasets), 467 *Bt* assemblies from NCBI Assembly, BPPRC (Bacterial Pesticidal Protein Resource Center) [45], and NCBI Genbank (bacteria non-redundant). Three-domain Cry toxins were extracted with CryProcessor in the “-*do”* mode [5]. We used CD-HIT v4.8.1-2019-0228 [79] for deduplication specifying the following parameters: word size of 5 and 95% identity threshold. We then excluded artificially modified toxins based on IPG annotations and characterized the general properties of the nucleotide sequences (Supplementary text).

### Phylogeny reconstruction

To reconstruct phylogenetic trees, we aligned sequences of deduplicated reference clusters with the MAFFT aligner v7.429 [80] and trimmed the alignments trimAl v1.4.rev22 utility [81]. Phylogenetic trees were built with RAxML-NG v1.1.0 [82] based on evolutionary models revealed by ModelTest-NG v0.1.7 [83]. For detailed parameters, see the Supplementary text. We then compared the topologies of the reconstructed trees with the quartet distance metrics using the tqDist v1.0.2 software [84], Robinson-Foulds distance implemented in RAxML-NG [82], and cophenetic distance by the ape v5.6-2 package [85]. Tree quality metrics, namely consistency index (CI) and retention index (RI) were calculated with phangorn v2.5.5 [86], and tree balance indices such as ‘CollesLike’, ‘Sackin’, and ‘Cophenetic’ were identified using the CollessLike v2.0 package [87]. A graphical display of topological discrepancies was illustrated as tanglegrams produced by the dendextend v1.14.0 package [88]. We also plotted topological dissimilarity in a heatmap-wise way: the order of the toxins corresponded to the full core toxin-derived tree with the values depicting the pair-wise identity of a certain domain.

### Detecting recombination events

We utilized the RDP4 v101 (recombination detection protocol) utility for predicting individual recombination events [89]. We also used three other tools such as fastGEAR [90], Gubbins v3.1.2 [91], and ClonalFrameML v1.12 [92] to calculate the recombination to mutation ratio (r/m). We then discarded events using a three-step filtration based on the RDP output, lengths of the transmitted region, and congruence of parents and children on domain-wise trees (Supplementary text). We then evaluated the quality of the datasets by comparing identities between domains of parents and recombinants and characterizing the mismatch ratio (Supplementary text).

Afterward, we employed two approaches to plot the map of recombination exchanges. First, we built a Circos plot using the circlize v0.4.13 [93] package to display each transfer between toxins arranged according to the reference tree. Second, we created the unweighted directed recombination graph with a custom Python script implementing networkX v2.4 [94] library and revealed connected components within it. Finally, we assessed the number of toxins possibly absent in our dataset based on the proportions of recombination-affected toxins in trees based on the individual domains (Supplementary text).

### Predicting possible mechanisms of domain exchanges

To access the overall recombination rate we reconstructed pangenomes using Panaroo v1.2.8 [95] of genome assemblies grouped according to the presence of *cry* genes and toxins subjected to domain swaps (Supplementary text). Estimates of the r/m ratio based on concatenated core gene alignments were obtained with the ClonalFrameML v1.12 program. Insertion sequences (IS) were detected with ISEScan v1.7.2.3 [96]. Prohages were predicted by Phigaro v2.3.0 [97], and genetic islands (GI) were found using IslandPath-DIMOB v1.0.6 [98] utility. We then compared the abundance of MGEs in the studied assemblies (Supplementary text).

To identify conservative blocks, we took positions of breakpoints in the parental sequences from the RDP4 output and aligned sequences of parents within the events with subsequent excision of 100 b.p. regions downstream and upstream of the breakpoints. Two types of these flanks were selected, namely, located between the boundaries of the first and the second domains, and the second and the third domains, respectively. We then calculated site-wise identity between parental sequences and ran the rollmean function from R package zoo v1.8-9 [99] with center alignment and a window size of 7 accordingly. For clustering these blocks, we chose an optimal clustering pipeline using silhouette analysis implemented in the factoextra v1.0.7 [100] package and utilized it accordingly (Supplementary text).

### Studying the impact of recombination on host specificity

To determine the influence of recombination on host specificity and toxins’ composition, we first collected the metadata from the IPG database and attributed the toxins to strains they are present in. We then gathered the spectrum of activities from BPPRC and exhaustive reviews [47–49] and manually searched for research articles and patents to ascertain the hosts infected by the toxins as well. The information obtained was attributed to toxins, both on the basis of fourth and third ranks in the established nomenclature, and strains in case individual proteins were not tested. The percentage of orders to which individual species pertained were displayed as event-wise scatterplots via the scatterpie v0.1.8 [101] package. We then compared both the sets of hosts affected by recombinants and parents as well as the compositions of toxins in strains containing the respective participants of recombination using different similarity metrics (Supplementary text). We also picked assemblies in which at least one recombinant and parent for the particular event co-occurred and visualized the genomic localization of MGEs and regular loci using the Proksee [102] web-based service. With this data, we constructed a weighted graph using networkX v2.4 [94] library with the nodes corresponding to strains with the edges containing the list of toxins shared by the pair of strains, and the weight indicated the number of common toxins. Afterward, we figured out which connected components harbored the parents and the recombinants and compared the sets of hosts using both Jaccard and Szymkiewicz–Simpson coefficients.

### Analyzing evolutionary selection

Due to the high computational resources required for evaluating evolutionary selection, we developed a multi-step pipeline to prepare the data for such calculations (Supplementary text, fig. S35b). Following that, we determined the optimal evolutionary model using ModelTest-NG v0.1.7 [83] and prepared the guiding tree with RaxML-NG v1.1.0 [82]. Owing to the fact that assessing evolutionary selection operates with codons, we aligned protein sequences with MAFFT v7.429 [80] and matched the nucleotide sequences to protein alignments with a custom Python script. We then examined three types of models using the ete3 package (fig. S35c). Second, we applied site models, bneut, and btree, indicating neutral selection and positive/negative selection, respectively, marking leaves harboring either recombinants or parents. For detecting alterations in selection patterns over branches, we opted for the models exhibiting significantly different LRT tests when comparing M0 with b_free, and b_free with b_neut models, implying different ratios on branches and that the evolution in the foreground branch is not neutral, respectively. Inferences on account of the non-synonymous to synonymous substitution rate ratio (ω = dN/dS) value were marked as being subjected to positive selection if the ω value on the foreground exceeded 1 and to purifying selection otherwise. Second, different site models were examined, namely, M0, M1, M2, M3, M7, and M8. The sites, both conservative and positively selected, were aggregated for the best-fit models determined by the likelihood. Third, we considered branch-sited models, namely, BsA, BsA1, and BsB, and summarized both sites and branch-based ω values for the events with significantly detected selection. To find relationships between different factors, including the number of toxins, ω on foreground and background branches as well as the number of conservative and positively selected sites we deployed linear regression. We also studied whether these factors delineate the sets of hosts using the same method accordingly.

## Acknowledgments

We express our gratitude to Oksana Belousova for English editing.

