## Supplemental figures for "Recombine and succeed: a story of Cry toxins to expand the host range"

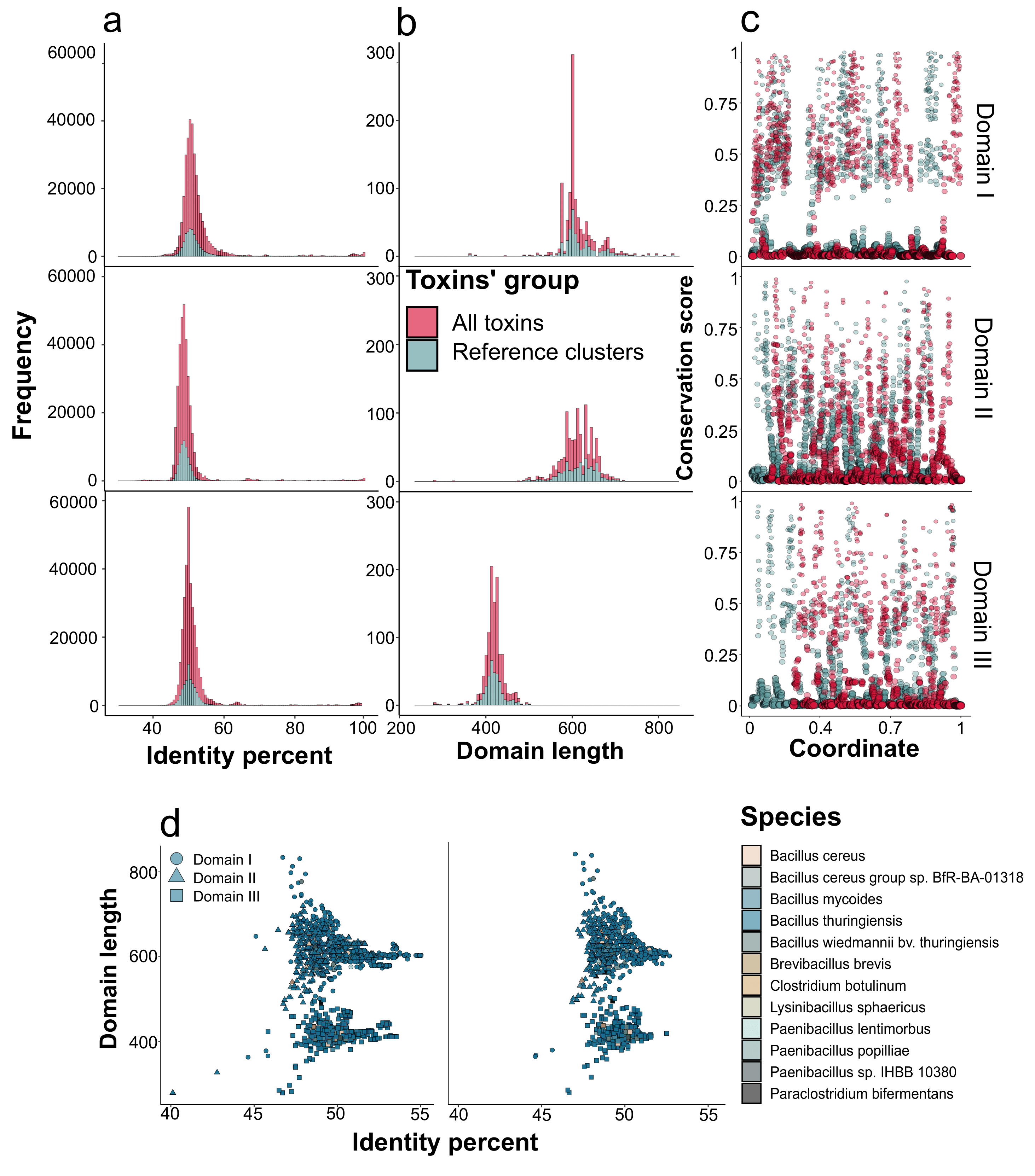

**Fig. S1**. Properties of Cry toxin sequences identified in four publically available datasets, namely, NCBI Assembly [1], Genebank, IPG (identical protein groups), and the BPPRC (Bacterial Pesticidal Protein Resource Center) [2]. (**a**) The total frequency of domain-wise similarity between all Cry toxins’ sequences used in the study. The color code depicts the type of the dataset, namely, all toxins and reference clusters obtained using CD-HIT with a 95% identity threshold. (**b**) Distribution of the domains’ lengths in the analyzed dataset. (**c**) Per-site conservation scores in the sequences of the domains calculated as the maximum frequency of non-gap symbols in the alignment of the particular domain. The first domain has a higher conservation score with a mean of 0.23 in comparison with the second and third domains (0.16). The number of highly conserved sites with conservation scores exceeding 0.85 reached 16, and 41 conserved sites for the first domain, the second, and the third domains, respectively. (**d**) The dependence between the length of the domain and mean pair-wise identity for the respective toxin with all toxins from the dataset. The color corresponds to the species’ attributions, while the shape encodes the domain type.

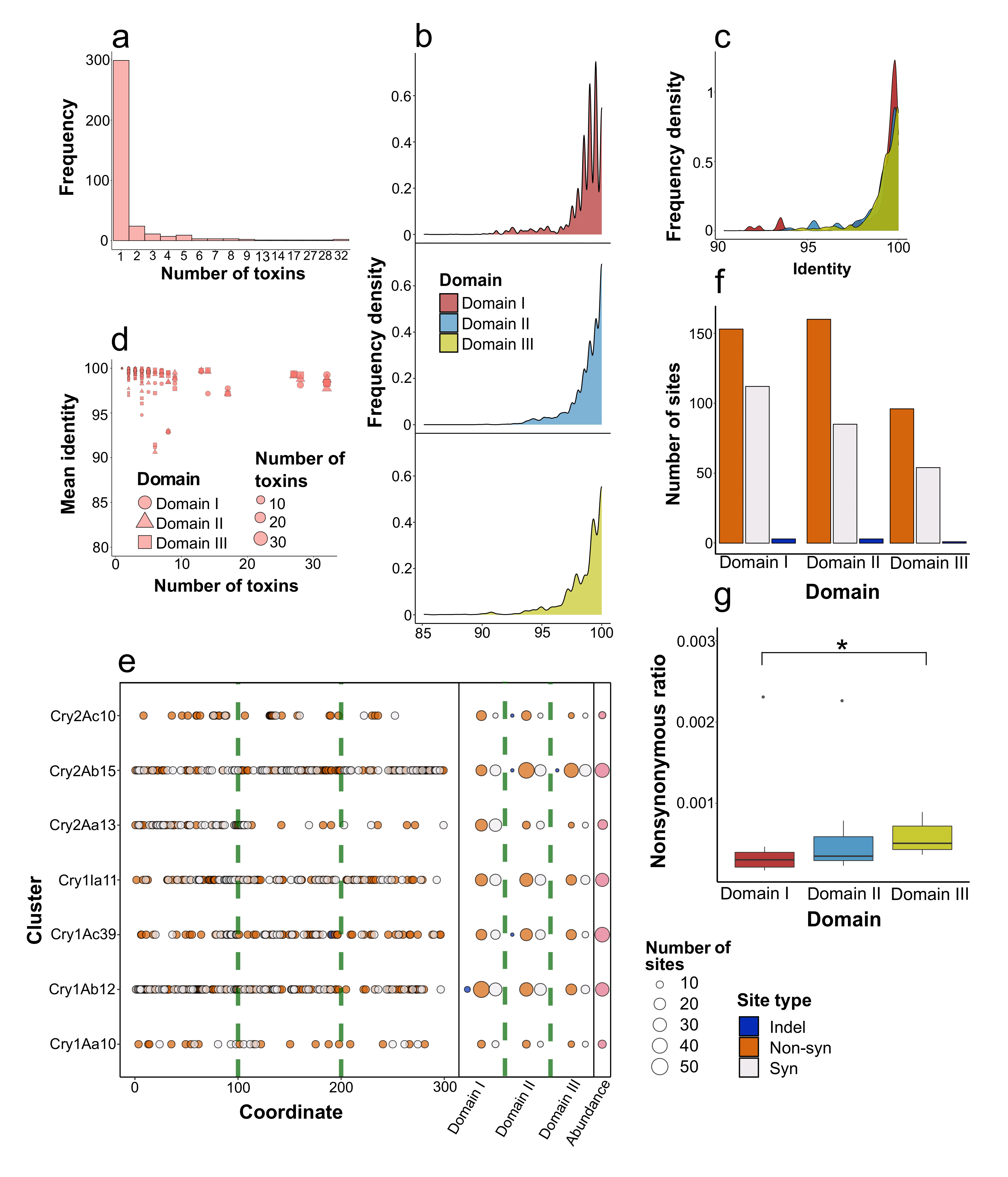

**Fig. S2**. Characteristics of cd-hit clusters of Cry toxins obtained using a 95% identity threshold. (**a**) Frequency of clusters according to the number of toxins within them. (**b**) Mean pairwise similarity of the domains’ sequences within all cd-hit clusters obtained using a 95% identity threshold. (**c**) The same distribution of domain-wise identities in clusters containing more than 10 toxins. (**d**) Dependence between the abundance of the cluster and mean domain-wise identity between toxins within the cluster. The shape of points denotes the domain, while the size encodes the number of toxins. (**e**) The distribution of pairwise identities of the domains within clusters encompassing more than 10 toxins. Codon-wise mismatches between the sequences of the domains within clusters with more than 10 toxins. The left panel shows the coordinates of mismatches colored according to their type (synonymous, non-synonymous, and indels) with a green dashed line designating the boundaries between the domains. The middle panel summarizes the number of such mismatches per cluster, while the right panel denotes the abundance of the cluster. (**f**) The total number of substitutions identified in the particular domain. (**g**) The normalized non-synonymous ratio for the particular domain calculated as the sum of non-synonymous substitutions and indels divided by the number of synonymous mismatches multiplied by the length of the cluster-wise domain-based alignment and the number of toxins within the cluster. Asterisks denote statistical significance using the Wilcox test when comparing the respective values of domains’ sequences.

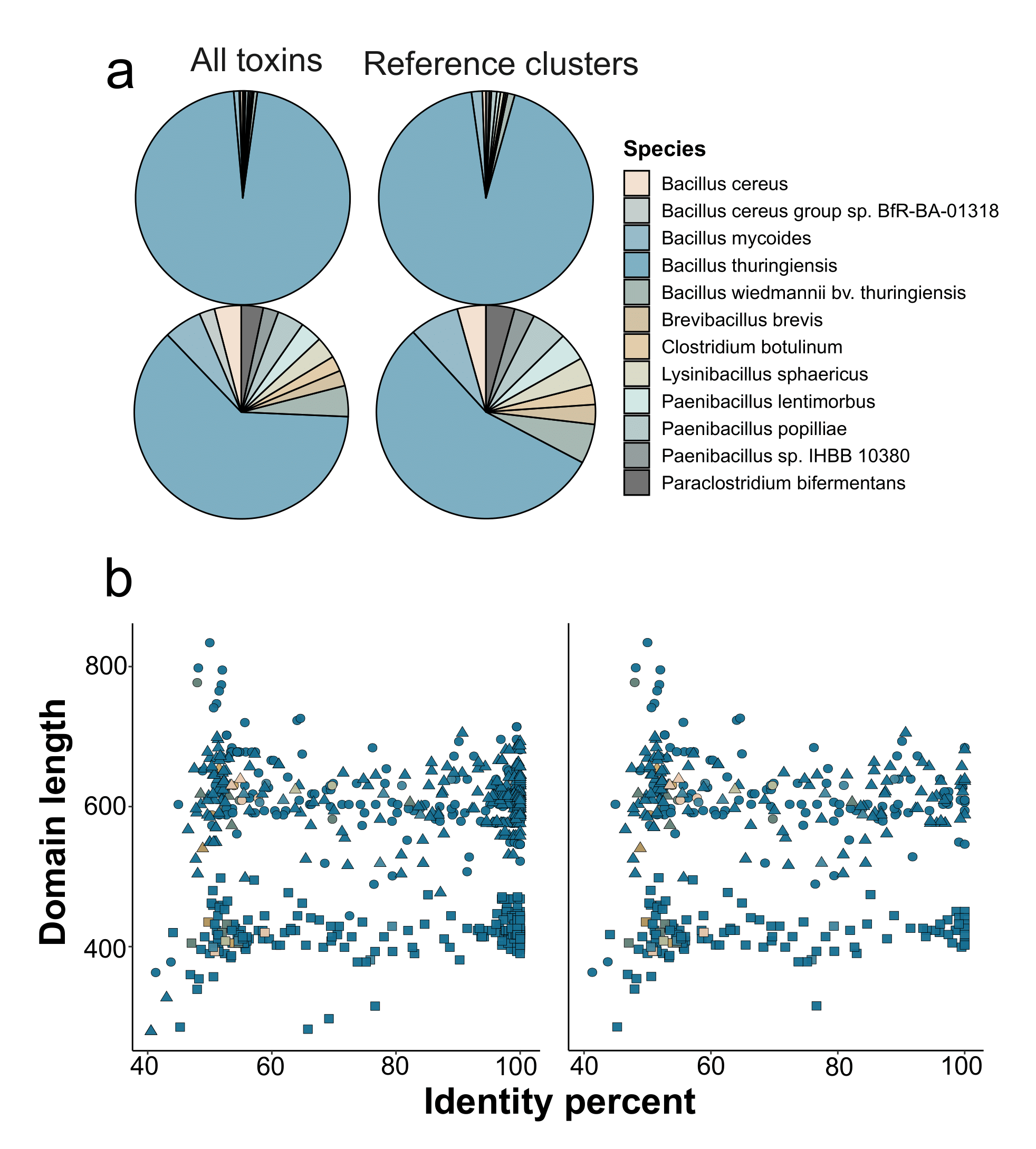

**Fig. S3**. Properties of Cry sequences of the analyzed dataset with respect to species in which toxins were identified and the identity with the respective known homologs. (**a**) Distribution of species’ attributions of 3D-Cry toxins dataset. Shown are the data for all toxins and reference clusters obtained using cd-hit with a 95% identity threshold. Upper plots display the absolute numbers, while the lower plots depict square root-adjusted values for better details. A total of 30 toxins were attributed to other species, including *Bacillus mycoides*, *B. cereus*, *B. wiedmannii*, etc., or pertained to different genera, such as *Paraclostridium bifermentans* and *Clostridium botulinum*. (**b**) The relationship between the length of the domains and the identity with the respective homolog from the *Bt* nomenclature for all toxins and only novel ones. The color corresponds to the species’ attributions, while the shape encodes the domain type.

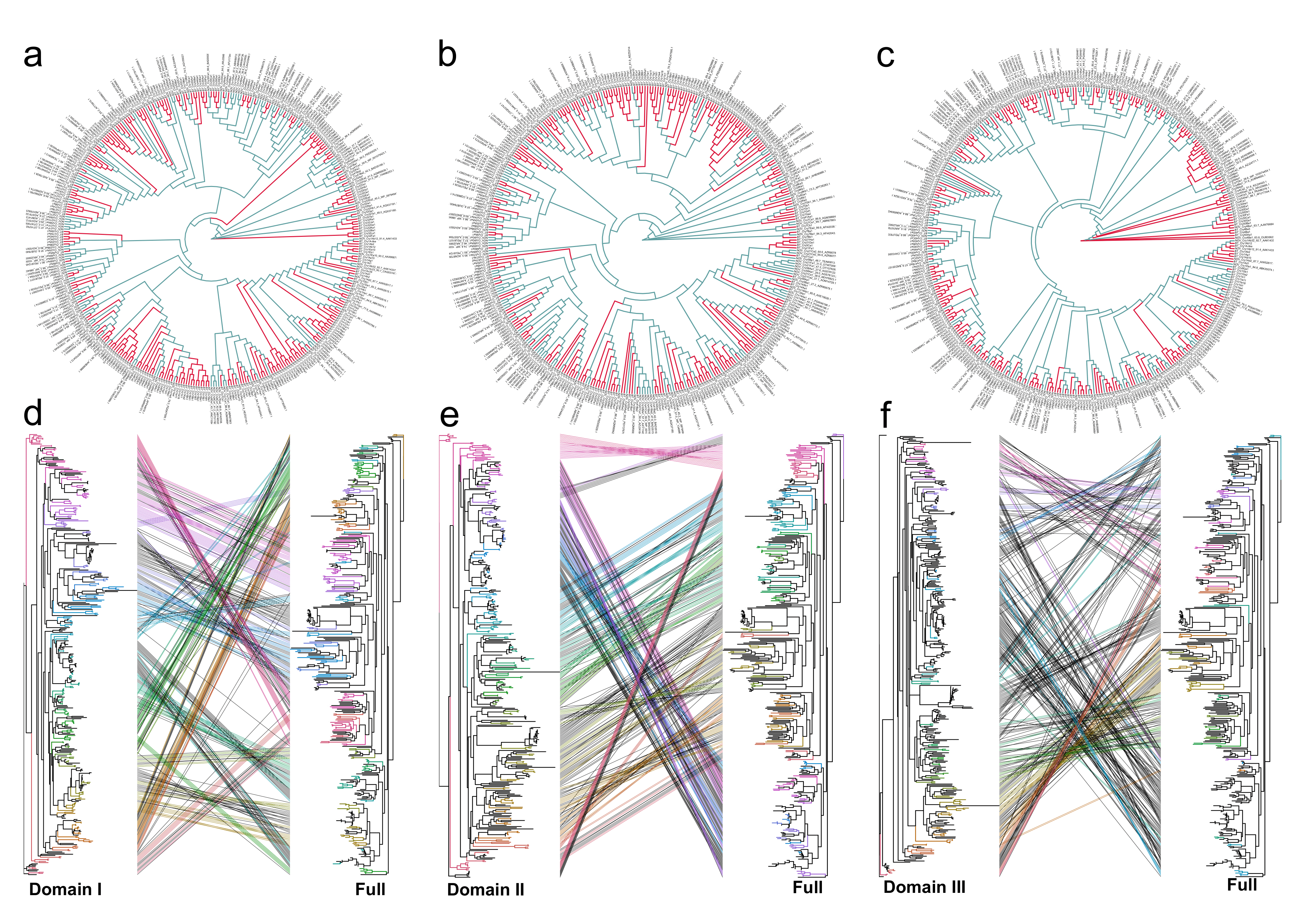

**Fig. S4**. Phylogenetic inferences based on the sequences of the domains in the analyzed dataset. The upper figures show reconstructed ML trees (maximum likelihood) obtained from the alignments of the first (**a**), second (**b**), and third (**c**) domains. Color code denotes the type of toxins with the red color corresponding to known toxins deposited in the *Bt* nomenclature and the blue color designating novel proteins. The lower figures depict tanglegrams demonstrating topological differences of reference ML phylogeny based on the concatenated alignments of the domains with partitioned evolutionary models with the respective trees of the first (**d**), second (**e**), and third domains (**f**). The optimal evolutionary model for the third domain differed from the other two domains (HKY+G4 vs. TVM+G4).

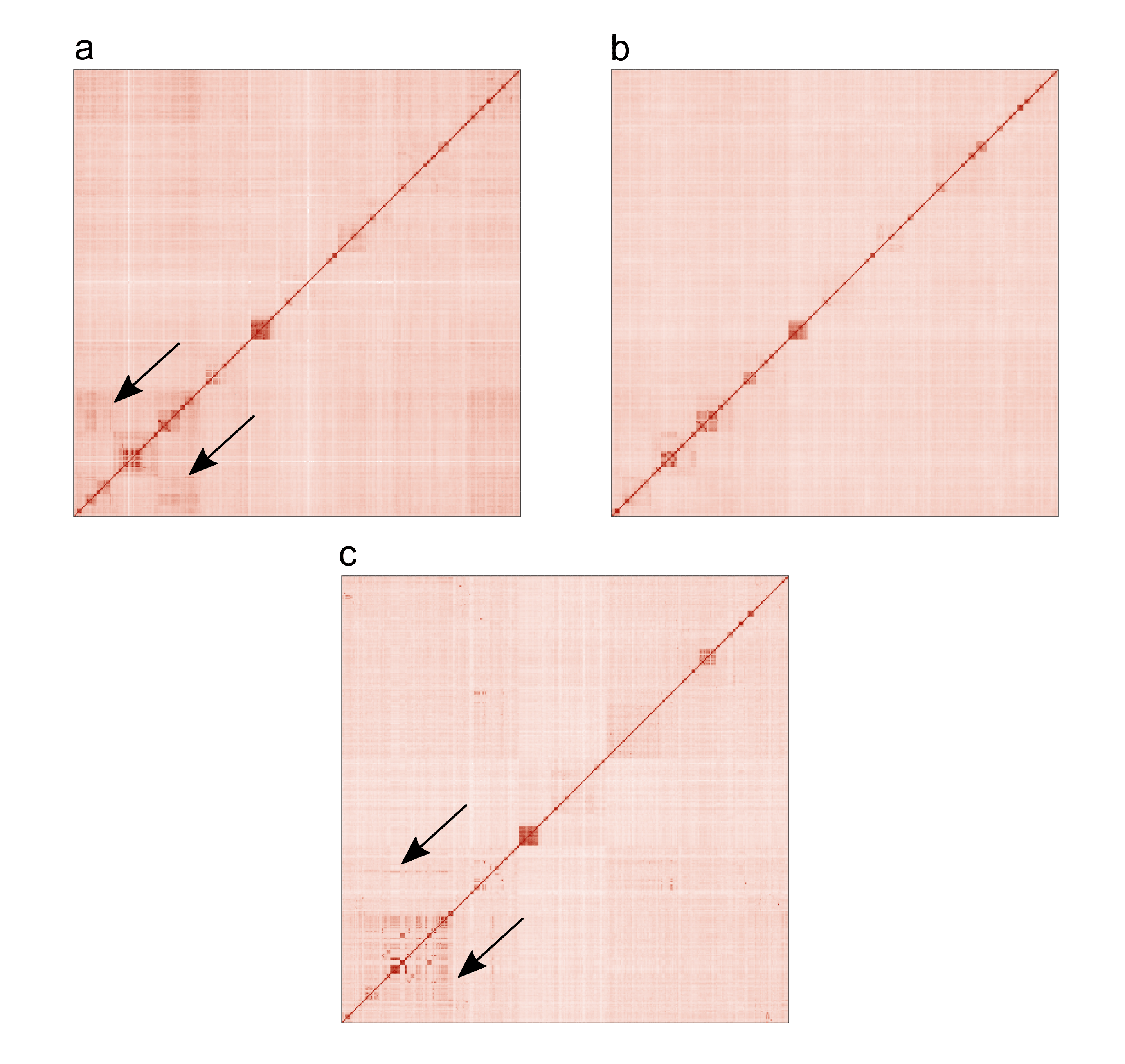

**Fig. S5**. Heatmap with the pair-wise identity of sequences of the first (**a**), second (**b**), and third (**c**) domains. The order of rows and columns corresponds to the order in the reference ML phylogeny based on the concatenated alignments of the domains with a partitioned evolutionary model. Arrows indicate the zones in which local identity differs from the expected order.

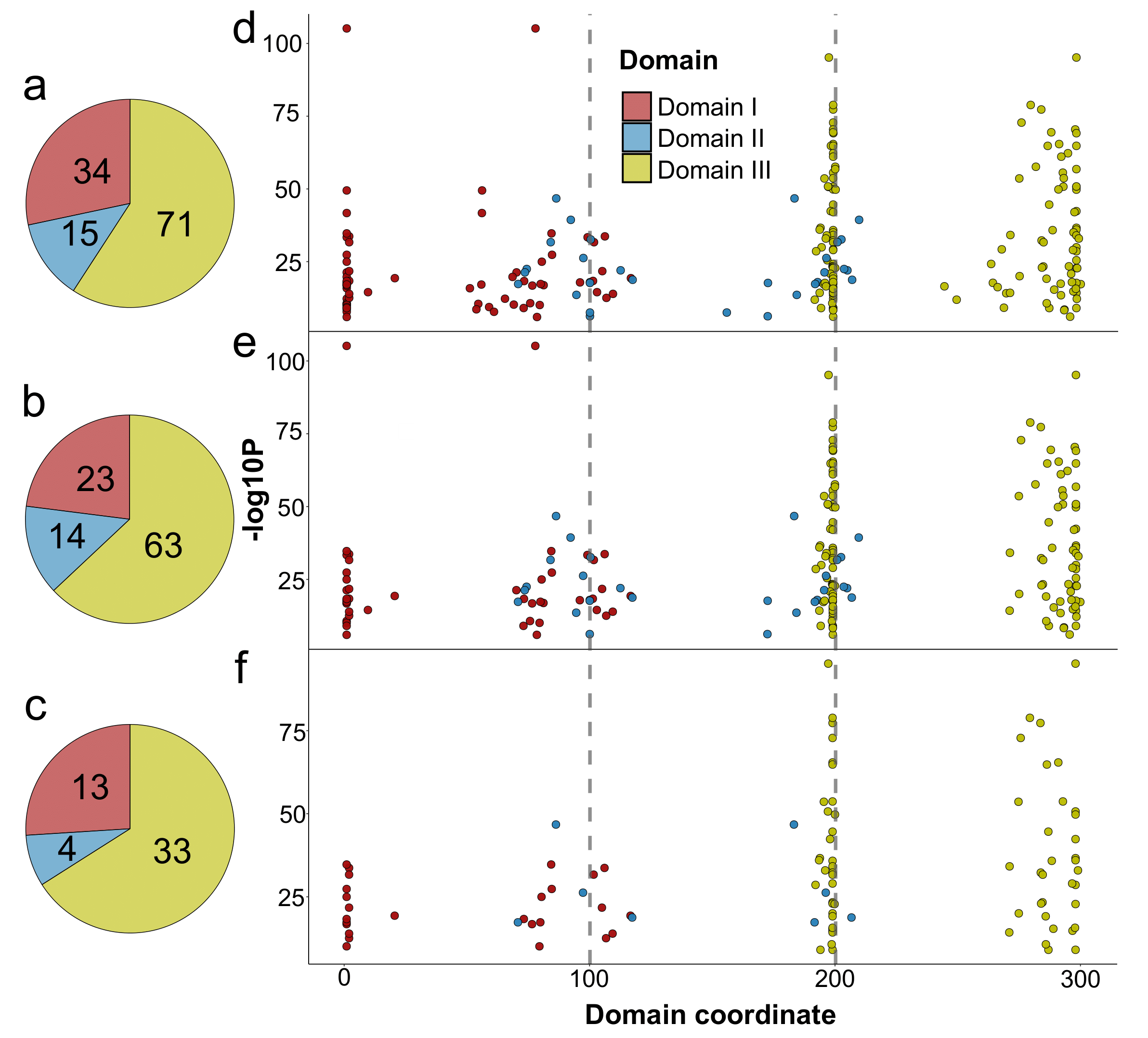

**Fig. S6**. Recombination events reported by RDP4 launched on MAFFT-aligned nucleotide sequences of 3-D Cry toxins. (**a**) The total number of raw events identified with RDP4. (**b**) The number of events after discarding partial exchanges, i.e., affecting less than 70% of the domains’ lengths (**c**) Types of events in the final dataset with filtering based on tree congruence. The color corresponds to the transferred domain. (**d**) Distribution of breakpoint coordinates for recombination events, as well as, those devoid of partial exchanges (**e**), and filtered on the basis of tree congruence (**f**). On the x-axis, the relative coordinate in the sequence is presented. Plotted on the y-axis, is the mean p-value from all recombination detection tests included in the RPD4 software. Grey dotted lines represent the boundaries of the domains. For a detailed description of the filtration process, see Fig. S34a.

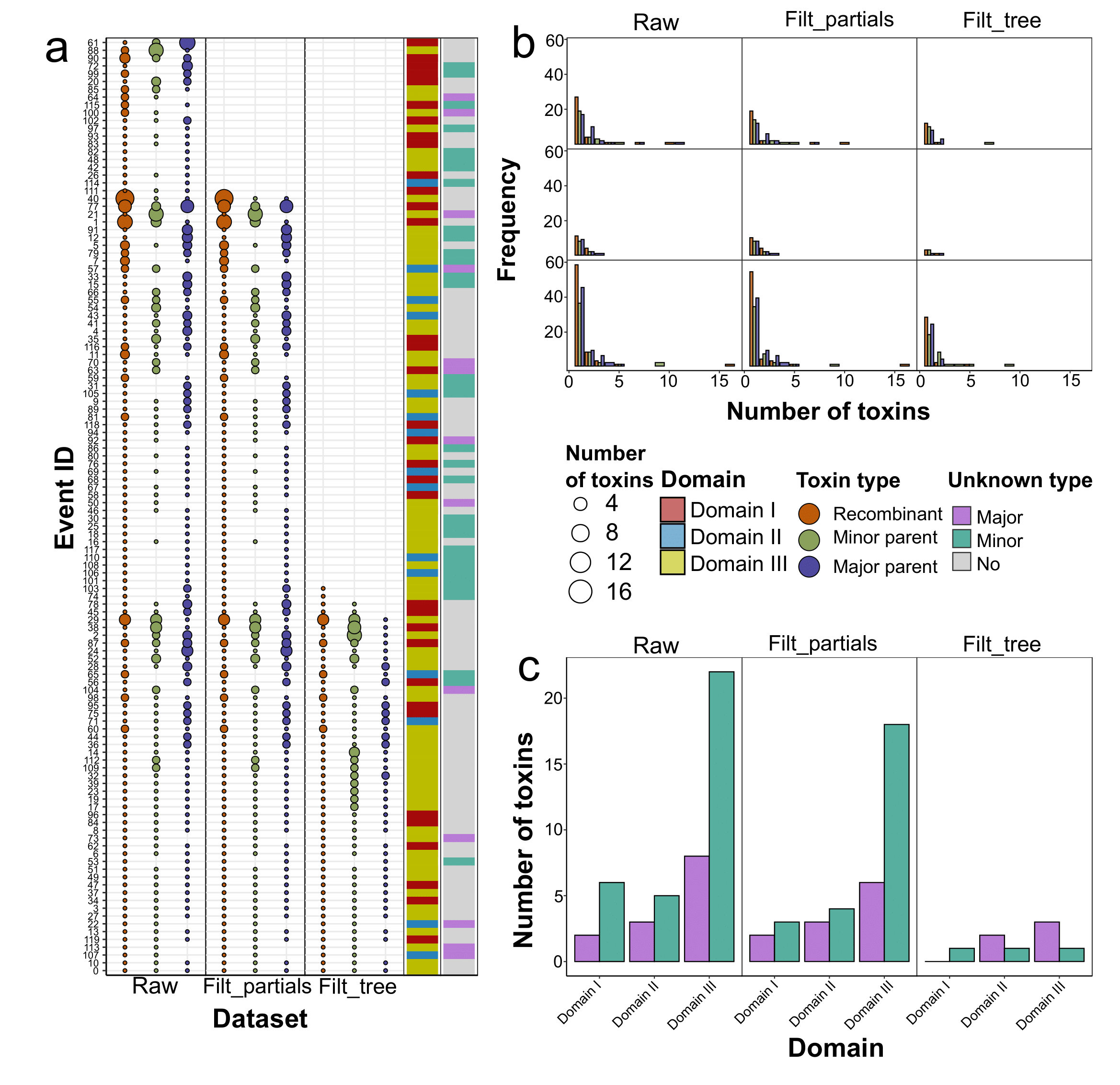

**Fig. S7**. Properties of recombination events detected with RDP4 launched on MAFFT-aligned nucleotide sequences of 3-D Cry toxins. Tree datasets are considered: “*Raw*” index encodes all events detected by RDP4, “*Filt_partials*” denotes the set of events affecting more than 70% of the respective domain, whereas “*Filt_tree*” marks the final dataset with filtering on the basis of tree congruence. For a detailed description of the filtration process, see Fig. S34a. (**a**) Event-wise characteristic of recombination exchanges within three datasets. Each point represents the set of toxins colored according to their type in the event. i.e., recombinants, and parents. Parents are marked as major or minor with the latter providing two domains, and the former transferring one domain. The size of the point is proportional to the number of toxins. The first adjacent line encodes the type of the event (the transferred domain), while the second line provides information about whether the parent is unknown, i.e., absent in the analyzed dataset. (**b**) Distribution of the number of toxins within the events. The color of the bar represents the type of recombination participants. Plotted on the x-axis is the number of toxins, while the y-axis denotes the number of events. In the final dataset, 45 events included one recombinant per event, while 4 events encompassed two toxins, and 1 event comprised 5 participants. The same goes for parents. Of all events, 31 were those with a single minor and major parent, 13 possessed either two or one major or minor parents (with 3 parents in total), and the rest of the events (6) were related to a greater number of parents. (**c**) The number of unknown parents within the recombination events. The total number of events grouped according to the transferred domain is presented. Color encodes the type of the parent (minor/major). The reduction in the number of events with unknown parents was from 46 (38%) to 8 (16%). In the final set, 5 major and 3 minor unknown parents remained.

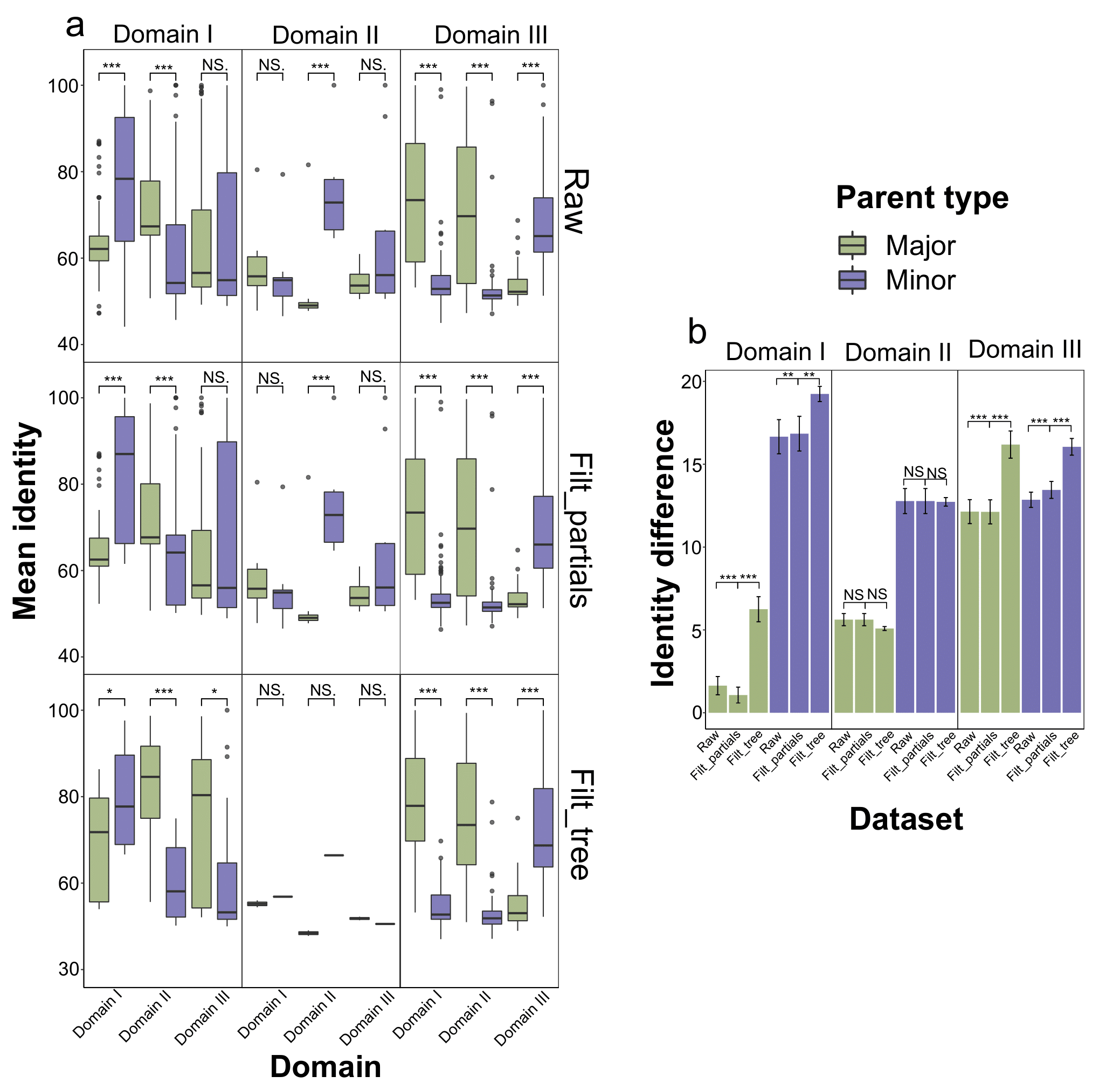

**Fig. S8**. Distribution of mean sequence identity and mismatch rates of recombination events detected with RDP4 launched on MAFFT-aligned nucleotide sequences of 3-D Cry toxins. Tree datasets are considered: “*Raw*” index encodes all events detected by RDP4, “*Filt_partials*” denotes the set of events affecting more than 70% of the respective domain, whereas “*Filt_tree*” marks the final dataset with filtering on the basis of tree congruence. For a detailed description of the filtration process, see Fig. S34a. All the measurements refer to three groups of recombination events according to the swapped domain. (**a**) Average domain-wise nucleotide identity between the sequences of recombinants and parents. Asterisks denote statistical significance using the Wilcox test when comparing the respective values of major and minor parents. The mean similarity between the transferred domain of recombinants and minor parents reached 80%, 66%, and 71% for events in which the first, the second, and the third domains were swapped, respectively. (**b**) The dataset-wise mean identity difference between thetransferred and non-transferred domains of major and minor parents compared with the domain sequences of recombinant toxins. Asterisks denote statistical significance using the Wilcox test when comparing three sets of recombination events. Error bars represent mean square error. Difference estimates reached 19%, 12%, and 16% when considering minor parents, and 6%, 5%, and 16% concerning majors for events with the first, second, and third domains, respectively.

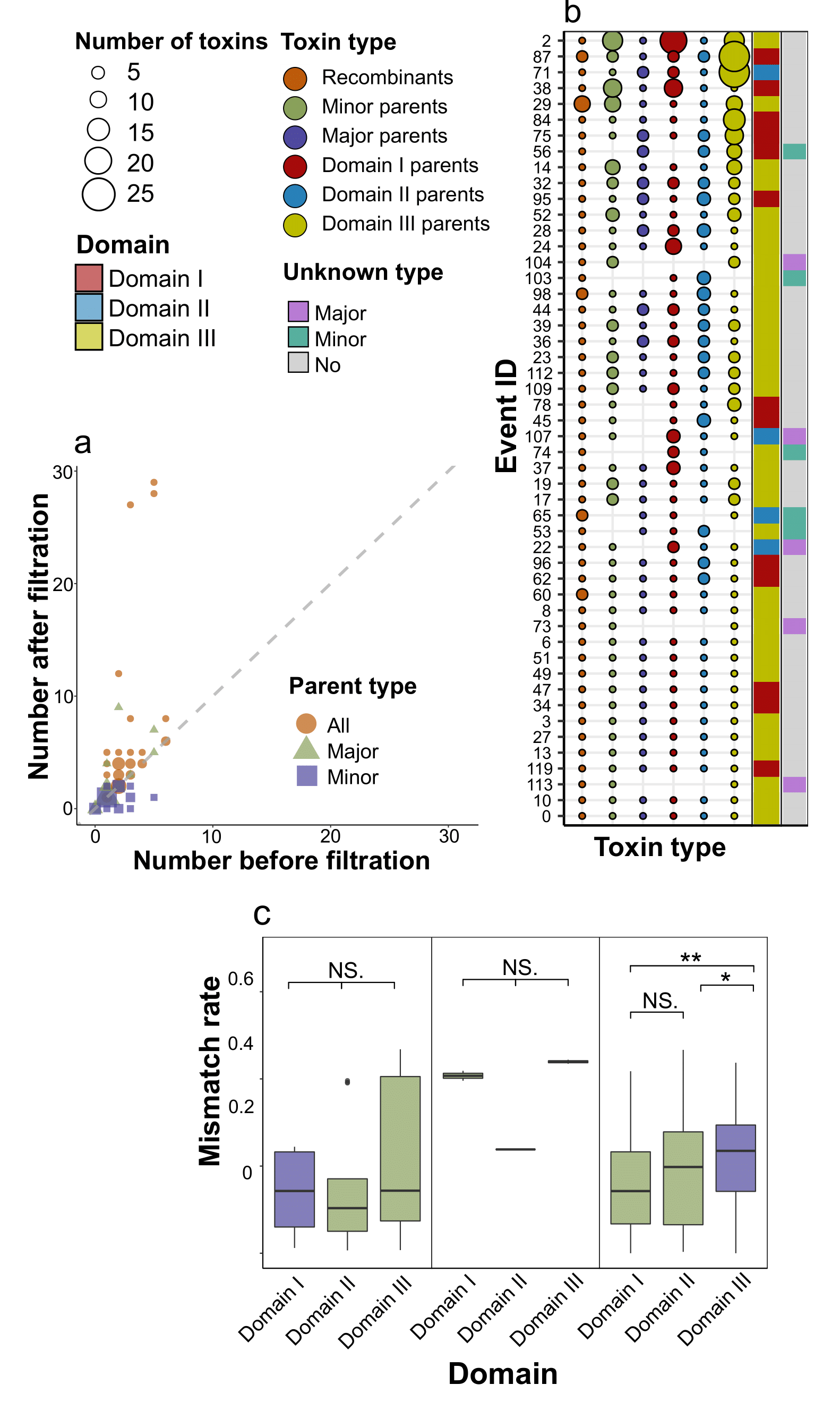

**Fig. S9**. Properties of the final dataset of recombination events detected with RDP4 launched on MAFFT-aligned nucleotide sequences of 3-D Cry toxins. For a detailed description of the filtration process, see Fig. S34a. (**a**) Event-wise characteristic of recombination exchanges. Each point represents the set of toxins colored according to their type in the event. i.e., recombinants, and parents. Parents are marked as major or minor with the latter providing two domains, and the former transferring one domain. Parents identified when considering domain-wise trees for each individual domain are also shown. The size of the point is proportional to the number of toxins. The first adjacent line encodes the type of the event (the transferred domain), while the second line provides information about whether the parent is unknown, i.e., absent in the analyzed dataset. (**b**) The effect of tree-wise filtration on the number of parents within recombination events. Plotted on the x-axis is the number of toxins reported by RDP, while the y-axis represents the number of parents after filtration. The shape and the color of points correspond to the type of parents. A total of 35 events remained unchanged, while for 8 events the number of majors increased, and for 7 events the number of minors was reduced. (**c**). Domain-wise number of mismatches between parents and recombinants normalized by the corresponding alignment. Mismatches were calculated only for transferred domains within the final filtered set of events. Asterisks denote statistical significance using the Wilcox test between domain-wise mismatch rates.

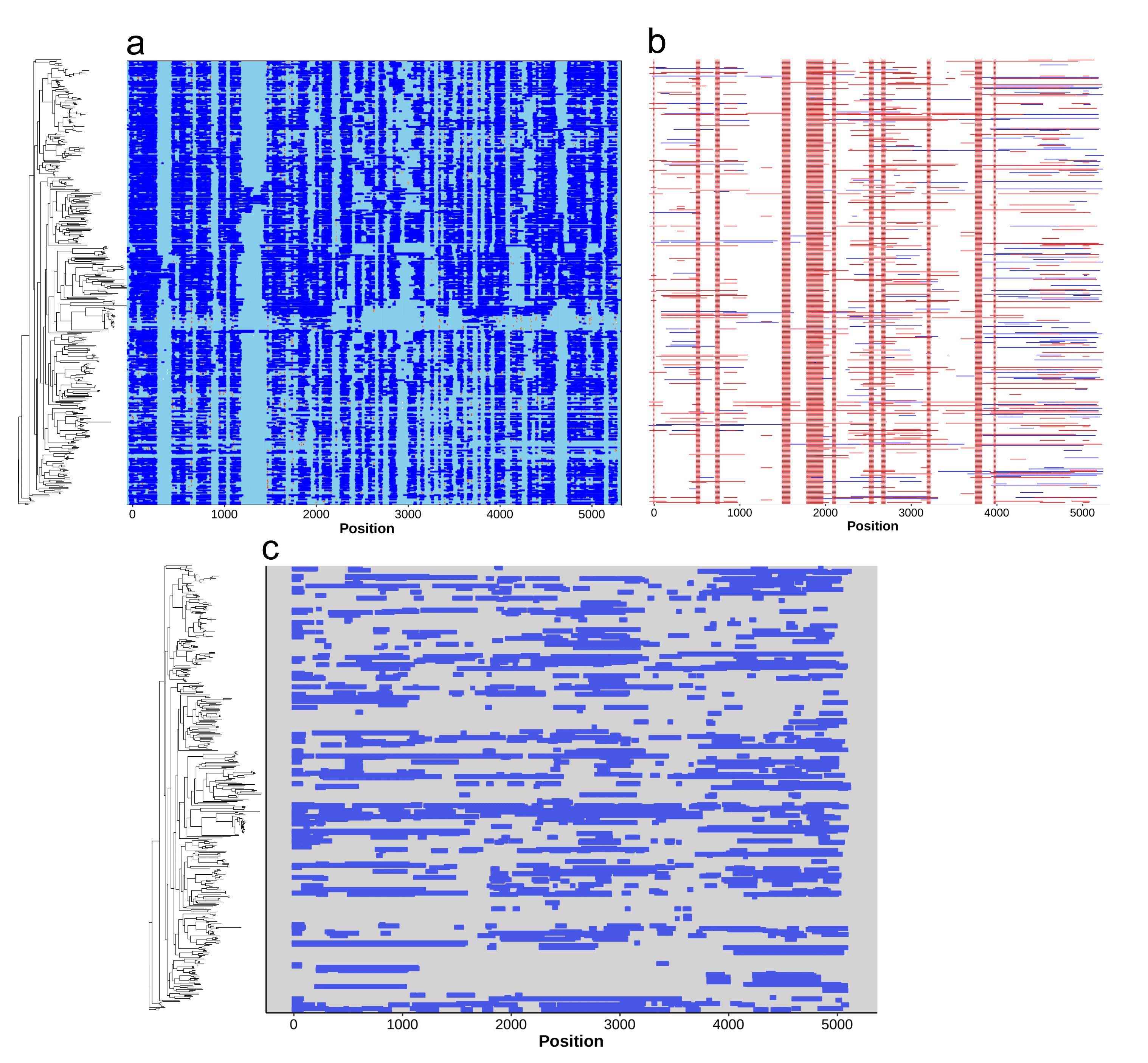

**Fig. S10**. Distribution of regions affected by recombination using ClonalFrameML (**a**), Gubbings (**b**), and fastGEAR (**c**). The order of rows corresponds to the reference phylogeny, and the positions refer to the MAFFT-generated alignment of processed toxins (regions from the beginning of the first and the end of the third domain). Putative exchanged areas are marked with blue and red colors.

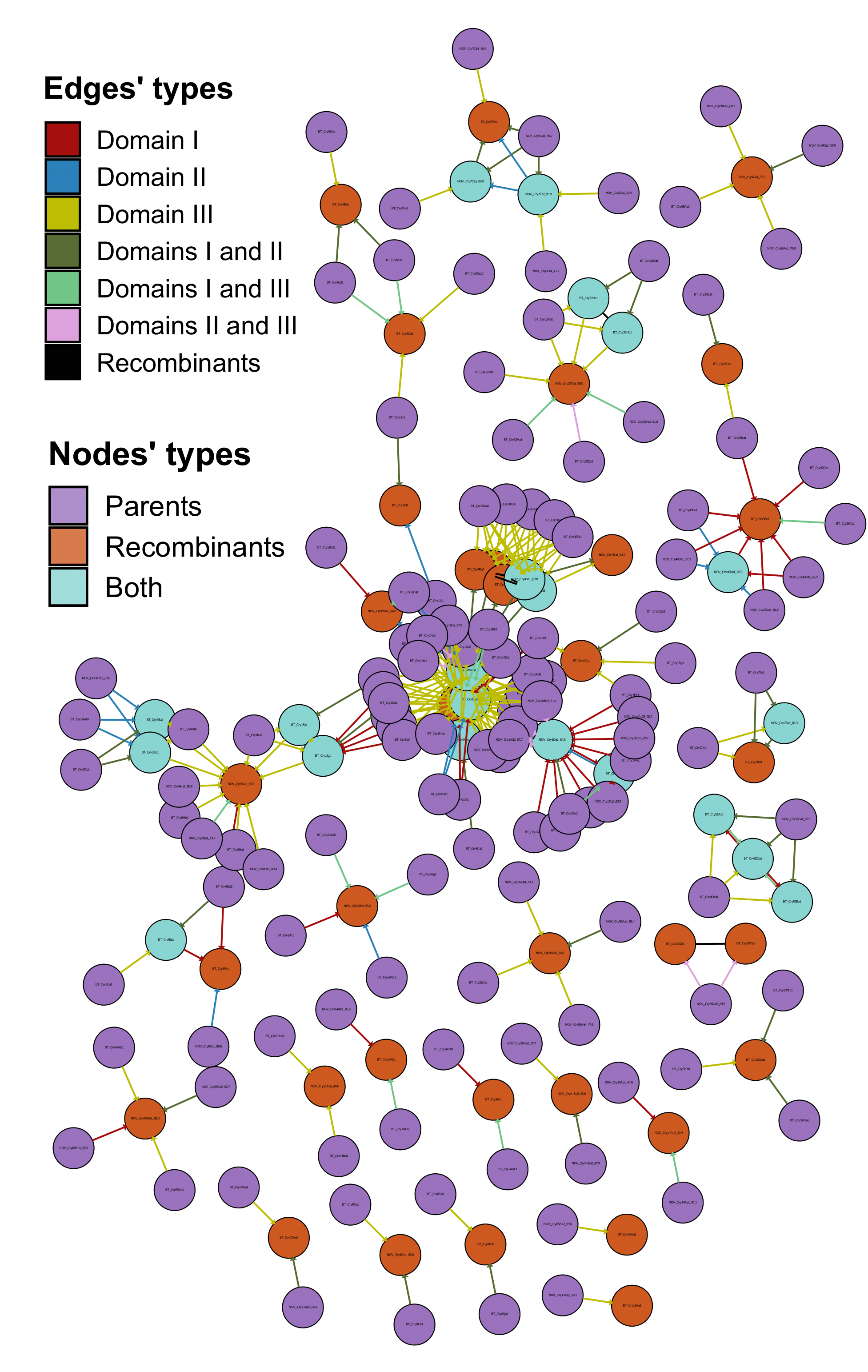

**Fig. S11**. The graph of domain exchanges between 3-D Cry toxins. The color of the node corresponds to the role of the toxin in recombination events, i.e., recombinants, parents, and those participating in multiple swaps thus being parents and recombinants simultaneously. Edges’ colors denote the domain transferred, namely, one domain for minor parents and two domains for major parents, respectively. Black edges connect toxins if a particular event contains multiple recombinants.

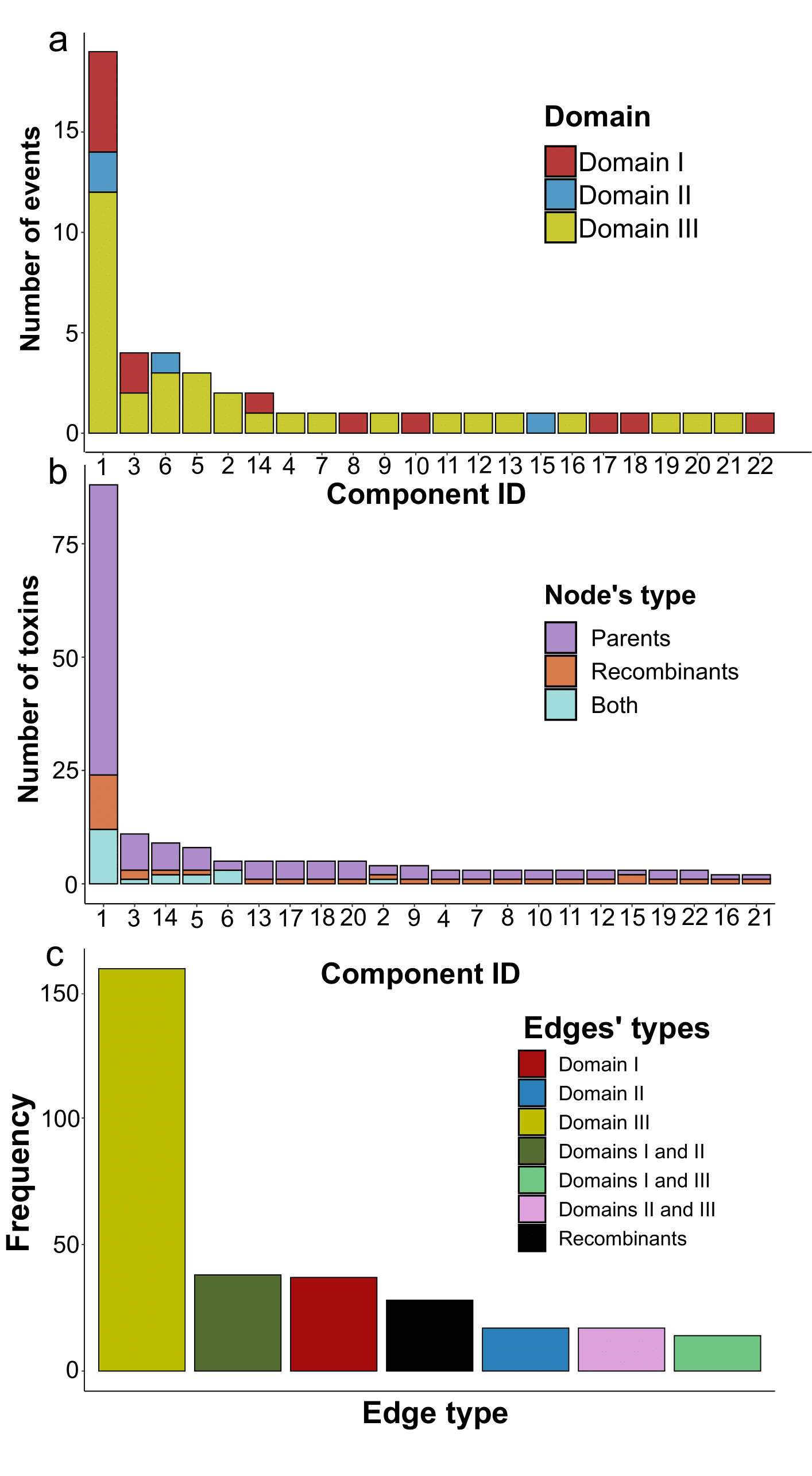

**Fig. S12**. Properties of weakly connected components of the recombination graph. (**a**) The number of recombination events within each connected component colorized according to the events’ types assigned with regard to transferred domains. (**b**) The number of toxins (nodes) within each connected component. The color corresponds to the role of the toxin in recombination events, i.e., recombinants, parents, and those participating in multiple swaps thus being parents and recombinants simultaneously. (**c**) The total number of graph edges within all connected components. Colors denote the domain transferred, namely, one domain for minor parents and two domains for major parents, respectively. Black edges connect toxins if a particular event contains multiple recombinants. In accordance with the distribution of events’ types, most of the edges refer to the third (161) and the first (41) domains as well as those presenting links from major parents amidst events involving the third domain with the first and the second domain transferred together (42).

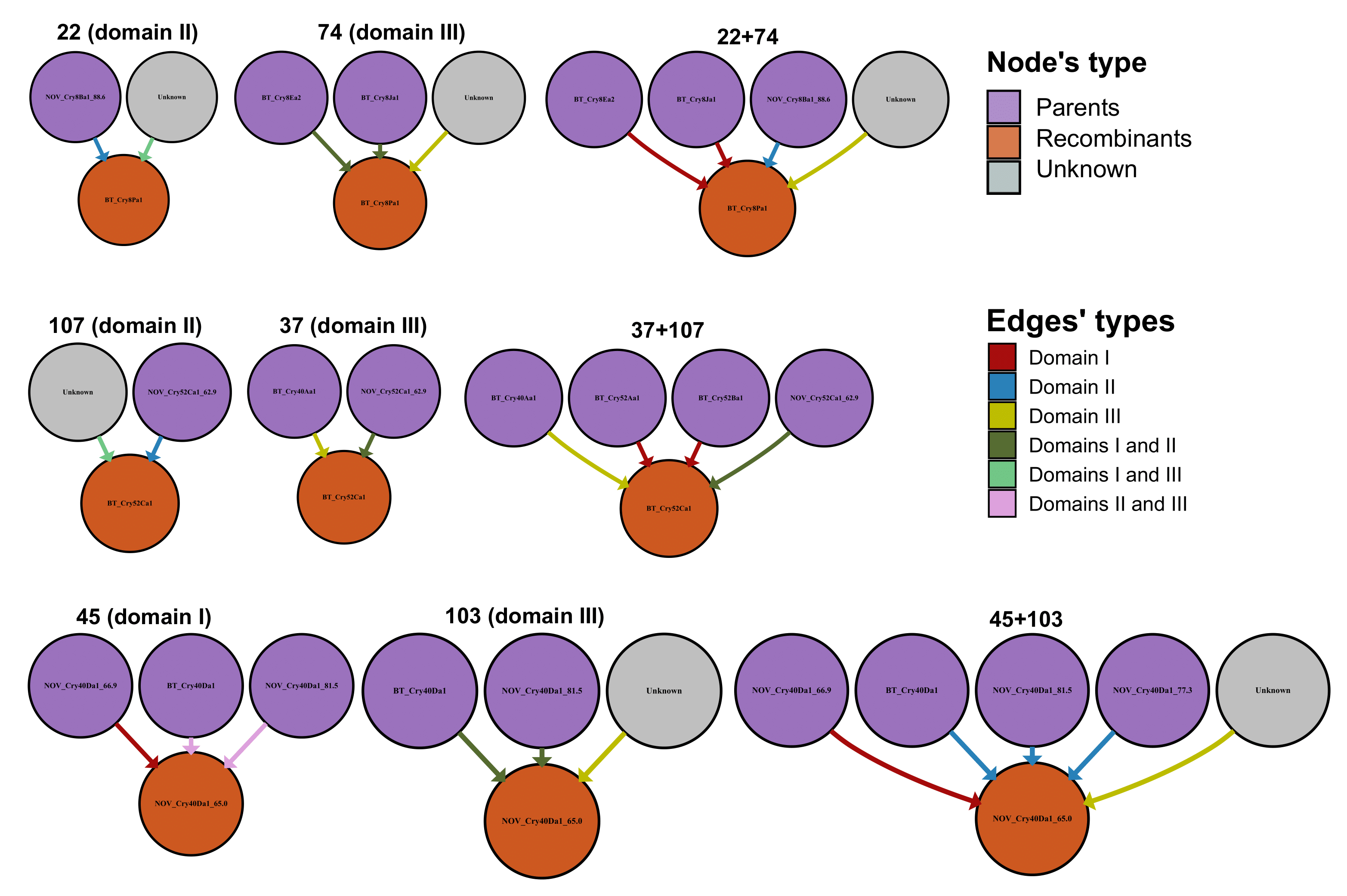

**Fig. S13**. Schemes of domain exchanges of linked recombination events, namely, those affecting the same recombinant. Shown are the initial parents reported by RDP4 and the corrected sets of parents for such events taken from the recombination graph. The color corresponds to the role of the toxin in recombination events, i.e., recombinants, parents, and unknown parents are shown as grey circles (unknowns are omitted in Fig. S11). The colors of the edges denote the domain transferred, namely, one domain for minor parents and two domains for major parents, respectively. In total, there were 3 pairs of events affecting 3 toxins: Cry8Pa1, Cry52Ca1, and Cry40Da1_65.0_CAH33957.1 with the latter demonstrating 65% similarity with the closest reference from the BPPRC (Bacterial Pesticidal Protein Resource Center) database (see tab. S23).

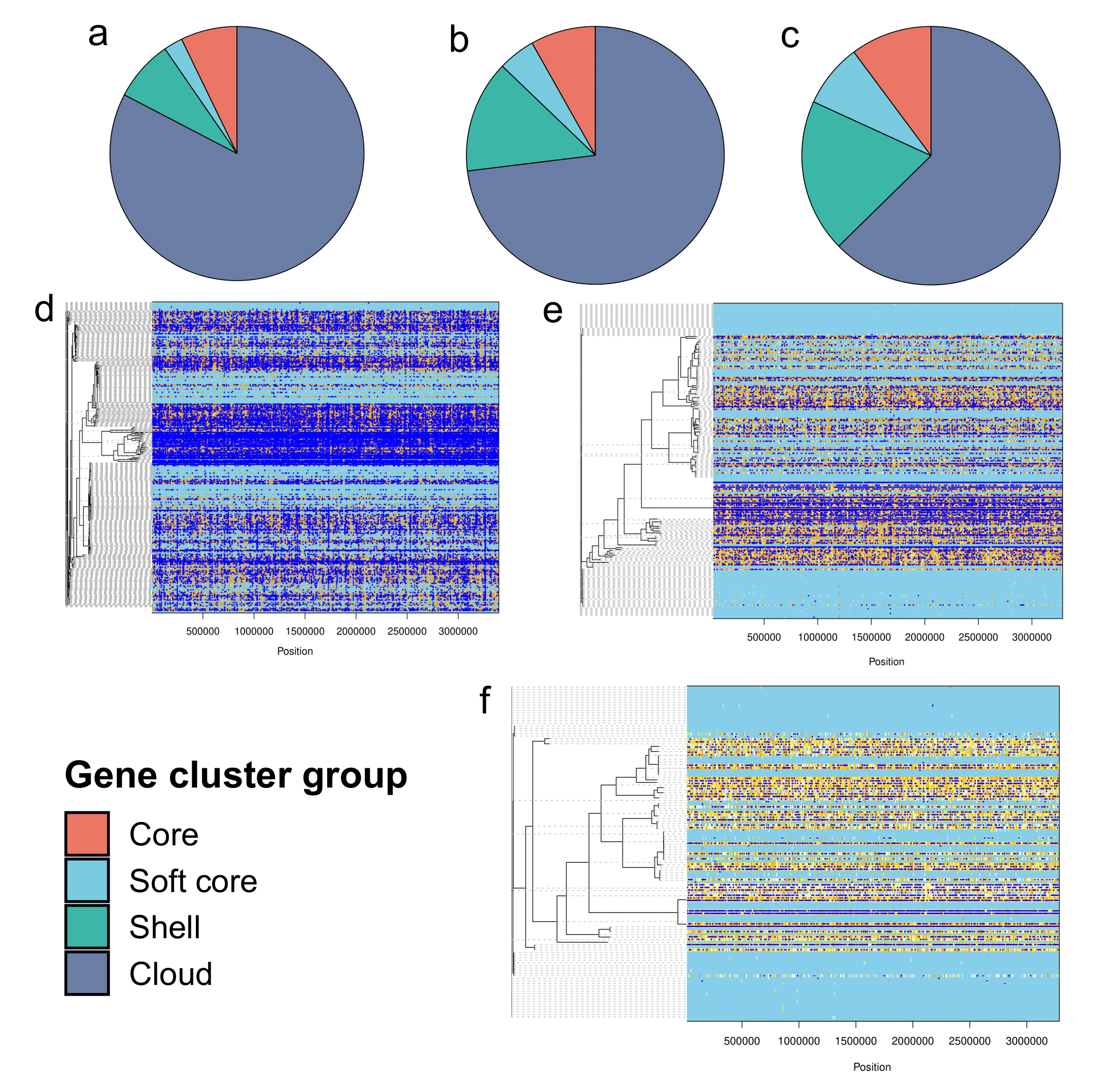

**Fig. S14**. Genetic diversity and recombination intensity if *Bt* assemblies bearing Cry toxins or lacking them. (**a**) The number of gene groups (core, soft core, shell, and cloud) within pangenomes reconstructed using Panaroo in all publically available *Bt* assemblies, as well as those containing 3-d Cry toxins (**b**), and participants of recombination events (**c**). (**d**) Regions susceptible to homologous recombination among all *Bt* genomes analyzed, assemblies bearing 3-d Cry toxins (**e**), and those harboring recombinants and parents involved in the domain exchanges (**f**). Recombination events detected with ClonalFrameML are marked with blue color and placed among concatenated core gene alignments obtained by Panaroo. Adjacent phylogenetic trees represent ClonalFrameML-corrected guiding trees reconstructed with FastTree from the core gene alignments.

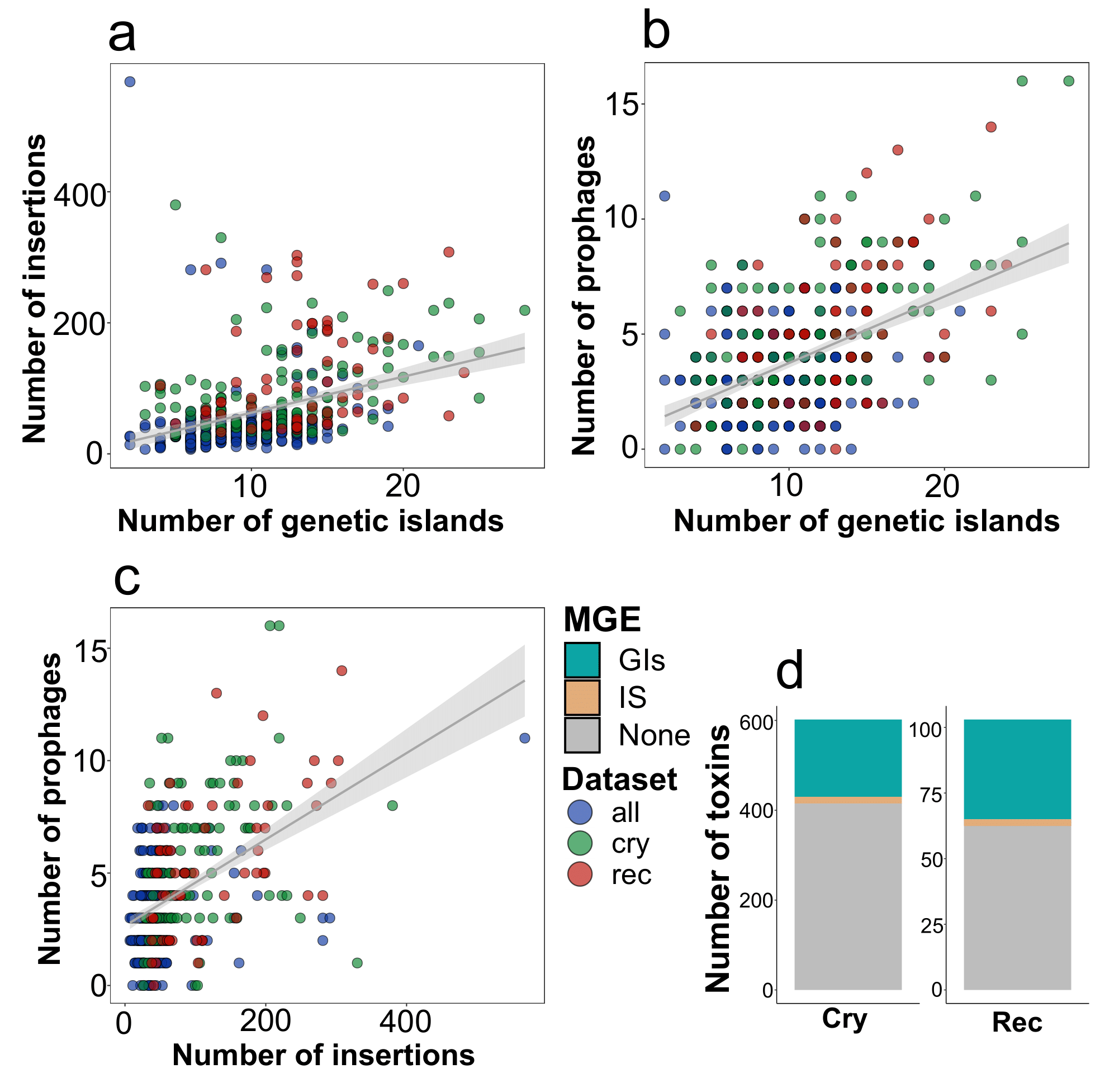

**Fig. S15**. The relation between mobile genetic elements (MGEs) and recombination events. The colors of points and box plots correspond to the sets of *Bt* assemblies, namely, all genomes (“*all*”), assemblies bearing Cry toxins (“*cry*”), and those incorporating participants in domain exchanges (“*rec*”). (**a**) Dependence between the total number of insertions (IS) and genetic islands (GIs), prophages and genetic islands (**b**), insertions and prophages (**c**) within the aforementioned sets of genomes. (**d**) The percentage of Cry toxins associated within the regions of IS and GIs. Shown are all 3-D Cry toxins and recombinants/parents involved in the domain exchanges.

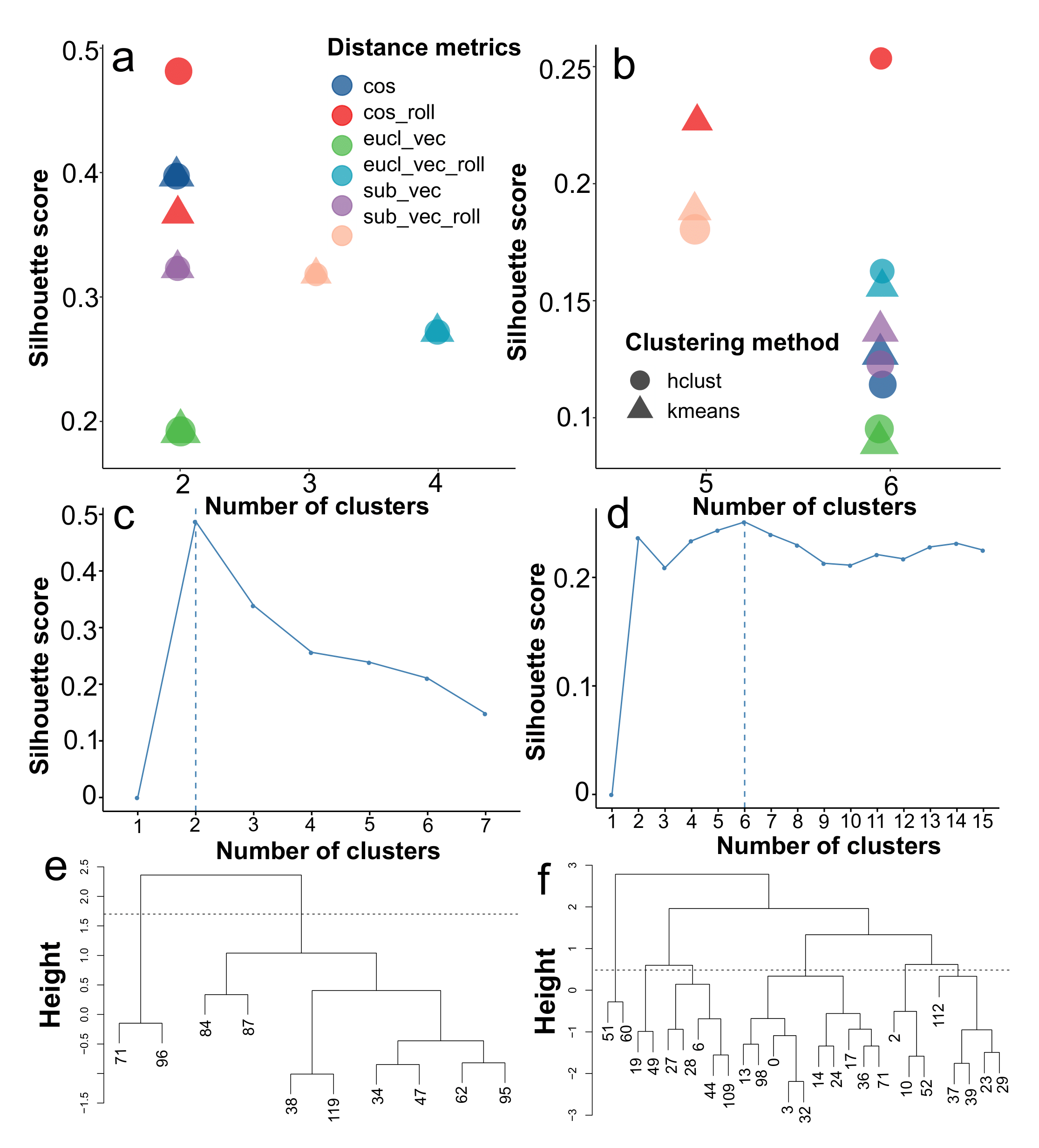

**Fig. S16**. Determination of optimal clusterization approach for event-wise groping of regions flanking breakpoints. The distance matrix used was based on integral estimates of per-coordinate identities between the sequences of major and minor parents within 100 b.p. regions surrounding the breakpoints. Three distance metrics were applied on the respective coordinate-wise estimated, namely Euclidean distance, Manhattan distance (the sum of the absolute distances), and cosine distance. The estimates are marked with prefixes ‘*eucl’*, ‘*sub’*, and ‘*cos’*, respectively. These methods were also used on the moving average of the identity values within the vectors to compare with a period of 7 b.p., which is delineated with the ‘*roll’* postfix. (**a**) Plotted is the most optimal number of clusters when using particular metrics according to the highest silhouette score calculated for the flanks located between the first and the second domain, and between the second and the third domains (**b**). The shape of the points encodes the clustering algorithm, namely, hierarchal clustering and k-means. Arrows indicate the best metric chosen for further analysis (**c**) The distribution of silhouette scores for a diverse number of clusters within the best method for the first and the second flanks (**d**). (**e**) Dendrogram visualizing hierarchal clustering patterns for the first and the second flanks (**f**). The numbers correspond to recombination events. The Grey dashed line indicates the threshold for clusters’ delineation.

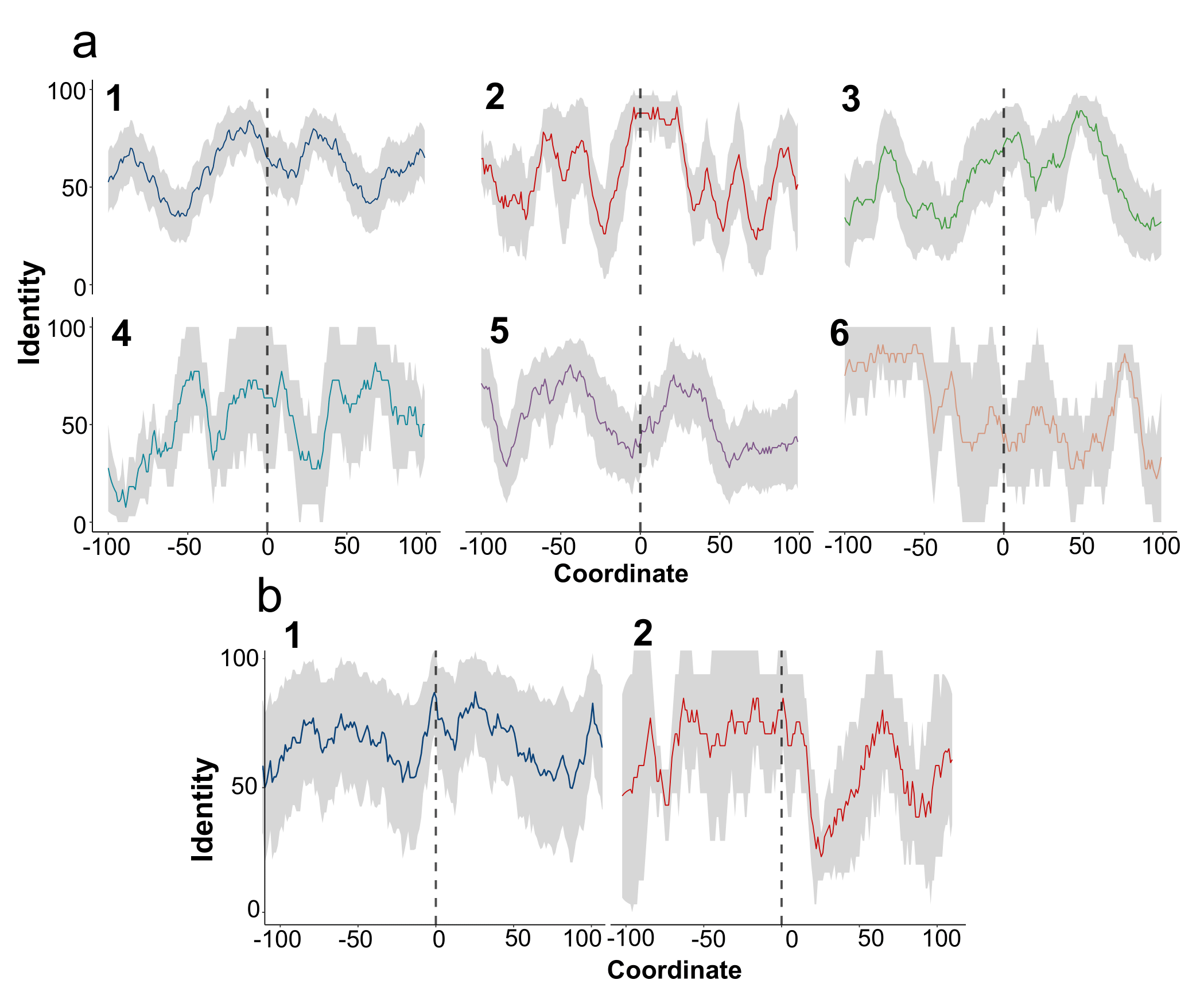

**Fig. S17**. Mean identity within regions surrounding recombination breakpoints. (**a**) Coordinate-wise mean identity between parental sequences in recombination events within two regions, namely between the second and the third domains and between the first and the second domains (**b**). Identity is calculated for 100 b.p. flanking regions before and after the respective breakpoints. Dashed lines indicate the location of breakpoints. Grey areas represent the standard error. The data is grouped according to the clusters of events obtained using the most optimal clustering pipeline and algorithm, namely hierarchical clustering based on the cosine distance metrics applied on the rolling average of the vectors with coordinate-wise identities.

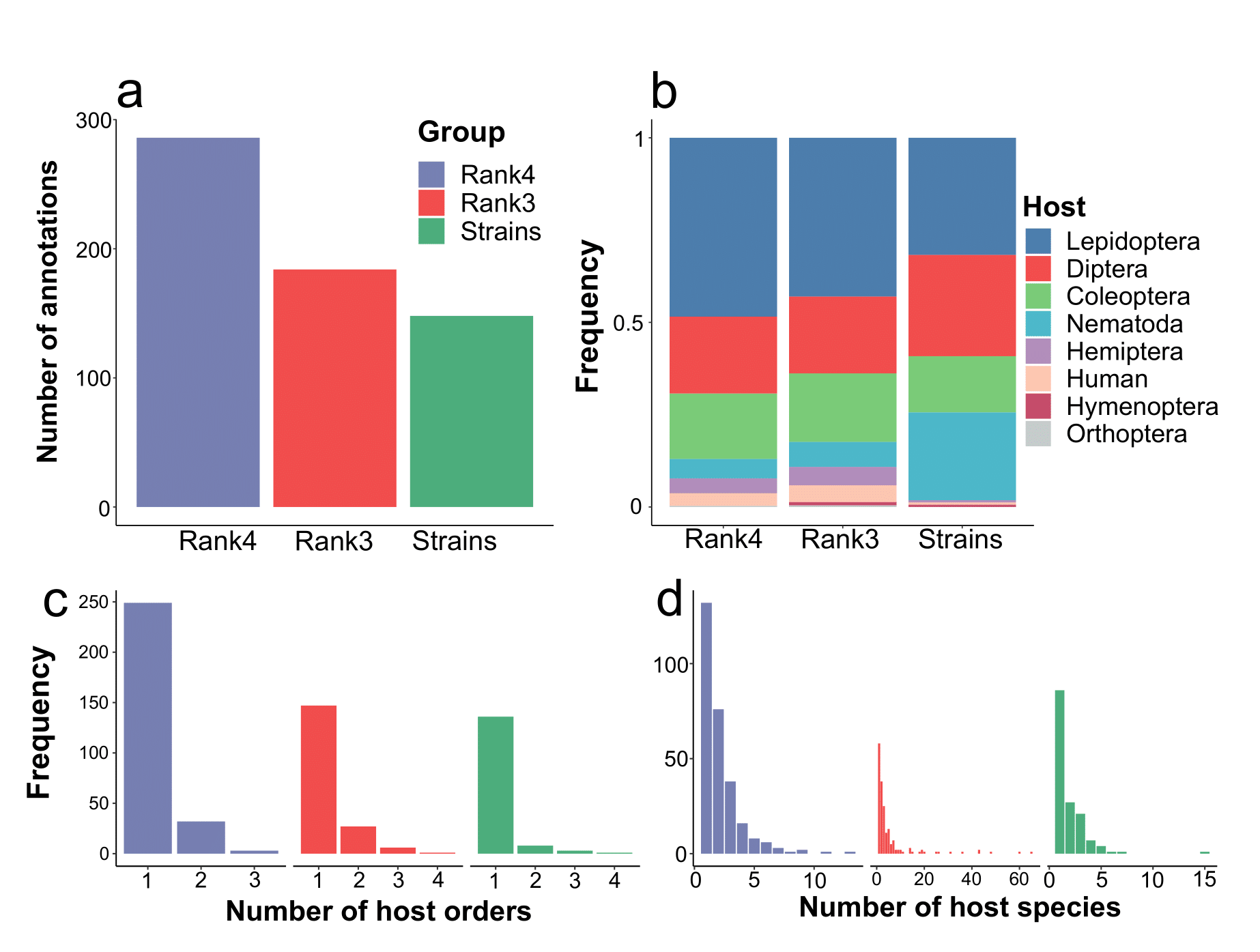

**Fig. S18**. Overall distribution of activities of studied Cry toxins and strains encompassing them. (**a**) The total number of instances having at least one known host to which the activity was shown. Three groups are considered, namely proteins within the fourth (100% identity) and the third rank (95% identity within the toxins’ group), and strains. (**b**) The proportion of individual bioassays made for toxins and strains. The colored within-bars fraction corresponds to the number of unique species attributed to higher taxonomical categories (**c**) The frequency of strains and proteins in the context of the amount of known affected host orders and species (**d**).

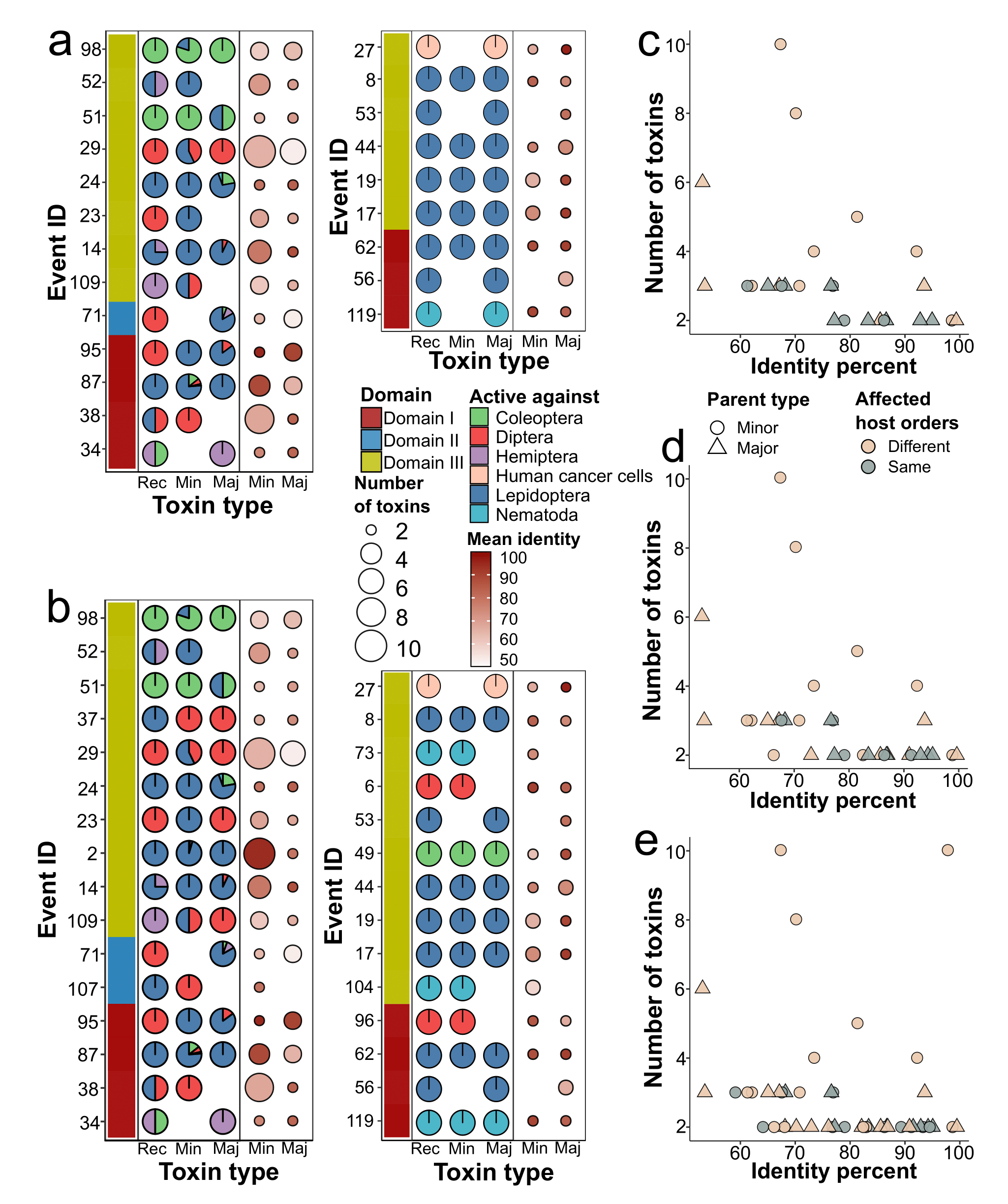

**Fig. S19**. The properties of parents and recombinants in terms of the hosts attributed to them. (**a**) The percentage of species attributed to different orders against which recombinants and parents are active. The data on the activity is based on the third rank of the toxins and strains if the respective protein-wise host spectrum is unknown (**b**). The portion of particular orders within the pie charts is color-coded. The events are split into two panels, namely those in which at least one toxin exhibits the activity to the order differing from other types of parent and/or child. The left adjacent bar corresponds to the domain transferred in the particular event. The right adjacent panel shows the number of toxins (recombinants and the parents, either major or minor ones). ‘*Rec*’ indicates recombinants, ‘*Min*’ – minor, and ‘*Maj*’ – major parents, respectively. The size of the dots is proportional to the number of participants, whereas the intensity of the color delineates the mean pair-wise identity between the transferred domains. (**c**) The dependence between the number of toxins and the mean identity of the transferred domain in the sets of events for which the associated hosts are available on the basis of third and fourth (**d**) ranks of the toxins and strains as well (**e**). The shape of the dots encodes the type of parent, and the color indicates whether the sets of host orders are the same or different.

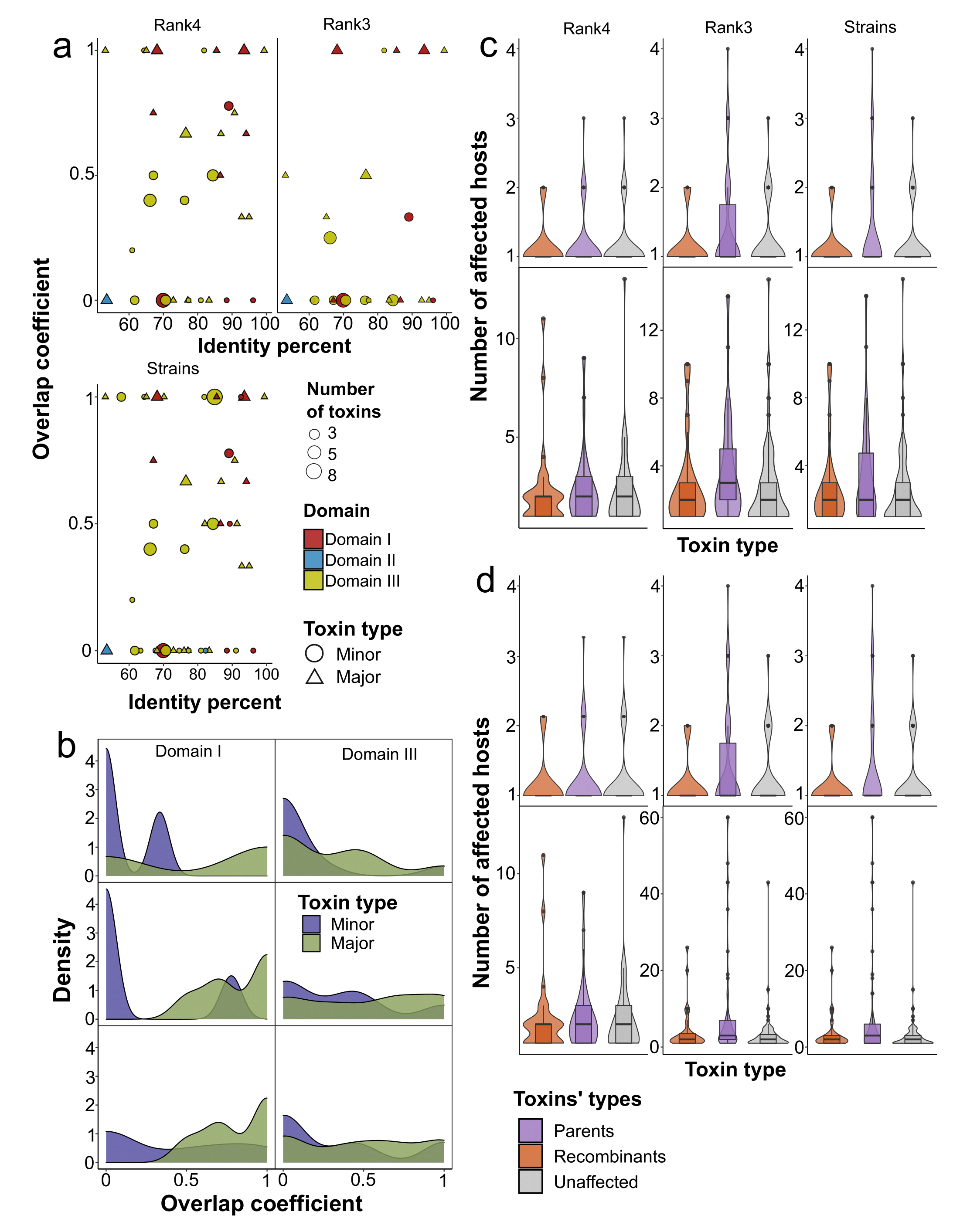

**Fig. S20**. The comparison between the composition and the number of affected hosts for parents and children within recombination events and toxins devoid of domain exchanges as well. (**a**) The relationship between the Szymkiewicz–Simpson coefficient (overlap coefficient), i.e., the intersection between the host species attributed to recombinants and parents divided by the size of the smaller set of hosts. Lots are based on the data either from the fourth or third ranks of toxins and strains. The size of the points is proportional to the number of toxins. The shape corresponds to parent types, and the color is determined by the transferred domain. (**b**) Density plots showing the frequency of overlap coefficient estimates within the events implying exchanges of the first and the third domains. The color of the plot corresponds to the type of parents, and the panels from top to bottom are determined by the sets of events dependent on the source of host metadata. The distribution for the second domain was unavailable to calculate, given the scarcity of the data. (**c**) The mean number of the affected host species and orders of different toxin types, namely recombinants, parents, and toxins without recombination signals determine the color of violins/box plots. Toxins that are associated with less than 15 species or all the proteins (**d**) were taken into account. Significant differences according to the Wilcox test with p-values adjusted using the Benjamini-Hochberg procedure are marked with asterisks.

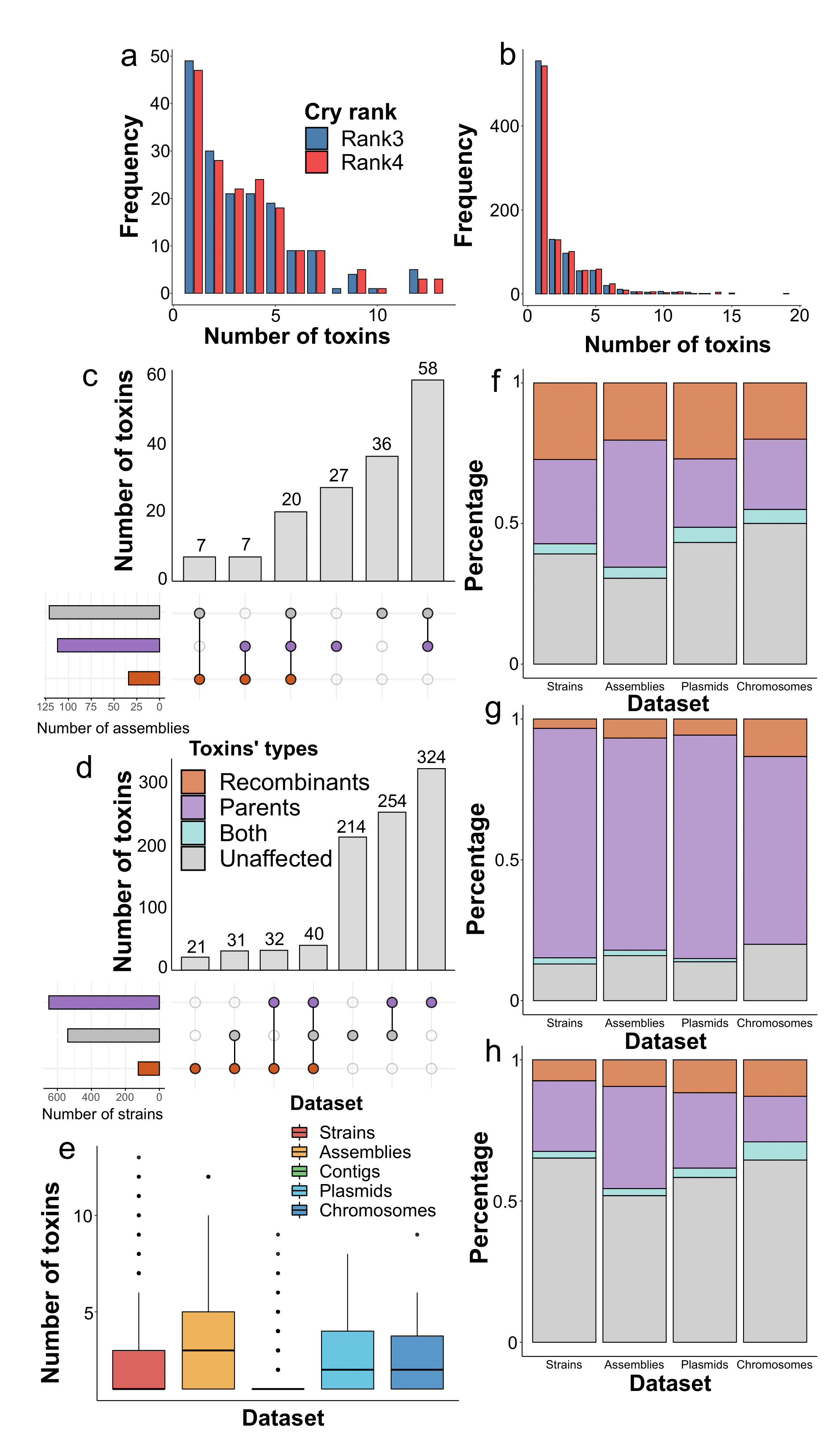

**Fig. S21**. The properties of assemblies and strains in terms of the types of toxins they contain in the context of recombination events. (**a**) The distributions of the number of toxins in the assemblies and strains (**b**) regarding the rank of the respective Cry proteins. (**c**) Upset plots displaying the number of assemblies and strains (**d**) in the context of toxin types they include. The respective instances are classified according to the composition of toxins, namely, parents, recombinants, and the unaffected Cry proteins. Intersections indicate that the set of assemblies/strains contains different types of toxins. (**e**) The mean number of Cry toxins within different datasets, namely, strains, assemblies, and genomic regions within the assemblies, single contigs, plasmids, and chromosomes. (**f**) The fraction of particular toxin types in the datasets containing recombinants, parents (**g**), and unaffected toxins (**h**).

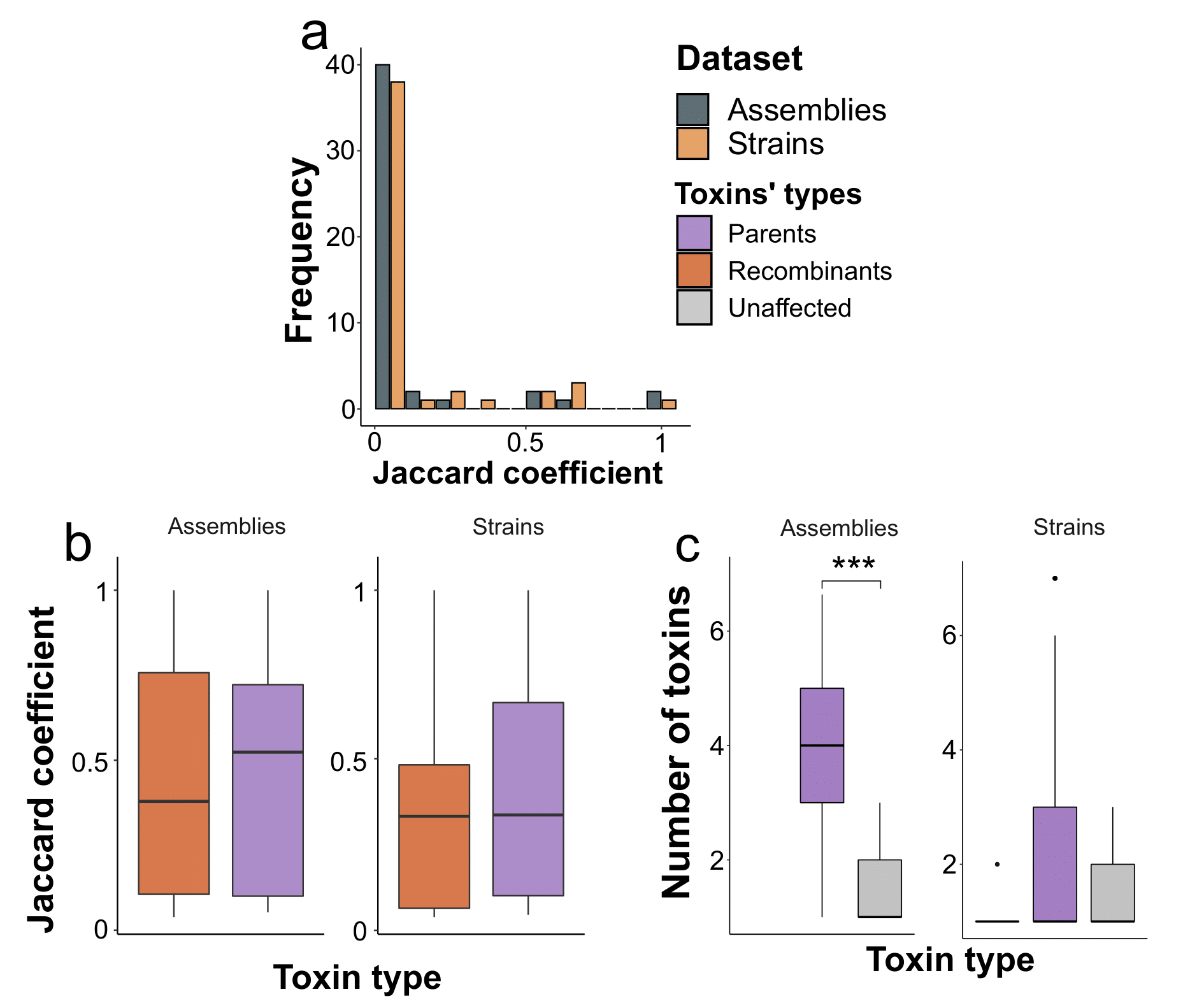

**Fig. S22**. Comparison between the amount and compositions of cry toxins belonging to strains and assemblies including participants in domain exchanges and the unaffected proteins. (**a**) The measure of similarity of all sets of toxins that strains and assemblies in which at least one parent is present. Event-wise comparisons using the Jaccard coefficient (the length of the intersection divided by the union of the sets) refer to toxins’ compositions in assemblies and strains attributed to minor and major parents of the recombination events. (**b**) Comparison between the above-described Jaccard coefficients exceeding zero when comparing recombinants with parents and only parents, respectively. (**c**) The number of toxins within assemblies and strains that contain toxins of the particular type exclusively such as recombinant, parents, and unaffected Cry proteins. Asterisks denote significant differences according to the Wilcox test with p-values adjusted using the Benjamini-Hochberg procedure.

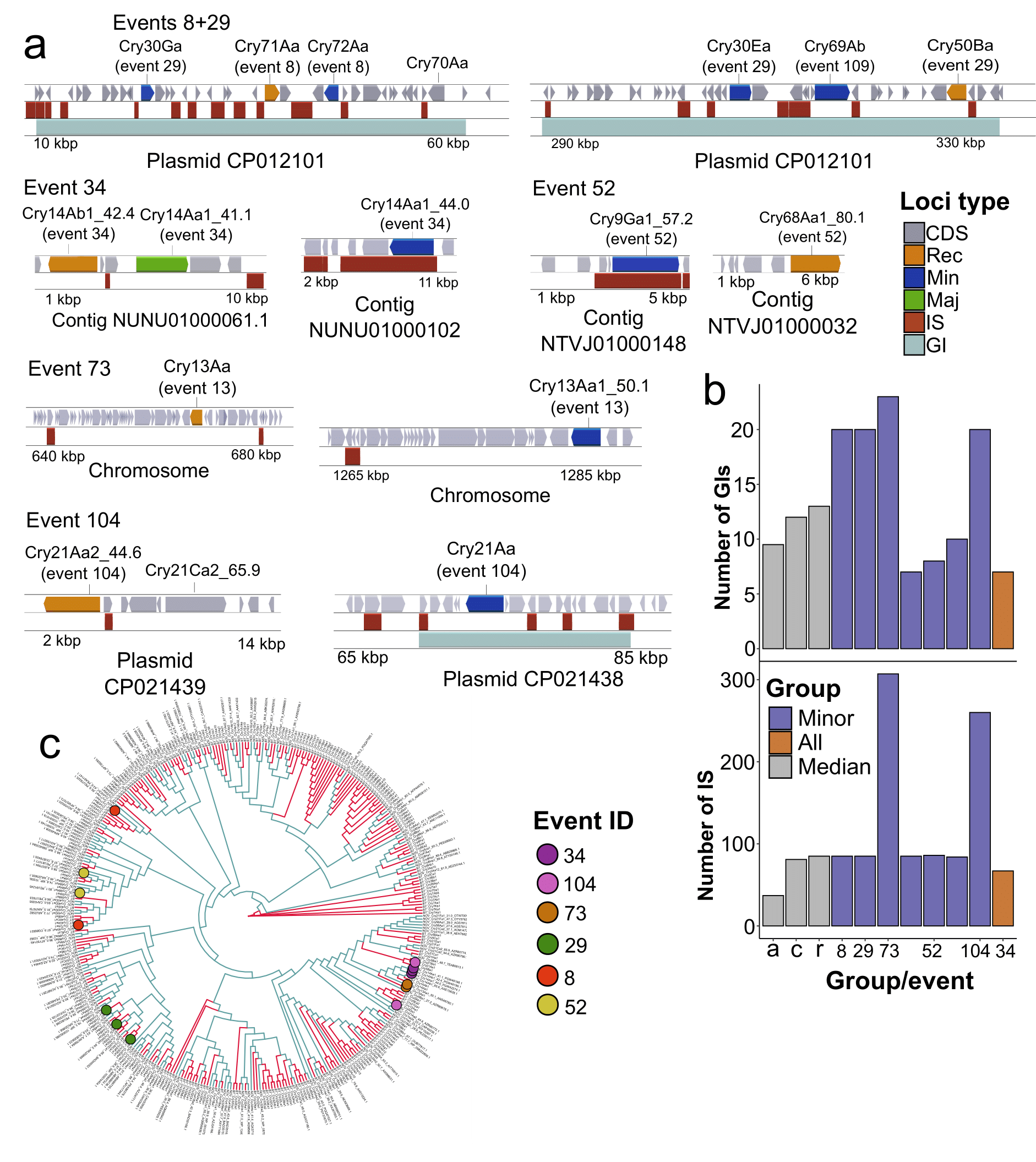

**Fig. S23**. Genomic context of genes encoding parents and recombinants from one event found in the same assemblies. (**a**) Loci surrounding recombinants and parents from individual recombination events. Shown are regions, plasmids, chromosomes, and contigs bearing the genes of interest. Three tracks are presented, namely, genes, insertion sequences (IS), and genetic islands (GIs). The color of the blocks on the first track corresponds to the toxins’ role in recombination events, e.g., major and minor parents, recombinants, and unaffected toxins. (**b**) Comparison of the total number of IS and GIs presented in genome assemblies that contain genes coding for recombinants and parents from the same events. Grey bars display median estimates for all assemblies studied, genomes with cry genes, and with toxins subjected to recombination, which are marked with letters ‘*a*’, ‘*c*’, and ‘*r*’ on the X-axis, respectively. Numbers represent IDs of recombination events. Event 52 was associated with genome assemblies, however, the genomic context was highly similar, thus, only one example is given. (**c**) Relative positions of Cry proteins on the full-lengthed partitioned phylogeny that co-occur in the same assembly. The color of circles defines individual recombination events.

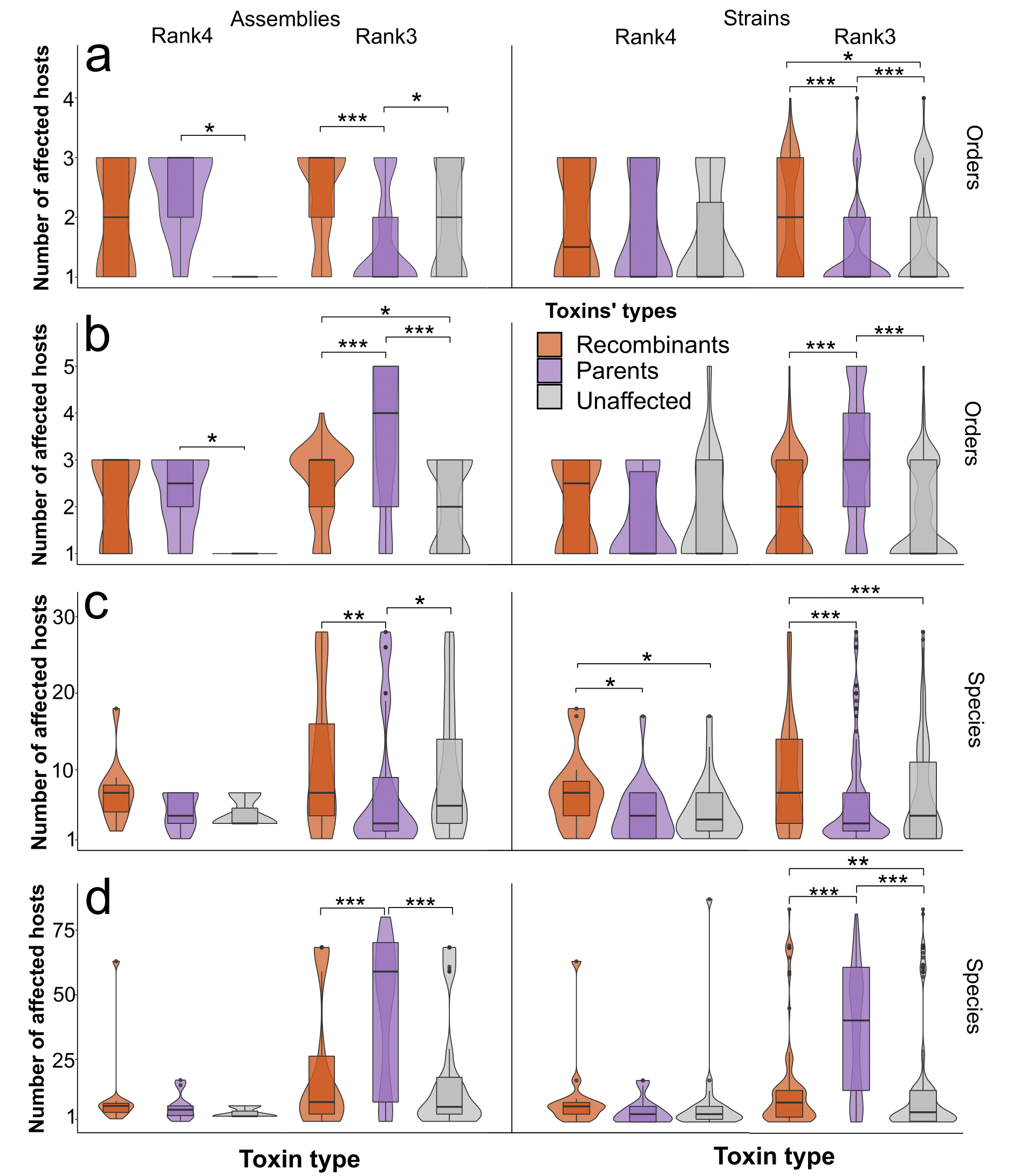

**Fig. S24**. The number of affected host species and orders by strains and assemblies with or without toxins subjected to domain exchanges. (**a**) Distributions for the number of affected orders within the sets of strains and assemblies omitting the toxins associated with more than 15 species or considering them (**b**). (**c**) The respective data for species without and with (**d**) well-studied toxins. The metadata taken is grouped according to the toxin ranks used. Significant differences according to the Wilcox test with p-values adjusted using the Benjamini-Hochberg procedure are marked with asterisks.

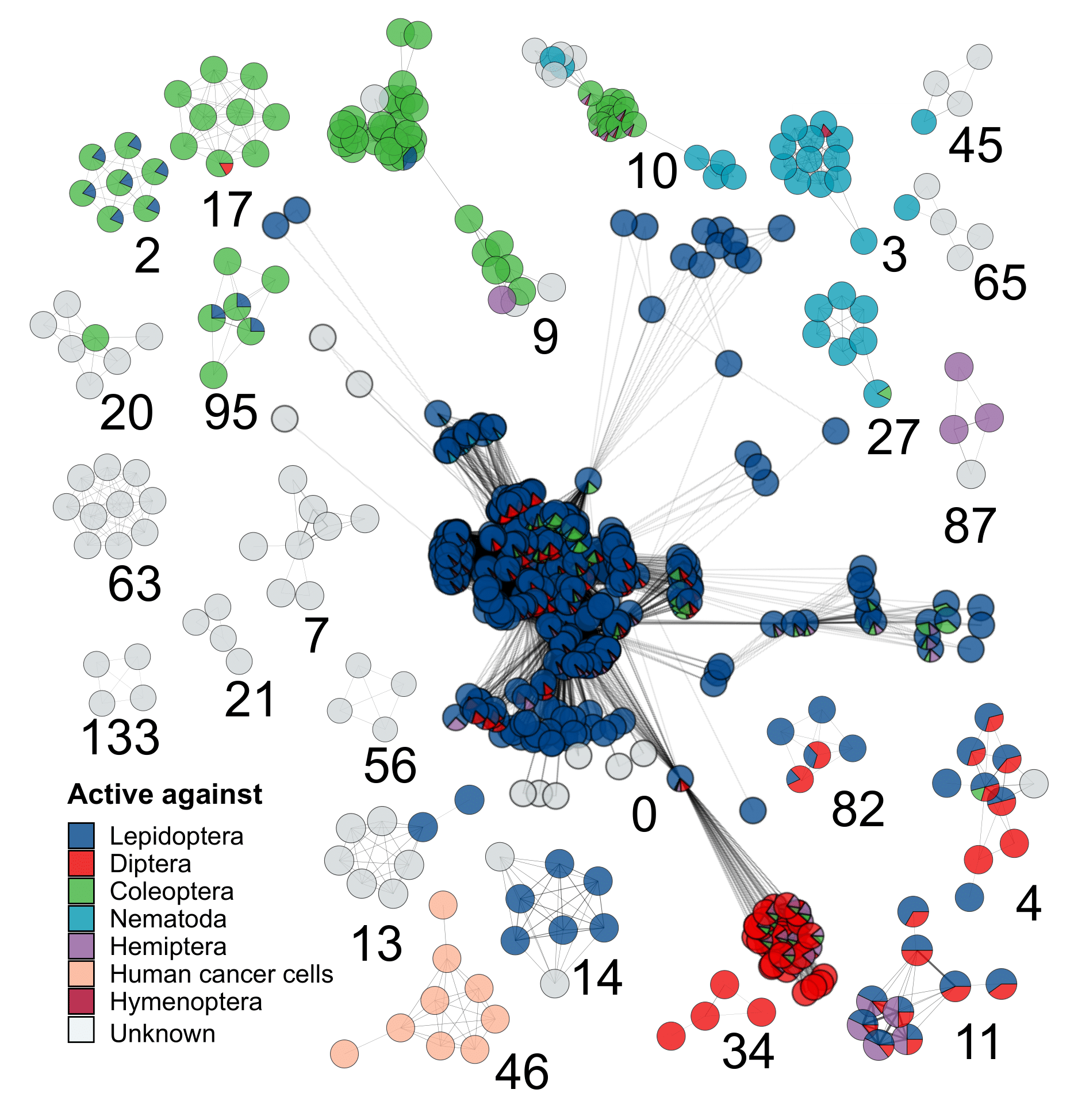

**Fig. S25** Weighted specificity-wise graph of strains sharing the common cry toxins. The nodes represent strains, whereas edges indicate that at least one toxin within the third rank is shared by the strain. The weight of the edges depends on the number of common toxins. The color of the node represents the overall proportion of species that are affected by the toxins found in the strain. Shown are connected components containing at least four nodes included. The IDs of the respective components are presented in numerical form.

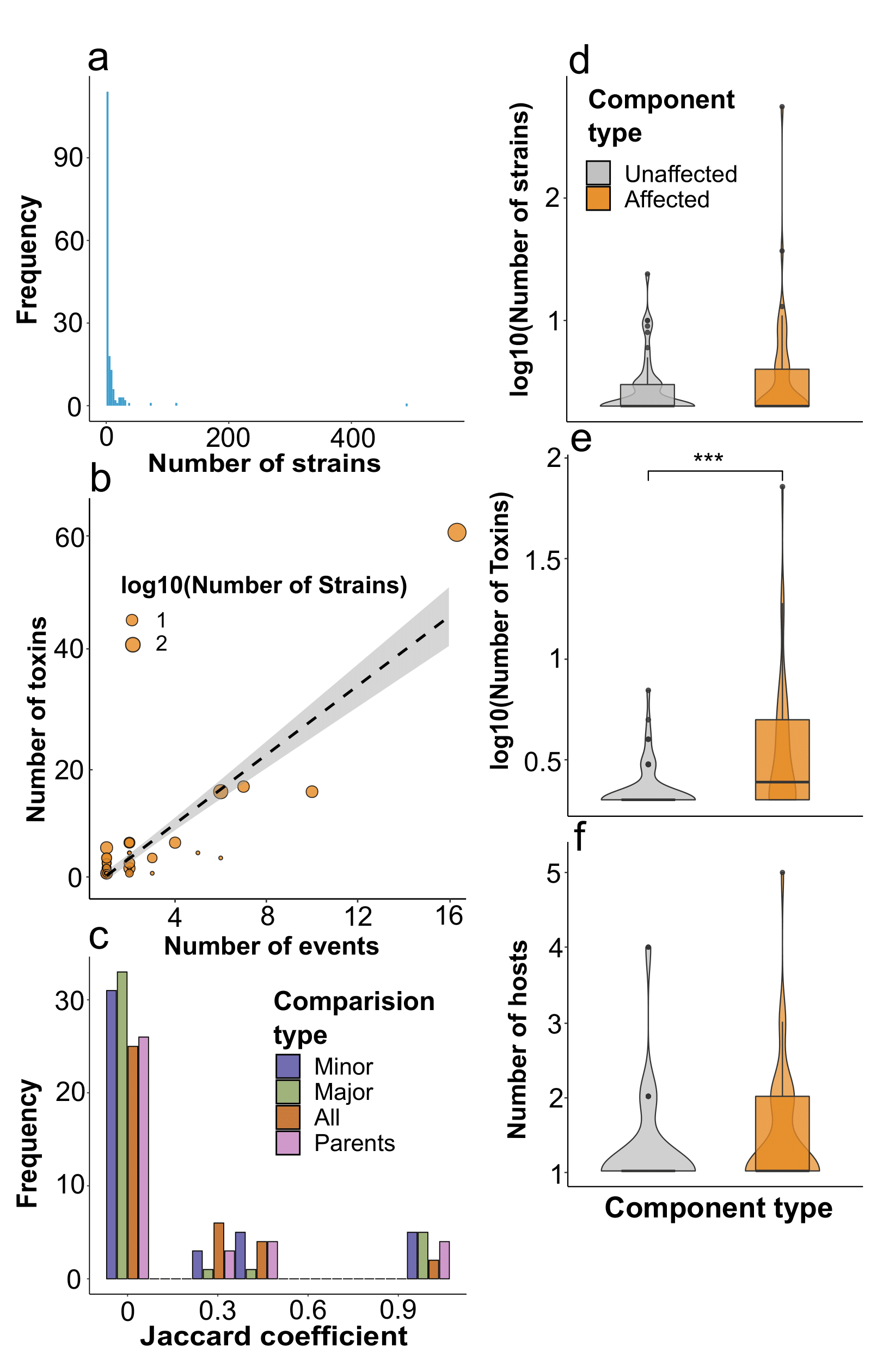

**Fig. S26**. Overall properties of connected components of the strains’ graph and comparison between components. (**a**) The frequency of connected components according to the number of nodes (strains) forming them, the arrow points to the largest component. (**b**) relationship between the number of toxins within the components and the number of recombination events, which are associated with them, the grey dashed line shows the dependencies estimated by the linear model. The size of the dots is proportional to the decimal logarithm of the number of nodes. (**c**) Distribution of the overlaps between the connected components in which parents and recombinants from the particular event reside. The color indicates the type of comparisons, namely, recombinants with parents, either minor, major, and all, and only between parents, respectively. The similarity is calculated using the Jaccard coefficient. (**d**) Comparisons between the components containing parents and children with the ones devoid of them in terms of the number of nodes (strains), individual toxin types (**e**), and orders affected (**f**). Significant differences according to the Wilcox test are marked with asterisks.

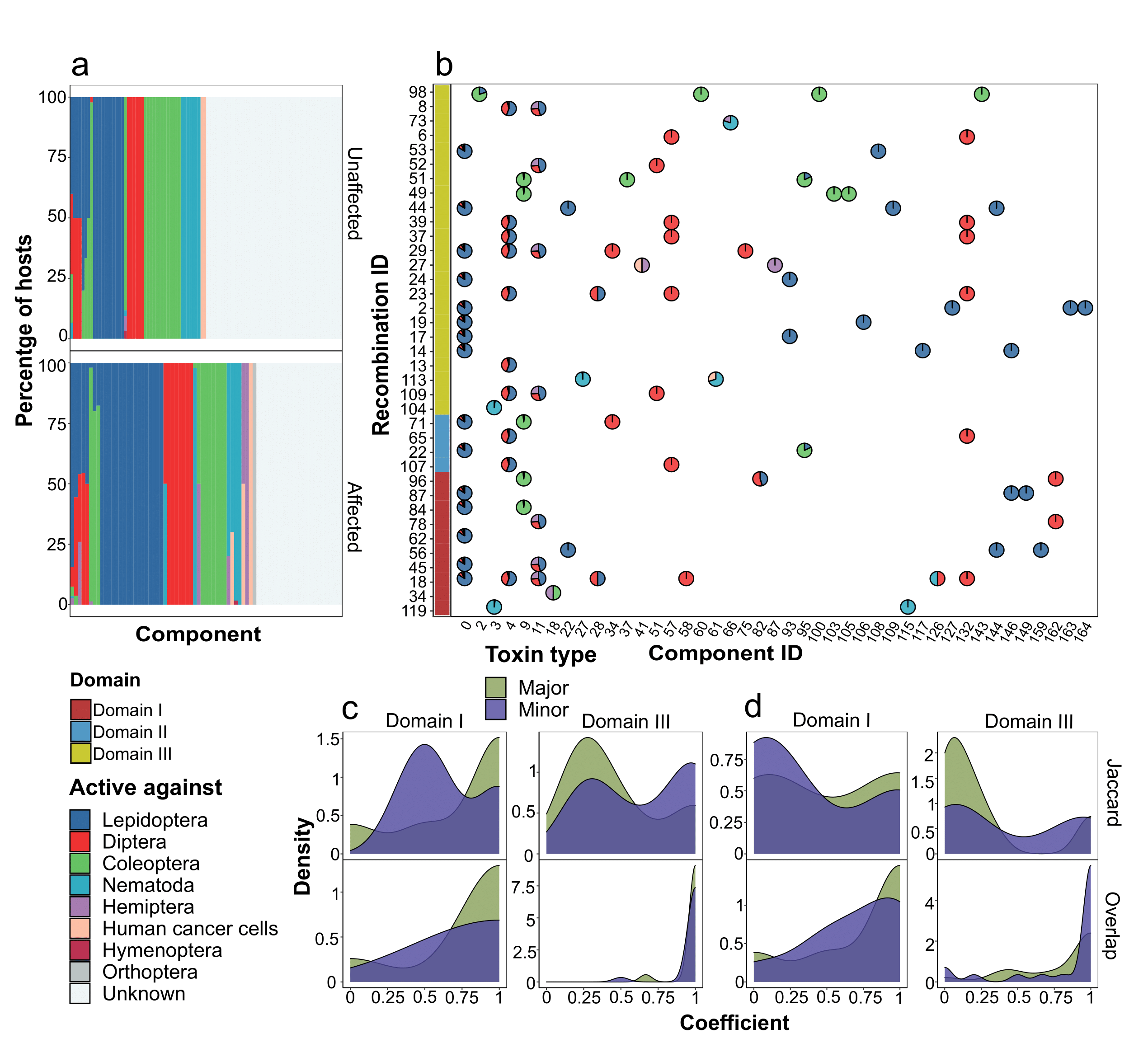

**Fig. S27**. Features of the connected components including recombinants, parents, and unaffected toxins in terms of the hosts they exert an effect on. (**a**) The percentage of host species attributed to their orders for strains within the connected components comprising toxins that have undergone recombination or not containing them. Plotted on the x-axis are individual components (**b**) The distribution of affected orders for the components associated with toxins from recombination events. The size of sectors of pie charts is proportional to the fraction of host species belonging to particular orders. The left adjacent panel indicates transferred domains within the events. (**c**) Density plots showing distributions of similarity estimates regarding the sets of orders and species (**d**) affected by recombinants and parents, either minor or major. The plots are presented only for the first and the third domains due to scarcity of the data regarding the second domain. Two similarity measures are considered, namely the Jaccard and overlap coefficients, respectively.

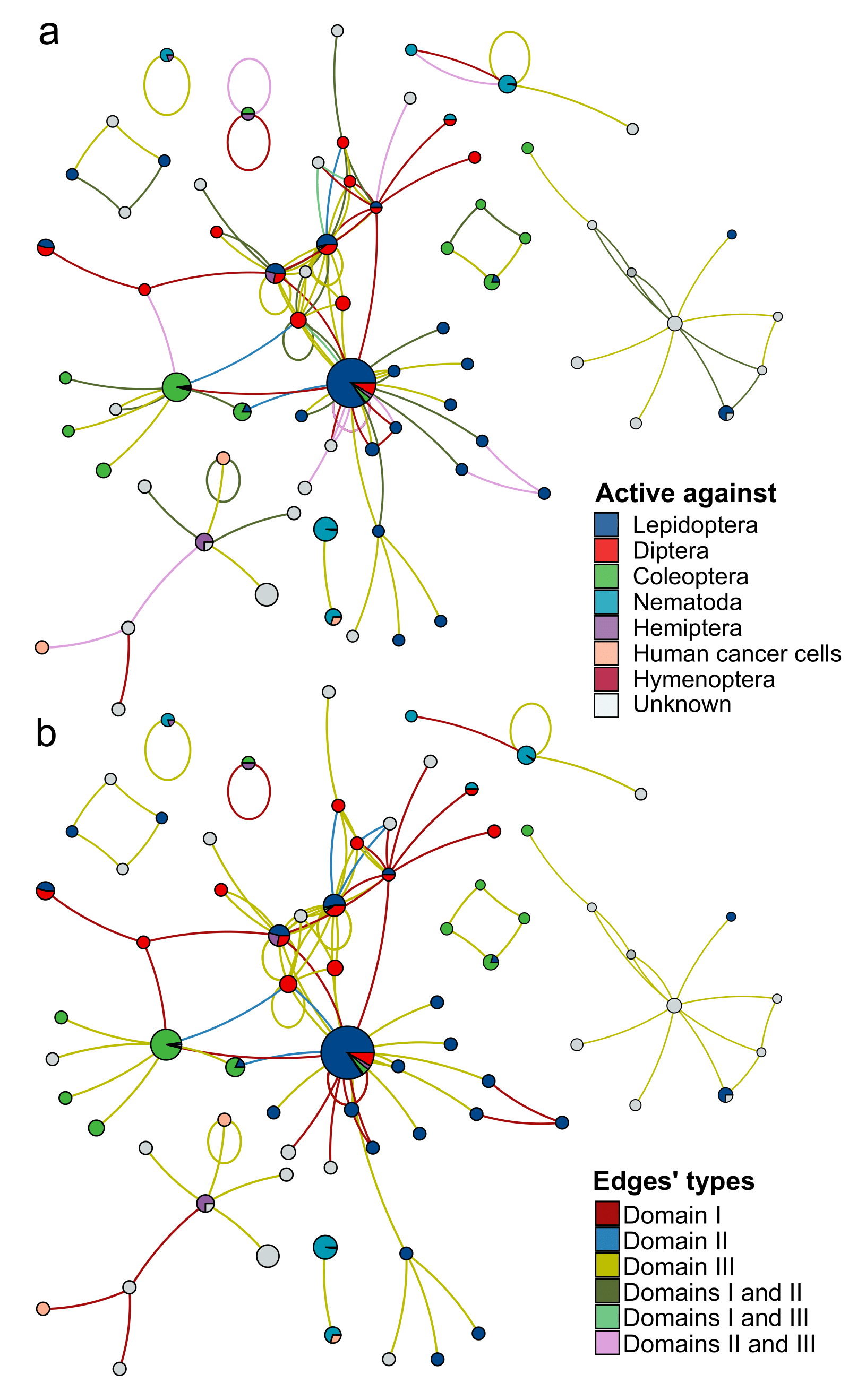

**Fig. S28**. Combined graph showing relationships between subpopulations of bacterial strains based on recombination events between toxins they produce. In the graph, nodes represent the connected components in the weighted specificity-wise graph of strains sharing the common Cry toxins (fig. S25). The color of the node represents the overall proportion of species that are affected by the toxins found in all strains within the connected components. Edges show domain exchanges between the toxins within the connected components colored according to the transferred domains. The edges represent all parents grouped according to domains (**a**) or the recombination events classified according to domains obtained from minor parents (**b**).

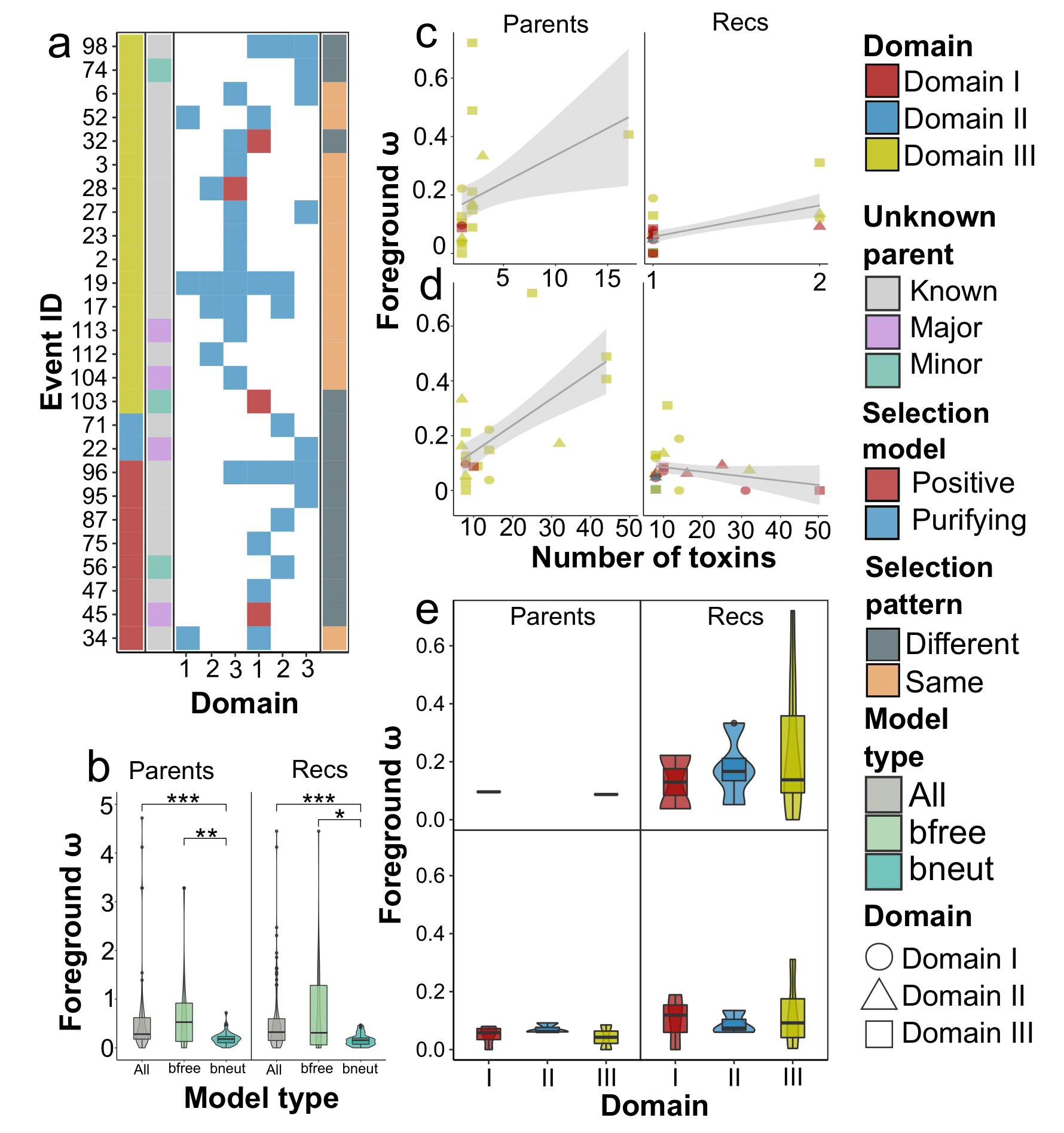

**Fig. S29**. Results of evolutionary selection assessment calculated on branches. (**a**) Optimal evolutionary models calculated on branches in which recombinant and parental sequences were marked. Here and in other figures the selection pressure was determined on the sequences of different domains independently. The heatmap shows the models reported for the particular domain. The left adjacent subpanels correspond to the domain that was transferred from the minor parent within domain exchanges and show whether a parent, either minor or major, is unknown, i.e., absent in the dataset analyzed. The color of the blocks represents the model chosen, namely, positive and purifying selection. The panel is divided into two blocks with the first one associated with parents and the second with recombinants, respectively. The right subpanel indicates whether for at least one domain a different selection pattern from the respective parental sequence exists. (**b**) Mean ω values (the nonsynonymous to synonymous substitution rate ratio, dN/dS) based on branch models. “All’ denotes non-filtered inferences, “bfree” indicates events for which the LRT (log-likelihood test) revealed significant differences from the neutral model, “bneut” marks the set of events for which selection within the foreground branches differs from the overall background (**c**) Relationships between ω and the number of marked toxins (recombinants and parents) or the total number of toxins (**d**) within the subtrees tested. The shape of the points indicated the domains for which models were assessed, while the color corresponds to the domain transferred during recombination. The ω value is positively and significantly correlated (p < 0.05 according to the linear model) with the number of marked toxins in the tree (both parents and recombinants), which, however, was backed only by several examples with a larger number of proteins (**c**). If all toxins are considered (**d**), significance is retained for parents only. (**e**) Average omega ω values based on branch models for the events affecting the first and the third domains of parents and children, respectively.

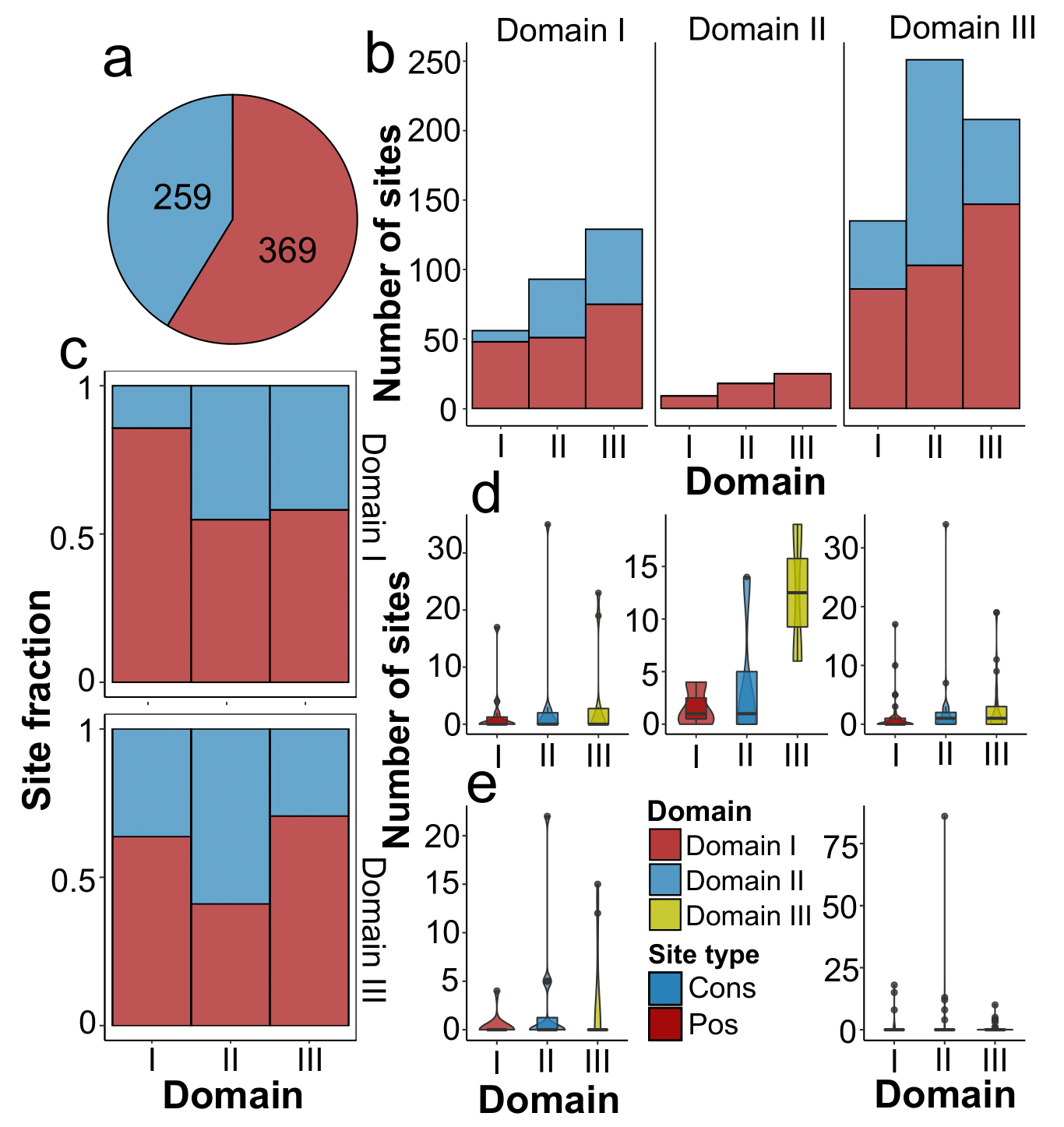

**Fig. S30**. Distribution of sites subjected to evolutionary selection according to site models. Here and in other figures the selection pressure was determined on the sequences of different domains independently. (**a**) The total proportion of conservative and positively selected sites revealed by best-fit site models determined by the LRT (likelihood ratio test). (**b**) The number of positively selected and conservative sites among domains within the sets of events grouped according to the transferred domain and the fraction of these sites (**c**) in exchanges of the first and the second domains. (**d**) The overall number of positively selected and conservative sites (**e**) for events affecting the first and the third domains. The color corresponds to the domain for which the selection model was determined.

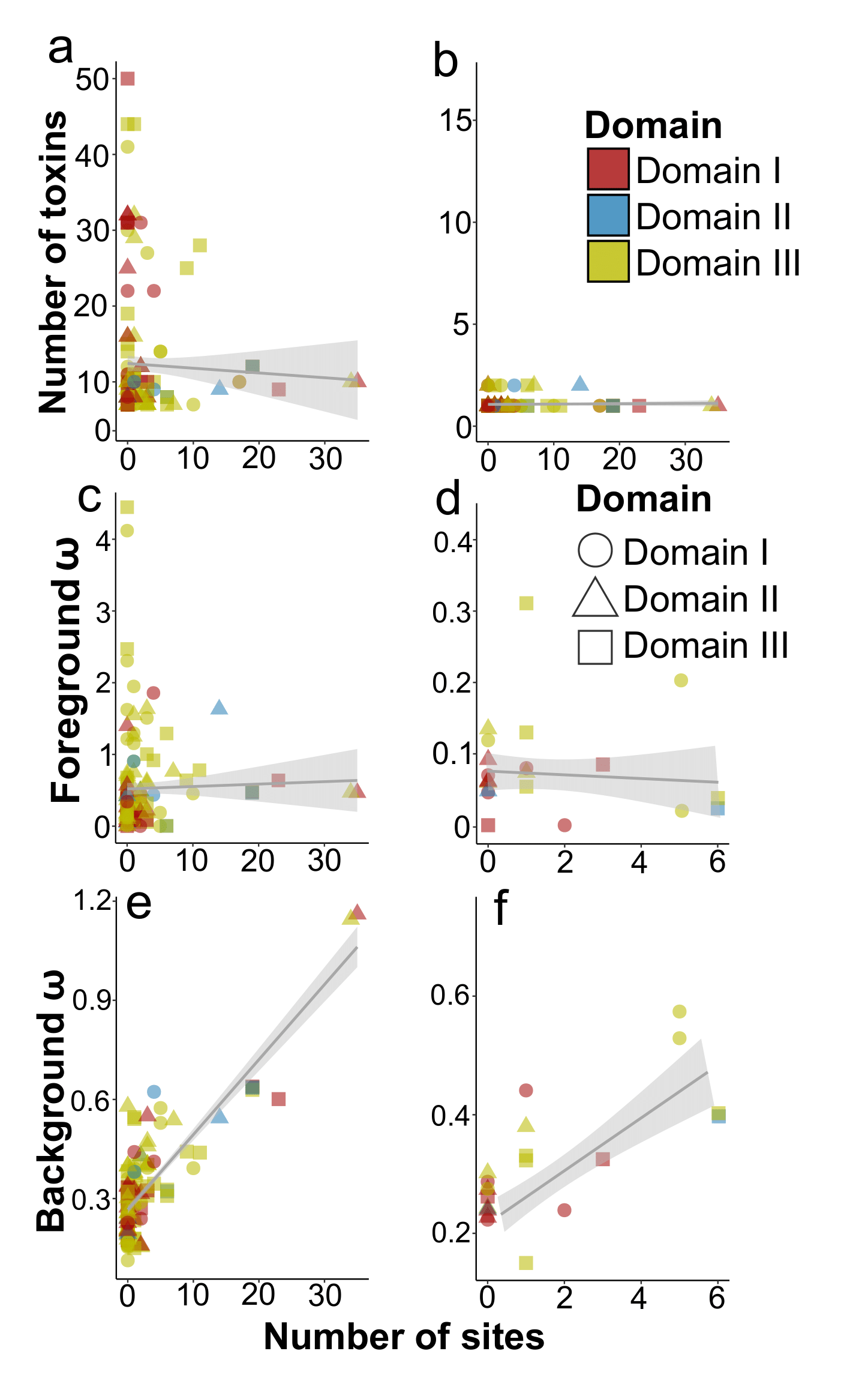

**Fig. S31**. Revealing factors possibly delineating the number of positively selected sites according to evolutionary selection models calculated on sites. Best-fit site models are determined by the LRT (likelihood ratio test). Here and in other figures the selection pressure was determined on the sequences of different domains independently. The shape of the points indicated the domains for which models were assessed, while the color corresponds to the domain transferred during recombination. (**a**) Relationship between the number of positively selected sites and the total number of toxins in the subtrees for which evolutionary selection was assessed, number of recombinants or parents (**b**), ω values (the nonsynonymous to synonymous substitution rate ratio, dN/dS) based on branch models within the foreground for all events (**c**) and those with a significant difference between branches (**d**). Figures (**e**) and (**f**) show the same estimates for background ω.

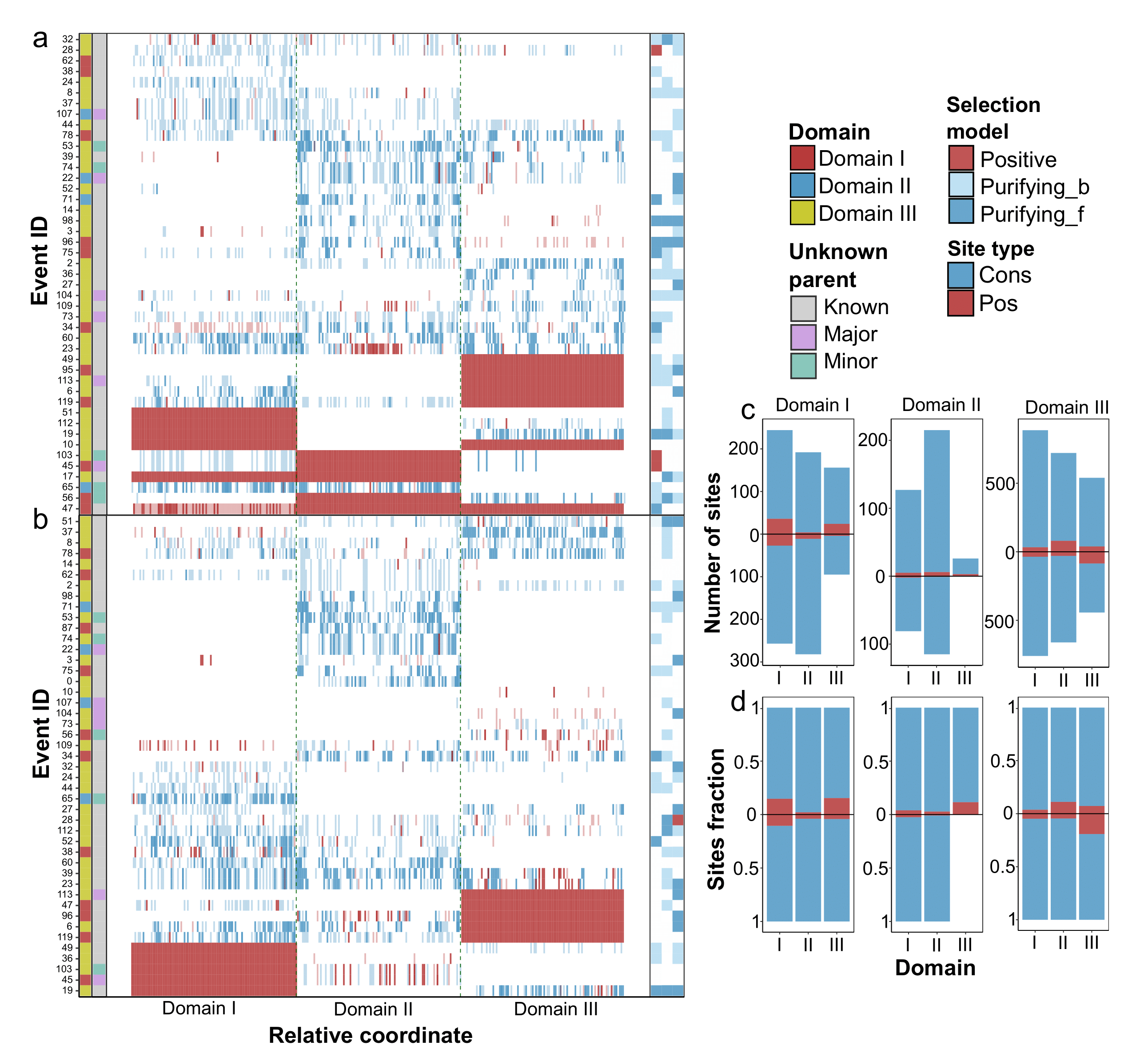

**Fig. S32**. Inferences of positively selected and conservative sites calculated on branch-site models in which recombinant and parental sequences were marked. Here and in other figures the selection pressure was determined on the sequences of different domains independently. (**a**) The heatmap sites with significant signals of evolutionary selection reported by best-fit branch-site models revealed by LRT (likelihood ratio test) in the cases when recombinants and parents (**b**) are marked. The left adjacent subpanels correspond to the domain that was transferred from the minor parent within domain exchanges and show whether a parent, either minor or major, is unknown, i.e., absent in the dataset analyzed. The site color represents if the site is positively selected, or conservative. The intensity of the color depends on the probability ranging from 0.95 to 0.99. the right subpanel indicates the significant model on branches with recombinants marked, namely, selection, and purifying selection either for the foreground or background only if the ω on branches is equal. (**c**) The number of positively selected and conservative sites determined using branch-site models among domains of recombinants and parents within the sets of events grouped according to the transferred domain and the fraction of these sites (**d**).

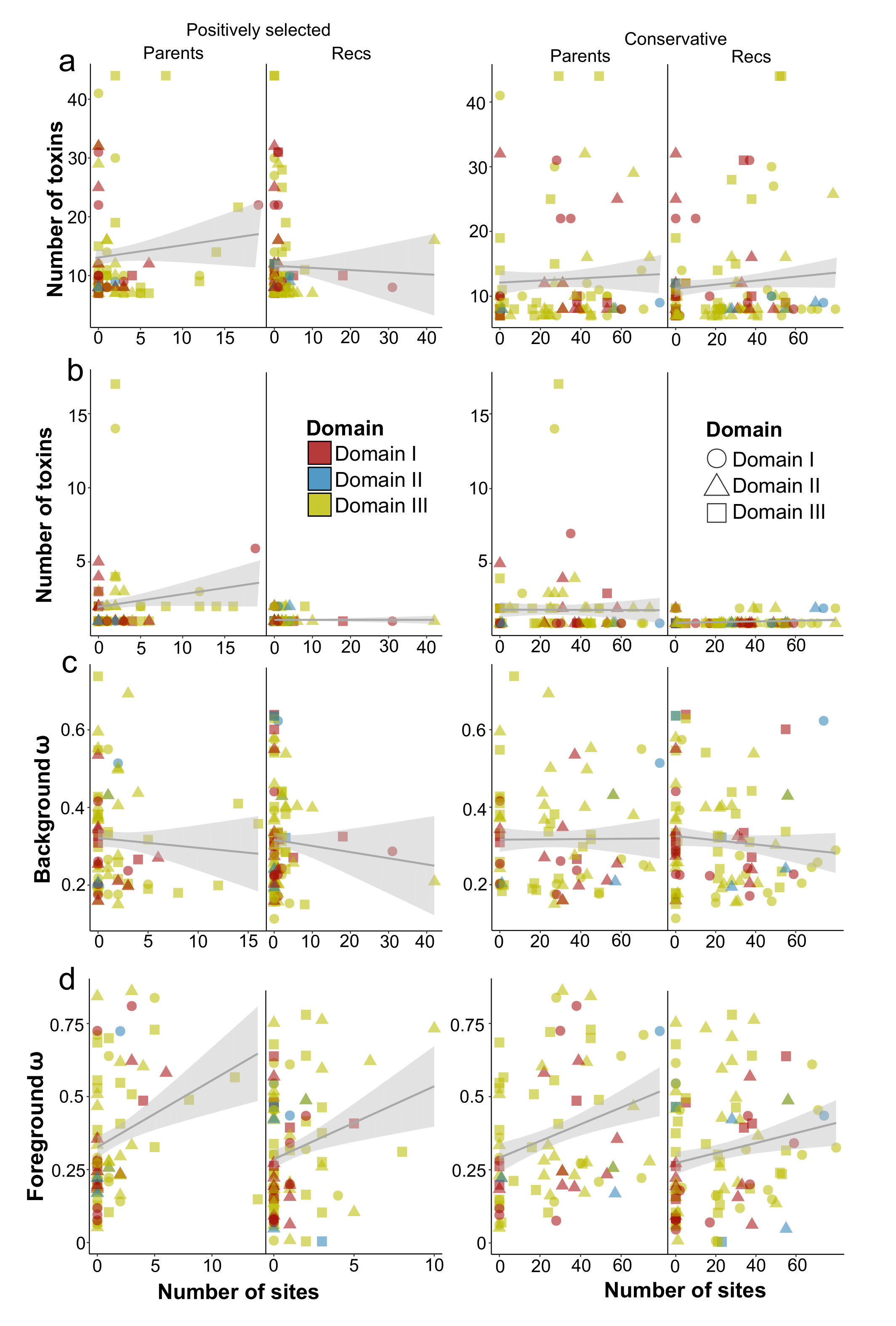

**Fig. S33**. Analysis of factors possibly delineating the number of positively selected sites according to evolutionary selection models calculated using branch-site models in which recombinants and parents were marked. Best-fit site models are determined by the LRT (likelihood ratio test). Here and in other figures the selection pressure was determined on the sequences of different domains independently. The shape of the points indicated the domains for which models were assessed, while the color corresponds to the domain transferred during recombination. (**a**) Relationship between the number of positively selected/conservative sites and the total number of toxins in the subtrees for which evolutionary selection was assessed, number of recombinants or parents (**b**), ω values (the nonsynonymous to synonymous substitution rate ratio, dN/dS) based on branch models within the background (**c**) and foreground in branches with marked recombinants and parents (**d**).

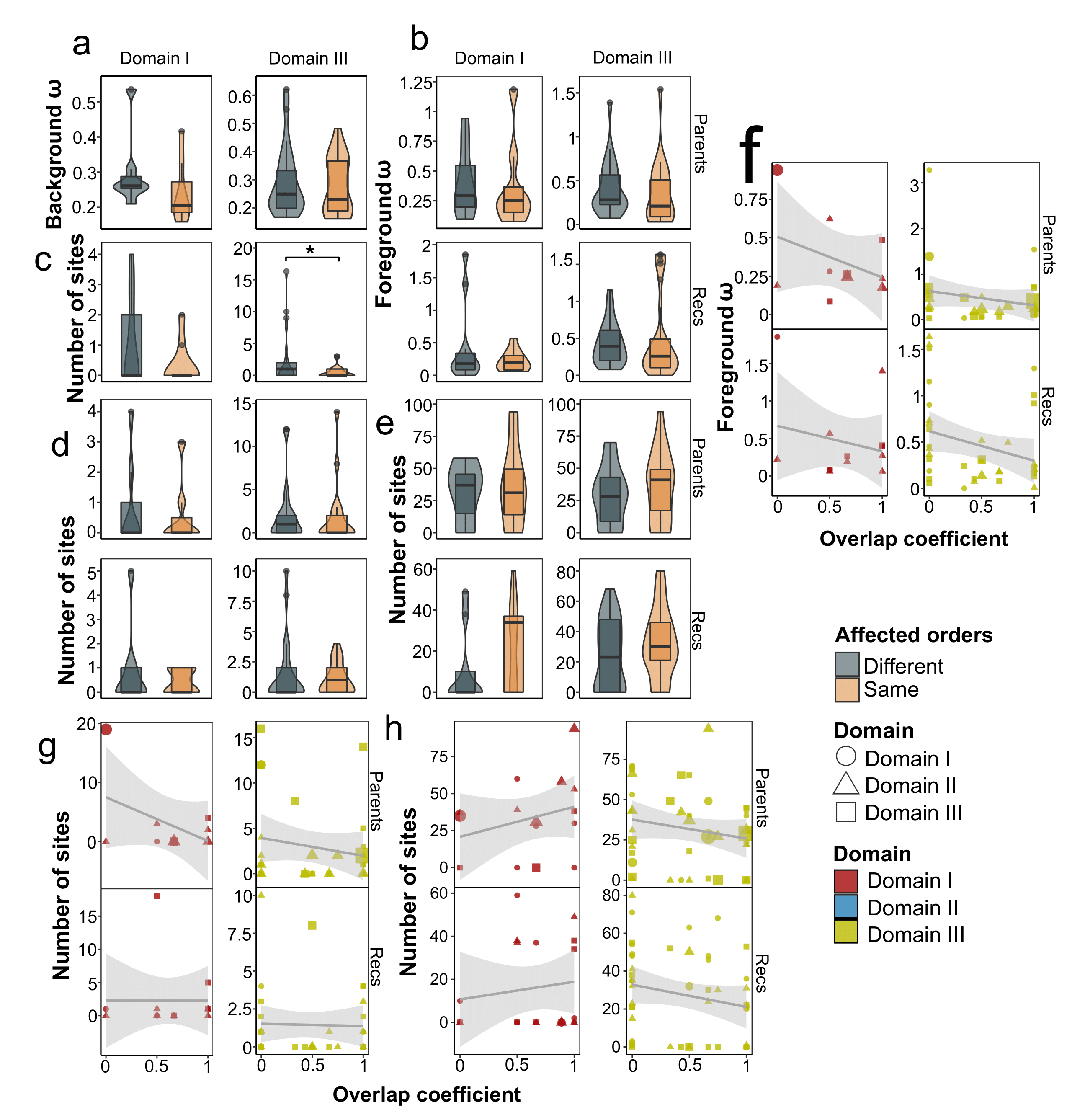

**Fig. S34**. Dependence between host specificity changes and features obtained when evaluating evolutionary selection. Best-fit branch, site, and branch-site models are determined by the LRT (likelihood ratio test). Here and in other figures the selection pressure was determined on the sequences of different domains independently. Relationship between changes in host orders (when at least one recombinant and one parent affect non-equal sets of orders) and ω values (the nonsynonymous to synonymous substitution rate ratio, dN/dS) based on branch models within the background (**a**) and foreground in branches with marked recombinants and parents (**b**), number of positively selected sites reported by site models (**c**), branch-site models (**d**), and the number of conservative sites (**e**). The relationship between the set of common species against which recombinants and parents are active and ω on foreground (**f**), and the number of positively selected (**g**) and conservative (**h**) sites revealed by branch-site models. The color indicates the domain transferred, while the shape of the dots corresponds to the domain for which the calculations were carried out.

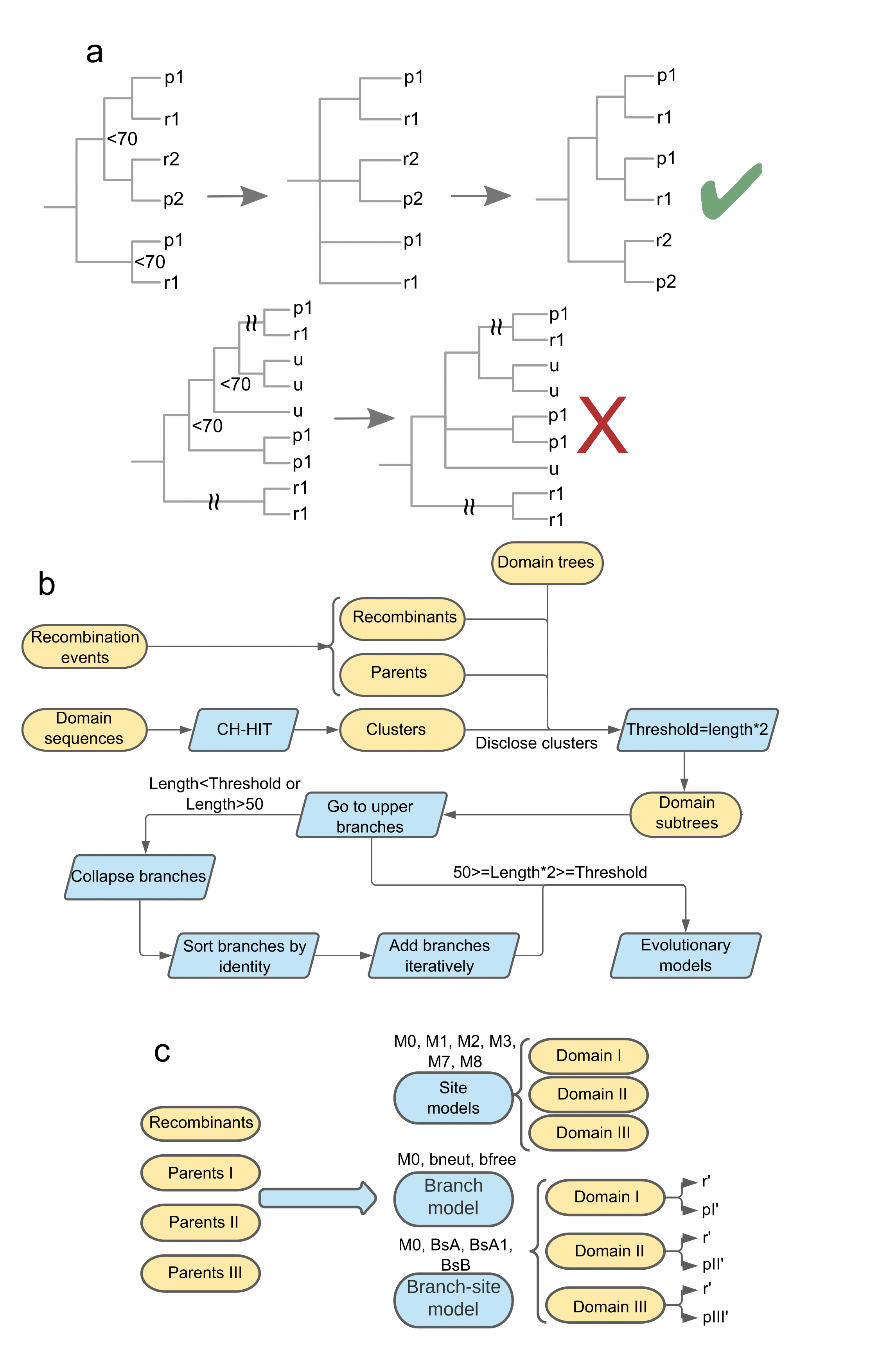

**Fig. S35**. Schemes of methodological approaches used in the study. (**a**) Filtering recombination events regarding the position of parents and recombinants on the phylogenetic tree. The analysis is performed for each domain-based tree independently, and the parent is determined by the type of transferred domain. In the first step, all nodes whose bootstrap supporting values are less than a predefined threshold of 70. In the next stage, we unified leaves with recombinants and parents in single branches by resolving polytomy. In case toxins missed by RPD4 predictions were found in the clades, the toxins were interpreted as parents of the individual domain according to the domain-wise tree. If the branches involving parents and recombinants were found in the root of the tree, their phylogeny was considered incongruent and the events were filtered out accordingly. Letters ‘r’, ‘p‘, and ‘u’ denote recombinants, parents, and unaffected toxins, respectively. (**b**) Data preparation scheme for evolutionary selection analysis. Domain-wise phylogenetic trees corrected for recombination as previously described were used to extract subtrees including the least common ancestor within the event. During the procedure, we disclosed clusters obtained with CD-HIT software. We then iteratively moved to upper branches and retained subtrees if their length exceeded the threshold defined as the number of recombination participants multiplied by two to get the background. If the procedure satisfied the threshold gained subtrees with more than 50 leaves, we screened all the subtrees with disclosed clusters arranged by the identity with the domain sequence of recombinants and iteratively added tree nodes until the required threshold was reached. The sequences in the trees were then aligned with MAFFT. Evolutionary models were selected using Modeltest-NG, and phylogenetic trees were reconstructed with RAxML-NG. The evolutionary selection was analyzed on the basis of the alignments and guiding trees using the ete3 package. (**c**) Shown are the evolutionary models tested. Data prepared during the previous steps were analyzed for each event separately. The best-fit models were chosen according to LRT (likelihood ratio test). Unlike site models, when examining branch and branch-site models, either recombinants or parents were marked as foreground branches and evaluated separately, which is displayed as apostrophes on the scheme.

**Supplementary text**

**Current databases contain toxins that can be included in the existing nomenclature**

To characterize the landscape of domain swaps between Cry toxins we first expanded the dataset through mining sequences bearing three functional domains and obtained 368 clusters accordingly with 110 of them sharing less than 95% homology with known sequences from the BPPRC database (fig. 1b). We should admit that in our aim to expand the existing nomenclature, however, it might be useful to consider some instances for reviewing and naming. Current criteria for consideration imply either >95% identity with the existing proteins or experimentally demonstrated activity against invertebrates [2]. While self-titled toxins found in genome assemblies such as cry13Ba1 [3] do not meet any of the criteria, some proteins like Cry3Aa12_81.9_AEZ53144.1 [4] were proved to form crystals and were toxic towards different hosts and worth consideration. The presence of Cry toxins in non-*Bt* species like *P. bifermentans* and *Paenibacillus popilliae* is a commonly known observation [5,6]. We, however, revealed the presence of these toxins in diverse species within the *Bacillus* genus, members of the Bacillales order, and, most intriguingly, *Clostridium* *botulinum* (fig. S3a), which corroborates the existing studies [7,8]. These observations allow us to assume the possible common origin of these toxins.

**Domain exchanges often accompany the evolution of bacterial toxins**

Current studies reported the SUKH superfamily of nuclease toxins combining C-terminal nuclease domains with N-terminal domains associated with secretion [9], *Clostridium botulinum* C2 toxin resembling C3 aside from NADbinding core [10], *Clostridioides difficile* TcdB toxin with multiple homologous recombination events generating new subtypes [11], mosaic LktA and LktB leukotoxins of *Mannheimia haemolytica* and *Pasteurella trehalosi* [12], and *Streptococcus salivarius* secrete glucosyltransferases [13]. Formation of hybrid toxins in prokaryotes, therefore, provides an adaptive advantage, hence the exchange of functional domains through recombination might be a common mechanism of evolution of pathogenicity determinants.

**Computational predictions of recombination events corroborate known examples of chimeric toxins**

The first *cry* gene was sequenced in 1985 [14]. Only a year later, when comparing sequences of crystal protein-encoding genes of *Bt* strains HD1 and HD73, Höfte et al. suggested that homologous recombination between genes occurs during the evolution of Cry toxins [15]. Lately, Almond and Dean conducted experiments to figure out the reasons behind the low diversity in domain composition of the Cry1Aa family [16]. After the construction of chimeric sequences from genes encoding Cry1Ac, Cry1Aa followed by transformation into *Bt*, intensive degradation of chimeric products by intracellular proteases was found, allowing us to propose that the respective exchanges lack evolutionary benefits [16]. These observations corroborate our results inasmuch as these toxins have not been detected in the same events mainly serving as donors of domains I and III for other proteins. When the *cryX* gene (later designated *cry9Ba*) was sequenced, Shevelev et al. noticed its partial similarity with *cry1G* localized downstream and suggested recombination events to play a role in the emergence of these toxins [17]. With more data available, our findings showed that Cry1Gb is a donor of domain III for toxin Cry9Ga underpinning evolutionary relationships between Cry1 and Cry9 families. Recently, an exchange of the domain I was reported for Cry2Aa17 receiving part of its sequence from Cry2Ab [18]. Our observations also included two events implying swaps of domain I within the Cry2 group, namely Cry2Am being a chimera of Cry2Al (source of domain I) and Cry2Ag together with Cry2Ab as well as Cry2Ai receiving the first domain from Cry2Ac and two remaining from Cry2Aa.

**Analyzing Cry toxins’ sequences**

To get the respective nucleotide sequences, a custom script implementing Biopython v1.73 [19] was utilized for identifying nucleotide accessions from the IPG annotations and downloading sequences accordingly, and the validity was checked through the comparison sequences translated via the Seqkit v0.10.1 utility [20] with the initial protein ones. The coordinates of the domains of Cry toxins’ clusters were obtained with CryProcessor v1.0, and the respective sequences were retrieved using the BEDTools v2.26.0 getfasta utility [21]. In order to exclude artificially modified toxins, we collected the sources of reference clusters based on all identical sequences from the IPG database. If the source of all entries belonged to the patents database, we further analyzed the patents’ descriptions and discarded the clusters that were described as artificially modified toxins.

After that, we summarized the lengths of the domains and their mean sequence similarities. To estimate sequence conservation, we calculated the highest per-site frequency of a non-gap symbol in the multiple alignments of toxins’ sequences with a custom Python script and plotted the distribution accordingly. We also analyzed the mean pairwise similarities of the sequences within a cluster. Cry toxins were compared with the known diversity deposited in the BPPRC by calculating identity percentages with the closest toxin from the database. We paid particular attention to the clusters containing 10 sequences or more by characterizing the spectrum of mutations classified as synonymous, non-synonymous, and frameshift mutations and plotting their distribution on the toxins’ sequences.

**Parameters for phylogeny reconstruction**

We aligned sequences of deduplicated reference clusters with the MAFFT aligner v7.429 [22] in the “localpair” mode with 1000 iterations and a gap penalty of 5. The alignments were trimmed with the trimAl v1.4.rev22 utility [23] with the following parameters: 30% minimum percentage of the positions in the original alignment to conserve, 0.3 minimum average similarity allowed, and the fraction of allowed gaps of 0.5 due to the high diversity of the Cry toxins. Best-fit evolutionary models were assessed by ModelTest-NG v0.1.7 [24] in the ‘ml’ mode according to the BIC value (Bayesian information criterion). Maximum likelihood (ML) trees were reconstructed using RAxML-NG v1.1.0 [25] with 1000 bootstrap replications. Four phylogenies were based on the sequences of each domain and a full toxin core with three domains partitioned according to the boundaries of the domains. We started with protein-based trees but later on preferred nucleotide ones due to better tree properties (lesser polytomy and higher bootstrap values).

**Detecting and characterizing recombination events**

Three-step filtration of the detected events was carried out. First, we excluded events caused by evolutionary processes rather than recombination marked in the RDP4 output. Second, we considered only the events spanning more than 70% of the domain, and partial events were analyzed separately. Third, we assessed the identity between the domains of parents and children and the phylogenetic congruence. Only the events in which the identity of the transmitted domain derived from a so-called minor parent was higher than the same domain from the major parent from which the other two domains were obtained. To examine phylogenetic congruence, we transformed phylogenies as follows: we used the single domain-based trees and collapsed branches with bootstrap support of less than 70% and grouped recombinants and their parents in a single subclade. Subsequently, we checked if parents and children formed a compact phylogenetic branch on a transformed tree devoid of toxins unrelated to the analyzed event. A detailed scheme of the filtering pipeline is presented in fig. S35a.

To access the quality of filtering we compared the identity difference between the transferred and non-transferred domains for initial and refined datasets and compared the obtained distributions with the Wilcox test corrected for multiple comparisons with the Benjamini-Hochberg (BH) procedure. Next, we summarized the number of mismatches between recombinants and the respective major and minor parents using the BioPython v1.73 library [19] in order to assess the evolutionary changes following the exchange events.

**Predicting the number of unknown toxins’ diversity**

To calculate the number of possible toxins representing unknown parents we first summarized the depths (d) of subtrees in a domain-wise way and obtained the sets of depths (D). The leaves with unknown events from collapsed trees were removed. We searched all subtrees with depth values in the respective trees and counted the number of nodes containing recombinants f(d) and revealed the number of leaves with parents p(d). In case unknown domains came from minor parents we used the respective trees, while for instances related to major parents, we considered adjacent domains. For each event with parents of unknown origin, we predicted the number of toxins as follows: U(d) = n(d)*p(d)/f(d). Finally, we summed up all predictions to get the total number of toxins potentially absent in the dataset.

**Assessing possible mechanisms of domain swaps**

Pangenome reconstruction was carried out with Panaroo v1.2.8 [26] with “--clean-mode strict” and “--remove-invalid-genes” options selected, and a 95% identity threshold for core genes with MAFFT aligner chosen for core genes extraction.

We collected the data on genomic coordinates for genes encoding Cry toxins and intersected them with the respective locations of mobile genetic elements (MGEs) using BEDTools v2.26.0 [21] getfasta. We further calculated the correlation between the number of different MGEs using the Pearson correlation coefficient and compared their abundance in assemblies’ groups with the BH-adjusted Wilcox test. The portions of *cry* genes localized in the MGEs were compared with Fisher’s exact test.

To perform clusterization of regions flanking breakpoints, three metrics were used, namely Euclidean, cosine, and Manhattan distances. The dissimilarity matrices were built based on vectors representing per-site identities, both raw and with roll mean procedure applied. Additionally, two clustering algorithms such as hierarchical clustering and k-means were tested. The best clustering pipeline was selected according to silhouette scores calculated with the factoextra v1.0.7 [27] package. We then plotted the distribution of per-site identity within clusters, manually picked peaks with high identity, and further compared mean identity values in these peaks with the overall similarity between transferred and adjacent domains using the Wilcox test.

**Revealing the impact of domain swaps on host specificity and toxins’ composition**

We first compared the sets of affected hosts of recombinants and parents using data for different ranks and strains and examined changes in orders treating events as different if parents and recombinants had at least one uncommon order. To study the effect of recombination on the composition of toxins, we compared the sets of toxins presented in the assemblies/strains comprising recombinants and parents using the Jaccard coefficient (the number of intersections divided by the length of the union). We also summed up the number of toxins in the respective isolates as well as in those devoid of recombination participants and compared them by applying the Wilcox test adjusted for multiple comparisons. We also performed similar comparisons in terms of the number of affected hosts and species. To exclude the possible impact of sets’ size, we utilized Szymkiewicz–Simpson metrics (the number of intersections divided by minimum length). Linear models were used to find factors (number of toxins, identity between domains, parent type, transferred domain) possibly delineating the intersection between the sets of hosts, and binomial regression to study the impact of factors on changes in the affected orders.

**Preparing the data for evolutionary selection analysis**

Before obtaining guiding trees and alignments, we discarded 100% duplicates for sequences of the domains within the reference clusters. We then analyzed each of the domains separately. To obtain the background when dealing with recombination events we set a threshold determining how many sequences should be included, no less than twice as many toxins as a sum of recombinants and parents pertaining to a particular domain. We utilized the ete3 v.3.1.1 package [28] to get a subtree containing the common ancestor of recombinants and parents and iteratively moved from the subtree to the upper nodes to get a set of sequences satisfying the threshold. During the procedure, we disclosed CD-HIT-based clusters picking no more than 5 random toxins for the cluster in case it incorporated more sequences. If the set satisfying the threshold exceeded 50 representatives, we collapsed nodes marked by bootstrap support of less than 70%, arranged nodes according to the similarity of protein sequences to the respective recombinant sequences, and iteratively added nodes until the length threshold reached the required threshold.
